# Chromosome-level scaffolding of haplotype-resolved assemblies using Hi-C data without reference genomes

**DOI:** 10.1101/2023.11.18.567668

**Authors:** Xiaofei Zeng, Zili Yi, Xingtan Zhang, Yuhui Du, Yu Li, Zhiqing Zhou, Sijie Chen, Huijie Zhao, Sai Yang, Yibin Wang, Guoan Chen

## Abstract

Scaffolding is crucial for constructing most chromosome-level genomes. The high-throughput chromatin conformation capture (Hi-C) technology has become the primary scaffolding strategy due to its convenience and cost-effectiveness. As sequencing technologies and assembly algorithms advance, constructing haplotype-resolved genomes is increasingly preferred because haplotypes can provide additional genetic information on allelic and non-allelic variations. ALLHiC is a widely used allele-aware scaffolding tool designed for this purpose. However, its dependence on chromosome-level reference genomes and a higher chromosome misassignment rate still impede the unraveling of haplotype-resolved genomes. In this paper, we present HapHiC, a reference-independent allele-aware scaffolding tool with superior performance on chromosome assignment as well as contig ordering and orientation. Additionally, we provide new insights into the challenges in allele-aware scaffolding by conducting comprehensive analyses on various adverse factors. Finally, with the help of HapHiC, we constructed the haplotype-resolved allotriploid genome for *Miscanthus* × *giganteus*, an important lignocellulosic bioenergy crop. HapHiC is available at https://github.com/zengxiaofei/HapHiC.

## Introduction

The construction of a high-quality reference genome serves as the basis for functional genomics research in a species. Chromosome scaffolding is a necessary step in *de novo* building eukaryotic chromosome-level genomes, except for directly assembling telomere-to-telomere (T2T) genomes^1^. Its objective is to determine the chromosome assignment of contigs or scaffolds in the assemblies, as well as the ordering and orientation of these sequences on the chromosomes. In early genome research, chromosome scaffolding was often achieved using the information from linkage groups and genetic distance in genetic maps^2^. However, in recent years, the high-throughput chromatin conformation capture (Hi-C) technology has gradually replaced genetic maps due to its simplicity, short cycle, and low cost, making it the most widely used chromosome scaffolding solution^3–8^. Hi-C links are generated by proximity ligation and massively parallel sequencing to indicate the frequency of chromatin interactions between different loci in the genome^9^. This information can be used to infer chromosome territories, as well as the distance and orientation between contigs or scaffolds^3^. Several Hi-C-based scaffolding tools, including LACHESIS^3^, HiRise^4^, 3D-DNA^5^, SALSA2^6^, and YaHS^8^, have been developed for haploid and haplotype-collapsed assemblies.

For heterozygous diploids or polyploids, a haplotype-resolved assembly consists of two or more sets of haploid sequences. In contrast to a haplotype-collapsed assembly, it provides additional genetic information, such as bi- or multi-alleles, and *cis*/*trans* configurations among non-allelic variations^10^. Recent advances in sequencing technologies and assembly algorithms have propelled the unraveling of haplotype-resolved genomes. HiFi sequencing from Pacific Biosciences (PacBio) and duplex sequencing from Oxford Nanopore Technologies (ONT) have both achieved a base accuracy level of Q30 (99.9%), which provides a solid foundation for more accurate phasing of alleles. Trio binning uses short reads from parental genomes to phase long reads, enabling phasing at the whole-genome level^11^. More recently, hifiasm takes advantage of Hi-C sequencing data for chromosome-level phasing without parental data^12^. These two methods have demonstrated high accuracy in dealing with diploid or diploid-like allopolyploid genomes. Consequently, subsequent chromosome scaffolding can be independently performed on each phased haplotype.

Autopolyploidy is prevalent in seed plants, especially in economically important crops^13^. Haplotype phasing in autopolyploid genomes facilitates the study of crop domestication history and genetic breeding^14^. It also lays the foundation for analyzing allele expression dominance and genome evolution after whole-genome duplication (WGD)^10^. However, assembling haplotype-resolved autopolyploid genomes is more challenging than diploid genomes. Trio binning is unsuitable for autopolyploids^11^, and the Hi-C-based algorithm in hifiasm produces unbalanced phasing results during the assembly of autopolyploid genomes^15^. Therefore, the most common strategy for constructing a haplotype-resolved autopolyploid genome is to perform allele-aware scaffolding, which utilizes Hi-C data to allocate contigs to different haplotypes simultaneously during chromosome scaffolding^7^. On the other hand, scaffolding each phased haplotype separately may result in errors because the Hi-C data from multiple haplotypes are aligned to a single haplotype, disregarding possible chromosomal structural variations between haplotypes. This once again emphasizes the importance of allele-aware scaffolding.

ALLHiC is a widely used Hi-C scaffolding tool specifically designed for allele-aware scaffolding^7^. It effectively identifies allelic sequences and removes the Hi-C links between them to reduce interference prior to clustering. ALLHiC has demonstrated robust performance in haplotype phasing and has been applied to resolve the diploid and autotetraploid genomes of several important crops^16–20^. However, this method requires a chromosome-level reference genome from a closely related species, which may not be available for many species. Although it is feasible to assemble and annotate a haplotype-collapsed genome as the reference^19^, it significantly increases the time and cost of genome research. Additionally, ALLHiC has been observed to introduce clustering errors when using the reference genome (as discussed in Results). These limitations and drawbacks have hindered the construction of haplotype-resolved genomes to some extent, especially in autopolyploids.

In this study, we introduce HapHiC, a Hi-C-based scaffolding tool that enables allele-aware chromosome scaffolding of autopolyploid assemblies without reference genomes. We conducted a comprehensive investigation into the factors that may impede the allele-aware scaffolding of genomes. Compared to existing methods, HapHiC demonstrated a higher scaffolding contiguity and lower misassignment rates when addressing these challenges. Additionally, HapHiC is fast, resource-efficient, and has been successfully validated in genomes with varying ploidies and taxa. By utilizing HapHiC, we finally constructed the haplotype-resolved genome of *Miscanthus* × *giganteus*, an important lignocellulosic bioenergy crop.

## Results

### Overview of HapHiC

To ensure concision and clarity, we will use the term “contigs” to refer to both contigs and scaffolds input into scaffolding tools. Assembly errors in phased assemblies and strong Hi-C signals between allelic contigs are considered the main obstacles that hinder allele-aware scaffolding. HapHiC addresses these challenges through two strategies. First, HapHiC prioritizes the chromosome assignment of contigs (**Fig. 1c,d**) before determining their ordering and orientation (**Fig. 1e,f**), similar to the approach used by LACHESIS^3^ and ALLHiC^7^. This is because determining scaffold or chromosome boundaries during or after contig ordering and orientation, as done by 3D-DNA^5^, SALSA2^6^, and YaHS^8^, can amplify the negative effects of assembly errors and undesirable inter-allele Hi-C signals on chromosome assignment by disrupting the ordering and orientation of contigs. Therefore, HapHiC employs this “divide-and-conquer” strategy to isolate their negative impacts between these two steps. Second, HapHiC applies several preprocessing steps before chromosome assignment to correct and filter input contigs and remove Hi-C links between allelic contigs (**Fig. 1b**). Specifically, HapHiC implements an efficient and stringent method to correct chimeric contigs with minimal impact on contig N50. Subsequently, low-information contigs, such as short contigs and those lacking Hi-C links, are temporarily removed before clustering due to their propensity for errors. In addition to the conventional method of identifying collapsed contigs based on Hi-C link density, we introduce a unified “rank-sum” algorithm to further filter out residual chimeric and collapsed contigs. Moreover, undesirable inter-allele Hi-C links are removed based on the distribution pattern of Hi-C links. These innovative approaches significantly enhance HapHiC’s tolerance to assembly errors and enable its capability of allele-aware scaffolding without reliance on reference genomes.

**Fig. 1.**
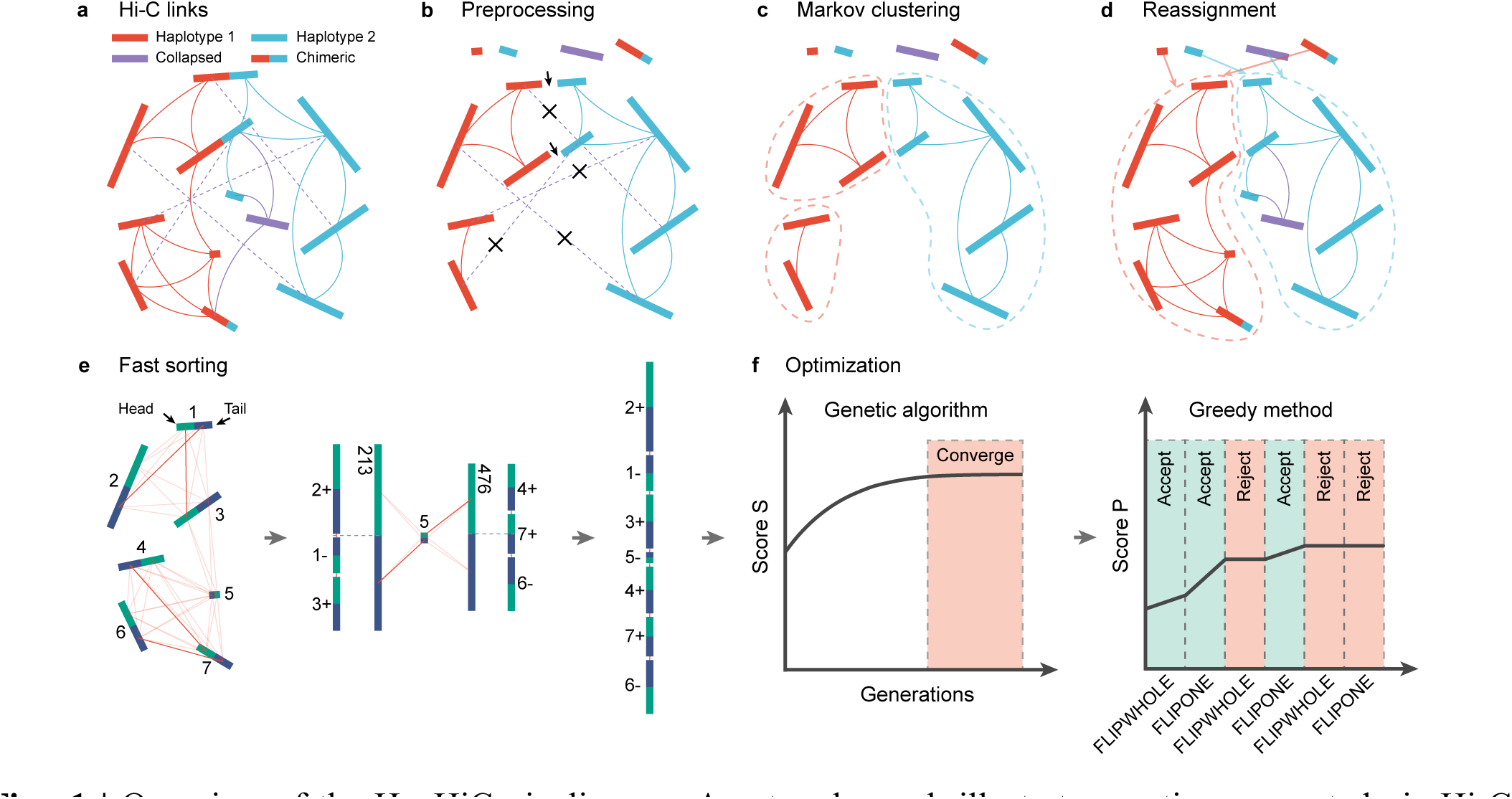
| Overview of the HapHiC pipeline. **a**, A network graph illustrates contigs connected via Hi-C links. Contigs from haplotype 1 and haplotype 2 are represented by red and blue rectangles, respectively. Collapsed and chimeric contigs are shown as purple and bicolor rectangles, respectively. Hi-C links within haplotype 1, within haplotype 2, and connecting collapsed contigs are depicted as red, blue, and purple curves, respectively. Inter-allele Hi-C links are represented by dashed purple lines. **b**, The preprocessing step involves assembly correction, filtering out low-information contigs, discarding collapsed and chimeric contigs, and removing inter-allele Hi-C links. Breakpoints of assembly correction are represented by black arrows, while crosses indicate the removal of inter-allele Hi-C links. **c**, Preliminary Markov clustering is performed with the remaining contigs and Hi-C links. **d**, The reassignment step rescues and reassigns contigs to the most suitable clusters and performs an additional agglomerative hierarchical clustering if the number of clusters exceeds the expected number of chromosomes. **e**, An efficiency-improved 3D-DNA iterative algorithm is used for contig ordering and orientation, referred to as “fast sorting”. In each round of iteration, a confidence graph is constructed using the hemi-parts (green and blue segments) of contigs or scaffolds. The graph is then filtered to retain only reciprocal best matching (opaque red lines). Unlike the original 3D-DNA algorithm, the hemi-parts of scaffolds are split at the approximate midpoints to eliminate the need for reconstructing graph from scratch. **f**, Optimization of contig ordering and orientation is based on the result of fast sorting using the genetic algorithm and greedy method.

The remaining contigs and Hi-C links are then used to construct a contact matrix. HapHiC performs preliminary contig clustering using a Markov cluster algorithm^21^ (MCL) and selects the optimal clustering result through automatic parameter tuning (**Fig. 1c**). In the subsequent reassignment step, the filtered-out and potential misassigned contigs are respectively rescued and reassigned to the most suitable clusters (**Fig. 1d**). If the number of clusters exceeds the expected number of chromosomes, HapHiC carries out additional agglomerative hierarchical clustering^22^ to group them into chromosome-level clusters. After chromosome assignment, HapHiC conducts contig ordering and orientation by integrating the algorithms from 3D-DNA^5^ and ALLHiC^7^. Initially, the contigs in each cluster are ordered and oriented using an efficiency-improved 3D-DNA algorithm, which we refer to as “fast sorting” (**Fig. 1e**). The results are further optimized by employing the genetic algorithm in ALLHiC to generate the final chromosome-level pseudomolecules (**Fig. 1f**). This integration enhances the accuracy of contig ordering and orientation while significantly reducing the execution time and the number of iterations compared to ALLHiC.

### Factors that may impede the allele-aware scaffolding of phased assemblies

In this section, we evaluated the negative impact of various factors on allele-aware scaffolding of phased assemblies and compares the performance of HapHiC with other mainstream Hi-C-based scaffolding tools, including ALLHiC^7^, LACHESIS^3^, 3D-DNA^5^, SALSA2^6^, and YaHS^8^. We used the haplotype-resolved autotetraploid genome of *Medicago sativa* XinJiangDaYe^17^ to establish a ground truth (**Supplementary Figs. 1** and **2**) and generated a series of fragmented assemblies by simulating multiple adverse factors of varying degrees (**Supplementary Fig. 3** and **Supplementary Tables 1-10**).

Contig contiguity is a crucial factor affecting allele-aware scaffolding. Phased assemblies typically have lower contig contiguity compared to collapsed assemblies. When the contig N50 decreased from 2 Mb to 25 Kb (**Supplementary Fig. 4**), all Hi-C scaffolding tools examined experienced a decline in final scaffold contiguity, anchoring rate, and an increase in misassignment rate between homologous chromosomes (**Fig. 2l**, **Supplementary Fig. 5**, and **Supplementary Data 1**). Among these tools, HapHiC consistently showed the highest scaffold contiguity and extremely low misassignment rates. In contrast, 3D-DNA and SALSA2 tended to produce highly fragmented scaffolds, with the scaffold contiguity values mostly less than 0.5. ALLHiC, LACHESIS, and YaHS exhibited much higher scaffold contiguity than 3D-DNA and SALSA2, but their misassignment rates were also elevated. Furthermore, when the contig N50 dropped below 100 Kb, the memory usage of YaHS became too high to scaffold the assemblies. Additionally, we assessed the impact of contig length distribution on allele-aware scaffolding, with the coefficients of variation (CVs) of the contig length ranging from 0.2 to 3 (**Supplementary Fig. 6**). All scaffolding tools performed stably in this regard, with HapHiC remaining the most outstanding among them (**Fig. 2l**, **Supplementary Fig. 7**, and **Supplementary Data 1**).

**Fig. 2.**
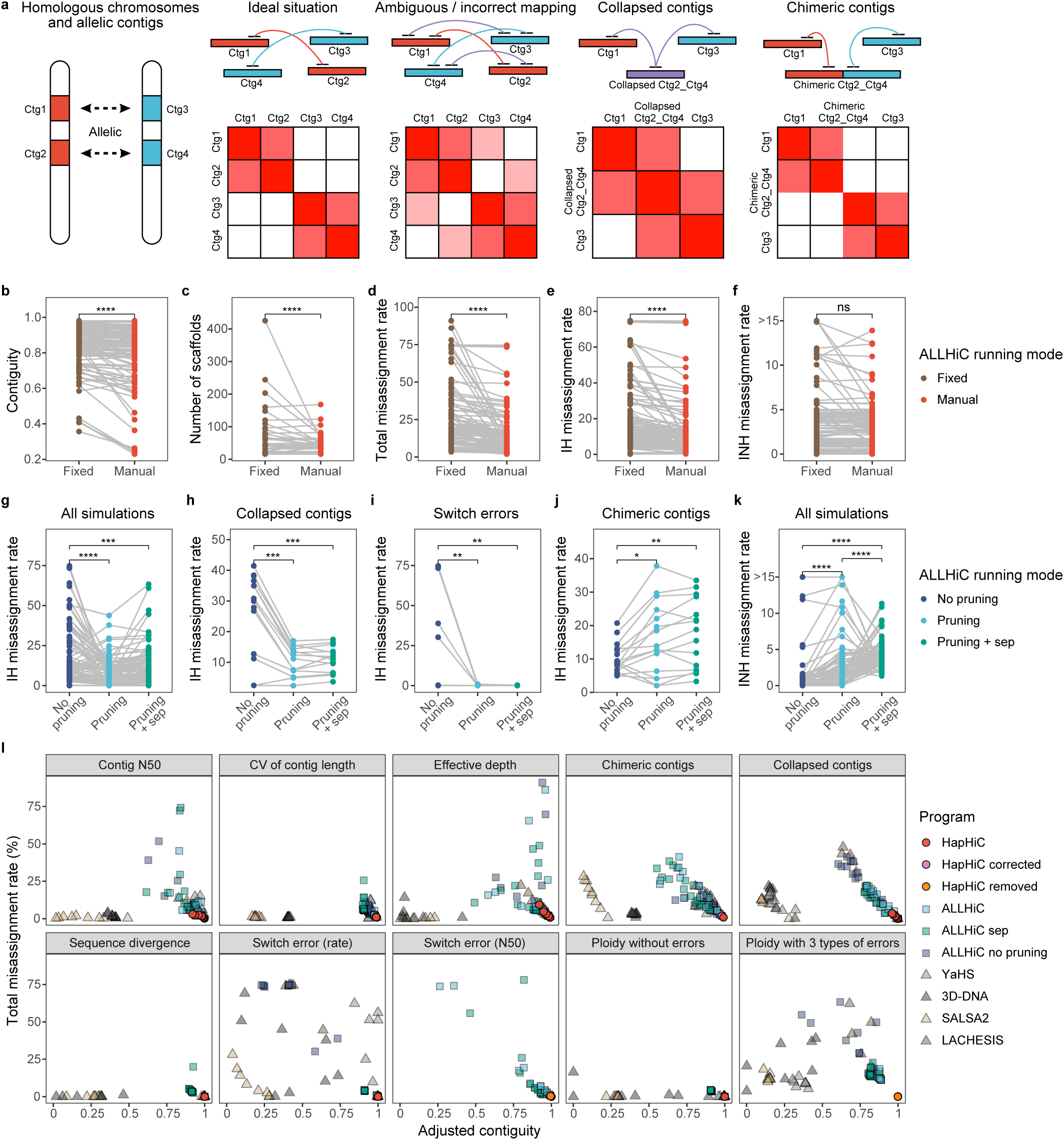
| Comprehensive performance analysis of Hi-C-based scaffolding tools in chromosome assignment under various adverse conditions. **a**, A schematic diagram illustrating the potential challenges in allele-aware scaffolding, including the presence of collapsed contigs, chimeric contigs, and ambiguous or incorrect mapping. **b**-**f**, The effect of manually tuning parameter *k* on ALLHiC performance (*n* = 141). **g**-**k**, The effect of pruning and separating homologous groups on ALLHiC performance (*n* = 152, 14, 14, 16, and 152 respectively). **i**, Performance analysis of Hi-C-based scaffolding tools on assemblies with various adverse factors of varying degrees. HapHiC was executed in default mode (HapHiC), with assembly corrected (HapHiC corrected) or with inter-allele Hi-C links removed (HapHiC removed). The total misassignment rate includes misassignment rate between both homologous and non-homologous chromosomes. The adjusted contiguity is calculated by multiplying the contiguity by the anchoring rate. *P* values were derived from two-sided Wilcoxon signed-rank tests.

As the cost of next-generation sequencing decreases, Hi-C sequencing depth seems to no longer be a limiting factor in scaffolding. However, in the context of allele-aware scaffolding of phased assemblies, Hi-C reads can be aligned to multiple allelic loci on homologous chromosomes simultaneously. These reads are often filtered out due to low mapping quality, resulting in a reduction of effective data. To demonstrate this, we simulated effective sequencing depths ranging from 11X to 0.02X (**Supplementary Fig. 8**). The results showed that the anchoring rate of 3D-DNA and SALSA2 declined rapidly with decreasing sequencing depth (**Fig. 2l**, **Supplementary Fig. 9**, and **Supplementary Data 1**). Notably, YaHS and SALSA2 failed to scaffold at depths below 1X and 0.05X, respectively. For other scaffolding tools, scaffold contiguity decreased and misassignment rates increased when the effective Hi-C data dropped below 0.05X. HapHiC performed well even at extremely low depths, exhibiting the highest scaffold contiguity and relatively low misassignment rates.

Chimeric contigs, a common assembly error in phased assemblies, result from misjoins between nonadjacent sequences (**Fig. 2a**). These misjoins can occur between homologous or non-homologous chromosomes (**Supplementary Fig. 10**), leading to chromosome misassignments during scaffolding. ALLHiC has been reported to be highly susceptible to chimeric contigs without assembly correction^7^. We evaluated the accuracy and precision of assembly correction for each tool in dealing with chimeric contigs of different lengths (**Supplementary Figs. 11-14**). When the length was below 800 Kb and 100 Kb, respectively, neither SALSA2 nor ALLHiC could break any chimeric contigs (**Supplementary Fig. 12**). Moreover, 3D-DNA, SALSA2, and ALLHiC tended to break too many non-chimeric contigs, leading to a significant decrease in contig contiguity (**Supplementary Fig. 13**). HapHiC achieved 84.2% to 94.9% of the true positive rate (TPR) of YaHS in identifying chimeric contigs, while maintaining a false positive rate (FPR) that was 0.4 to 7.1 times lower than that of YaHS. This suggests that HapHiC has comparable sensitivity to YaHS in identifying chimeric contigs, but with significantly higher stringency. Additionally, HapHiC consistently demonstrated exceptional precision in determining breakpoints, as evidenced by the highest proportions of breakpoints within 10 Kb of simulated misjoins, ranging from 81.0% to 95.7% across varying contig lengths (**Supplementary Fig. 14**).

With assembly error correction, some chimeric contigs may still slip through the net. To address this, we introduce a “rank-sum” algorithm in HapHiC for further contig filtering (**Supplementary Fig. 15**). This algorithm is based on the following principles: (1) the one- or three-dimensional neighborhoods of a specific genome region should also be neighborhoods to each other, and (2) this is not applicable to chimeric contigs, which may be misjoined from different regions of the chromosome or even different chromosomes. Thus, potential chimeric contigs can be identified by measuring the density of their respective neighborhoods (**Supplementary Fig. 16**). The receiver operating characteristics (ROC) curve demonstrates the superior performance of the rank-sum algorithm in identifying chimeric contigs, whether formed between homologous chromosomes or non-homologous chromosomes (**Supplementary Fig. 17**). Although the algorithm is not sensitive to chimeric contigs formed within the chromosome, this type of error does not adversely affect chromosome assignment. Using this algorithm alone, HapHiC can tolerate up to 20% chimeric contigs (**Fig.2l**, **Supplementary Fig. 18**, and **Supplementary Data 1**). With assembly correction enabled, HapHiC can accurately assign contigs into chromosomes even when up to 40% of contigs are chimeric. In contrast, other scaffolding tools exhibited significantly higher misassignment rates between homologous chromosomes. We also found that when the proportion of chimeric contigs was below 25%, the performance of ALLHiC with assembly correction was even less effective than without any correction (**Supplementary Fig. 19**). This could be explained by a hypothesis that the negative impact of ALLHiC correction on contig contiguity is more severe compared to a low proportion of chimeric contigs.

Another type of assembly error that can lead to misassignments between homologous chromosomes is collapsed contigs. These contigs are consensus sequences collapsed from highly similar allelic regions (**Fig. 2a** and **Supplementary Fig. 20**). LACHESIS^3^ and ALLHiC^7^ simply identify and filter out collapsed contigs based on Hi-C link density. However, there are two potential issues that can adversely affect the performance of this method (**Supplementary Fig. 21**). First, in phased autopolyploid assemblies, collapse frequently occurs and can involve more than two haplotypes, resulting in a higher average or median Hi-C link density. Using a fixed cutoff for classification is inefficient in such cases. Second, both scaffolding tools neglect intra-contig links when calculating Hi-C link density, leading to bias against contig length. Similar to chimeric contigs, the neighborhoods of collapsed contigs are expected to exhibit a lower density compared to normal contigs. As anticipated, the rank-sum algorithm has proven to be a unified approach that is also effective for identifying collapsed contigs (**Supplementary Fig. 22**). Additionally, the two methods showed complementary trends in relation to the number of collapsed haplotypes (**Supplementary Fig. 23**). Specifically, the link density method exhibited higher sensitivity to four-haplotype collapsed contigs, while the rank-sum algorithm demonstrated greater efficiency in identifying two-haplotype collapsed contigs. Consequently, we integrated these two methods in the preprocessing step of HapHiC. This integration enabled HapHiC to tolerate up to 25% of collapsed contigs in chromosome assignment, significantly surpassing other examined Hi-C scaffolding tools (**Fig. 2l**, **Supplementary Fig. 24**, and **Supplementary Data 1**). In contrast, the pruning process of ALLHiC only partially mitigated the adverse effects of collapsed contigs.

One commonly held perspective is that low sequence divergence between haplotypes can hinder allele-aware scaffolding by causing incorrect mapping of Hi-C reads (**Fig. 2a**). In real cases, strong signals of inter-allele Hi-C links are often observed to be diagonally distributed between homologous chromosomes (**Supplementary Fig. 25**). However, our simulation tests yielded contradictory results. The relative proportion of inter- and intra-homologous chromosome Hi-C links did not change significantly with sequence divergence after filtering with mapping quality and edit distance (**Supplementary Figs. 26** and **27**). Furthermore, most Hi-C scaffolding tools performed well even when the sequence divergence between haplotypes was as low as 0.1% (**Fig. 2l**, **Supplementary Fig. 28**, and **Supplementary Data 1**).

To address this contradiction, we constructed two phased assemblies using the same PacBio HiFi reads but with different genome assemblers, hifiasm^12^ and HiCanu^23^. We observed stronger signals of inter-allele Hi-C links in the HiCanu assembly (**Supplementary Fig. 29b**) compared to the hifiasm assembly (**Supplementary Fig. 29a**). This suggests that the presence of unfavorable inter-allele Hi-C signals is more likely caused by a type of assembly error. Based on their distribution patterns on the Hi-C contact maps (**Supplementary Figs. 29** and **30**), it is evident that these errors are not due to large-scale chimeric or collapsed sequences. Instead, we hypothesize that they could be switch errors at the base level. To verify this hypothesis, we simulated switch errors by randomly shuffling single nucleotide polymorphisms (SNPs) and small insertions/deletions (InDels) between haplotypes. As a result, we were able to reproduce similar inter-allele Hi-C signals along the diagonal (**Supplementary Fig. 31**), which became stronger as the switch error rate increased (**Supplementary Fig. 32**). Our findings indicate that incorrect mapping of Hi-C links is introduced by switch errors rather than the inherent sequence divergence between haplotypes.

ALLHiC identifies allelic contigs by examining gene synteny between the assembly and an annotated, chromosome-level reference genome from the same or a closely related species. During the pruning process, Hi-C links between allelic contigs are removed. However, such a reference genome is not always available for all species. As a result, we developed a reference-free method in HapHiC that relies on the distribution pattern of Hi-C links (**Supplementary Figs. 33** and **34**). In simulation tests, our reference-free method allowed HapHiC to tolerate a switch error rate of up to 25% (**Fig. 2l**, **Supplementary Fig. 35**, and **Supplementary Data 1**) and exhibited higher efficiency than ALLHiC in identifying allelic contigs of low contiguity (**Fig. 2l**, **Supplementary Figs. 36** and **37**). In contrast, scaffolding tools that are not allele-aware or executed without removing inter-allele Hi-C links were severely disrupted when the switch error rate exceeded 5% (**Supplementary Fig. 35**).

While tetraploids constitute the majority of published autopolyploid genomes, species with higher ploidies are prevalent in both wild and cultivated plants. We assessed the impact of genome ploidy on allele-aware scaffolding. In the absence of assembly errors, all Hi-C scaffolding tools, except for 3D-DNA, demonstrated stable performance when handling ploidies ranging from 1 to 16 (**Fig. 2l**, **Supplementary Fig. 38**, and **Supplementary Data 1**). However, when we introduced 5% each of chimeric, collapsed contigs, and switch errors in simulated assemblies of various ploidies, only HapHiC consistently produced perfect chromosome assignment results (**Fig. 2l**, **Supplementary Fig. 39**, and **Supplementary Data 1**). Additionally, the performance of the reassignment process in HapHiC was validated through separate tests (**Supplementary Fig. 40**).

Our results confirm that the reference-dependent pruning method in ALLHiC effectively and robustly reduces misassignments between homologous chromosomes, particularly when dealing with collapsed contigs and switch errors (**Fig. 2g-i**). However, the main concern is that the reference genome may not be available or may require significant effort and cost for construction and annotation. Additionally, using reference genomes is a double-edged sword that can increase misassignments between non-homologous chromosomes and exacerbate the adverse effect of chimeric contigs (**Fig. 2j,k**). Furthermore, substantial parameter tuning is often necessary for ALLHiC to achieve improved chromosome assignment results (**Fig. 2b-f**). HapHiC has addressed these problems, demonstrating stronger tolerance to various assembly errors and unfavorable factors (**Fig. 2l** and **Supplementary Data 1**). These improvements enhance its adaptability and capability in tackling more intricate allele-aware scaffolding problems.

#### Accuracy of contig ordering and orientation

After chromosome assignment, the contig ordering and orientation of a phased assembly become similar to those of an unphased assembly. To evaluate the accuracy of each Hi-C-based scaffolding tool in contig ordering and orientation, we simulated genome assemblies of rice^24^ (*Oryza sativa*), *Arabidopsis*^25^ (*Arabidopsis thaliana*), and human^1^ (*Homo sapiens*) with varying contig N50 values (**Supplementary Table 11**).

Initially, the performance of HapHiC and ALLHiC was compared using the built-in scoring system of ALLHiC^7^ (**Supplementary Fig. 41** and **Supplementary Data 2**). Even when only fast sorting was employed, the initial scores of HapHiC were already comparable to or even higher than the final scores achieved by ALLHiC. In assemblies with low contig contiguity, these scores were further improved during the subsequent optimization process, resulting in a significant reduction in the number of iterations required for the genetic algorithm to converge.

Subsequently, we introduced two objective metrics to assess the accuracy of contig ordering and orientation for all Hi-C-based scaffolding tools (**Fig. 3a**). The first metric, Lin’s concordance correlation coefficient^26^ (CCC), measures the consistency between the results and the reference chromosomes on a large scale. The second metric is the “cost” calculated by DERANGE II^27^, which approximates the number of moves required to adjust the results for complete consistency with the reference chromosomes via transposition and inversion. This cost can also represent the number of steps needed to achieve optimal results in Juicebox^28^. As the cost is independent of contig length, it is suitable for quantifying the results on a smaller scale. By employing these two metrics, we can categorize the results into four quadrants based on their distinct tendencies (**Fig. 3a**).

**Fig. 3.**
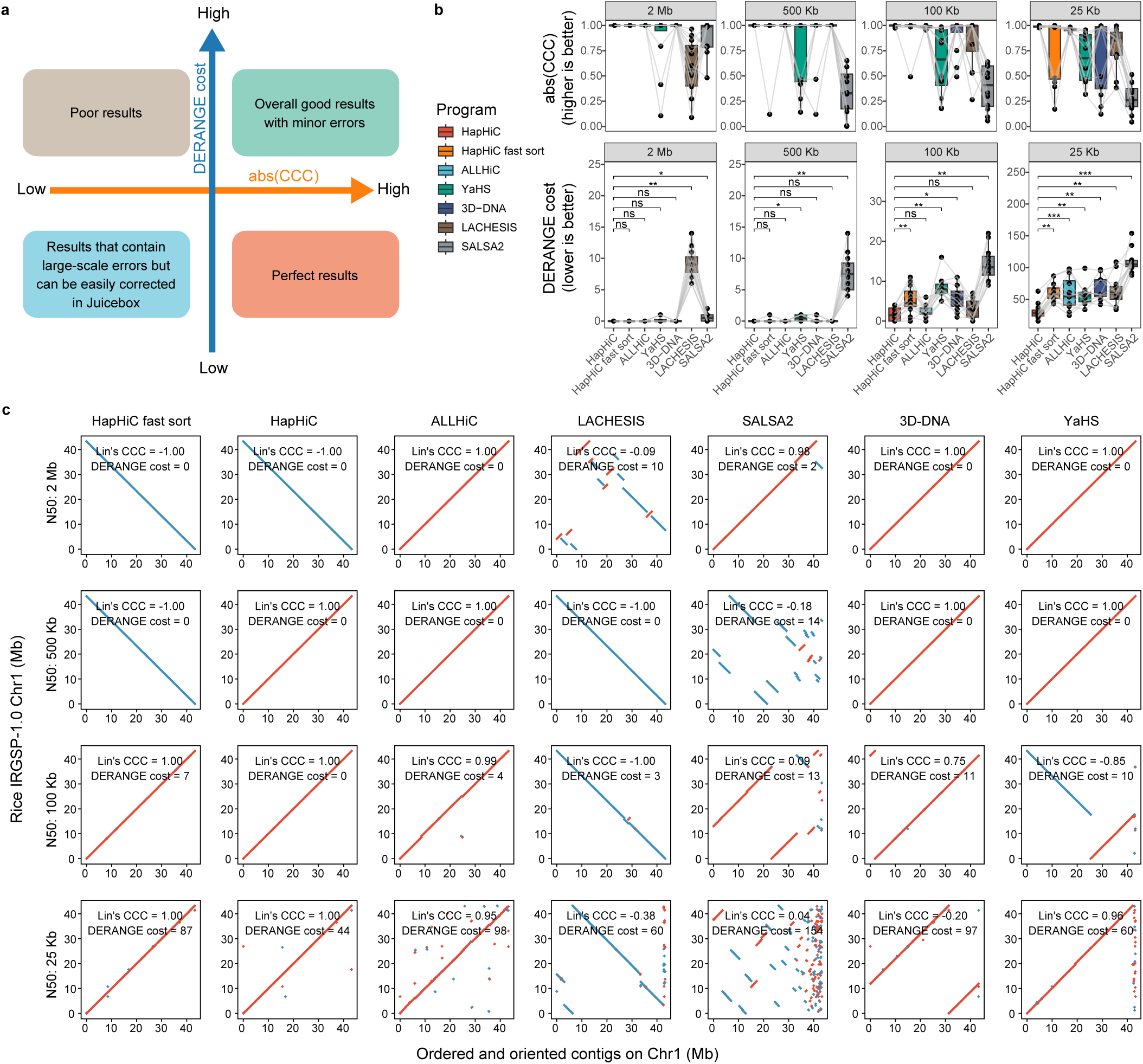
| Evaluation of Hi-C-based scaffolding tools’ performance in contig ordering and orientation across assemblies with varying contig N50 values. **a**, A schematic diagram categorizes the contig ordering and orientation results into four quadrants based on their distinct tendencies, using the absolute value of Lin’s concordance correlation coefficient (CCC) and DERANGE cost as metrics. The former metric assesses the large-scale consistency between the results and the reference chromosomes, while the latter one quantifies the agreement on a smaller scale. **b**, The absolute values of Lin’s CCC and DERAGE costs for each Hi-C-based scaffolding tool in ordering and orienting the contigs of the rice IRGSP-1.0 chromosomes with varying contig N50 values are presented (*p* value from two-sided Wilcoxon signed-rank tests, *n* = 12). **c**, The dot plots illustrate the concordance between chromosome 1 of the rice IRGSP-1.0 genome and the contig ordering and orientation result produced by each Hi-C-based scaffolding tool. Red and blue dots represent forward and reverse complementary alignments, respectively.

**Fig. 3b,c** and **Supplementary Figs. 42-45** illustrate the performance of each scaffolding tool in terms of contig ordering and orientation (**Supplementary Data 2**). SALSA2 performed poorly with a low contig N50, exhibiting the lowest absolute values of CCC and highest DERANGE costs among the evaluated tools when the contig N50 was less than or equal to 500 Kb. In contrast, LACHESIS struggled with high contig contiguity, exhibiting a trend opposite to that of SALSA2. YaHS primarily generated large-scale errors, as indicated by the relatively high absolute values of CCC, while the small-scale errors it produced were at an average level with moderate DERANGE costs. In line with previous findings, 3D-DNA and ALLHiC outperformed LACHESIS and SALSA2 in all aspects, producing fewer large- and small-scale errors. As expected, HapHiC yielded results similar to 3D-DNA when only fast sorting was applied due to the use of the same algorithm. Furthermore, additional optimization using the genetic algorithm significantly reduced small-scale errors, particularly when the contig contiguity was low. This optimization allowed HapHiC to excel in the ordering and orientation of contigs.

### Execution speed and memory usage

HapHiC not only reduces the number of iterations in the genetic algorithm by introducing fast sorting, but it also optimizes the storage and transfer efficiency of Hi-C links. These optimizations result in significant improvements in wall time, CPU time, and peak memory usage during the process of contig ordering and orientation compared to ALLHiC (**Fig. 4a-c** and **Supplementary Data 3**).

**Fig. 4.**
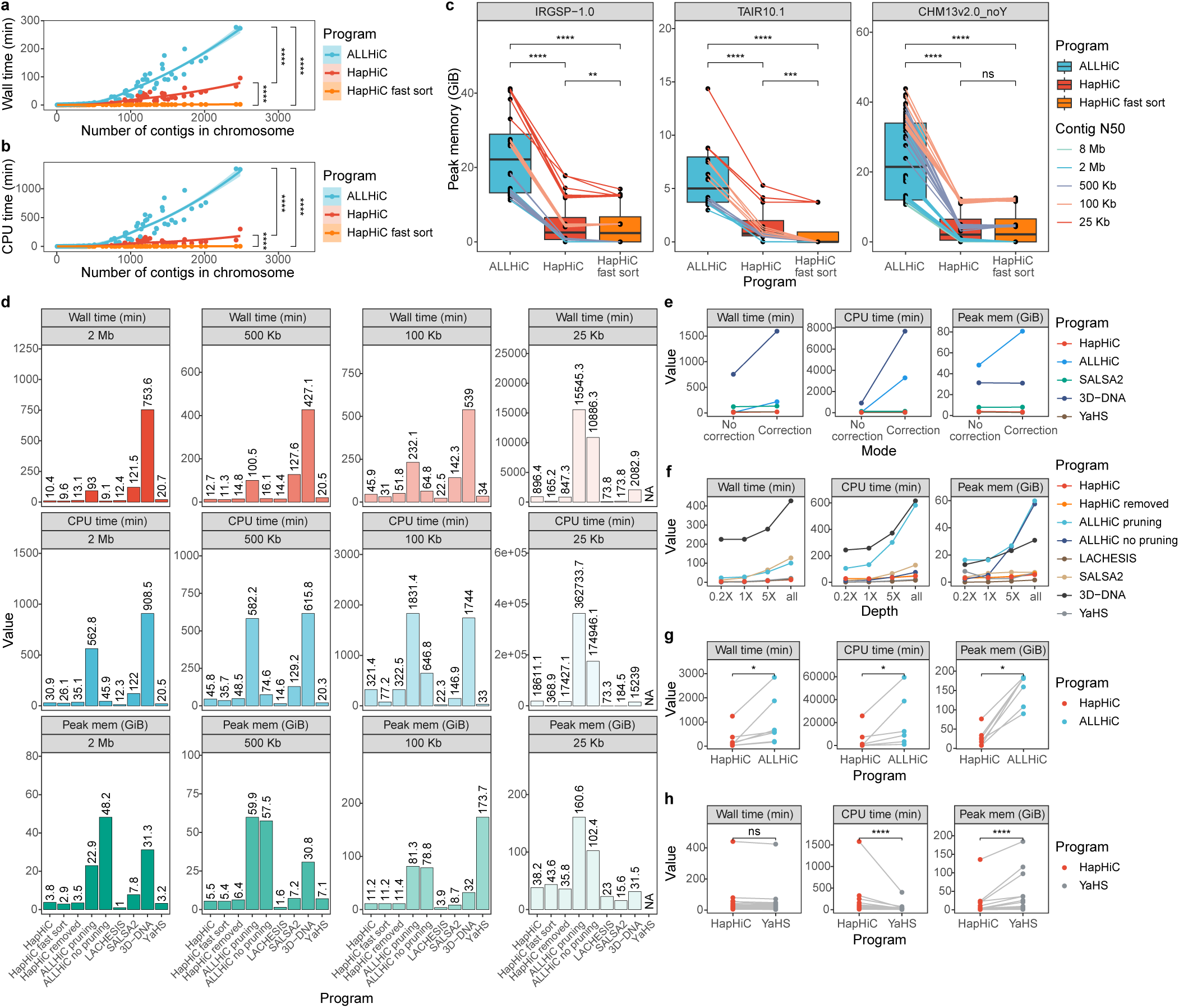
| Comparative analysis of execution time and memory usage for Hi-C-based scaffolding tools. **a**-**c**, A comparative analysis of execution time and memory usage between HapHiC and ALLHiC during contig ordering and orientation. The wall time (**a**, *n* = 160), CPU time (**b**, *n* = 160), and peak memory (**c**, *n* = 48, 20, 92, respectively) for HapHiC and ALLHiC were recorded for each chromosome of rice IRGSP-1.0, *Arabidopsis* TAIR10.1, and human CHM13v2.0_noY genomes under varying contig N50 values. **d**, The total time and memory usage of the entire pipeline for each Hi-C-based scaffolding tool while scaffolding genome assemblies with different contig N50 values simulated from the *M*. *sativa* ground truth. **e**, The time and memory usage of each Hi-C-based scaffolding tool during assembly correction. **f**, The time and memory usage of each Hi-C-based scaffolding tool while processing Hi-C data at different depths. **g**, Comparisons of execution time and memory usage between HapHiC and ALLHiC when scaffolding published haplotype-resolved assemblies (*n* = 7). **h**, Comparisons of execution time and memory usage between HapHiC and YaHS when scaffolding published haplotype-collapsed assemblies (*n* = 20). *P* values were derived from two-sided Wilcoxon signed-rank tests.

Additionally, we conducted a comparative analysis of the execution speed and memory usage of all evaluated Hi-C-based scaffolding tools (**Fig. 4d** and **Supplementary Data 3**). Under varying levels of contig contiguity and sequencing depth, LACHESIS emerged as the most efficient tool. YaHS exhibited satisfactory execution speed but demonstrated a significant increase in peak memory usage with decreasing contig N50. When the contig contiguity was high, SALSA2 was several times slower than the scaffolding tools in the highest speed category and showed a higher susceptibility to low sequencing depth. 3D-DNA performed even worse than SALSA2, proving to be the slowest among all. The efficiency of the ordering and orientation process in ALLHiC was significantly hampered by the decline of contig contiguity. As a result, when the contig N50 was 25 Kb, it took more than ten thousand hours to complete the entire ALLHiC pipeline. Furthermore, 3D-DNA and ALLHiC required more time to correct contigs (**Fig. 4e**) and handle data with high sequencing depth (**Fig. 4f**).

By exclusively applying fast sorting, HapHiC achieved a remarkably high execution speed, second only to LACHESIS (**Fig. 4d**). With the optimization step, HapHiC only fell behind LACHESIS and SALSA2 at extremely low contig contiguity, significantly outperforming 3D-DNA and ALLHiC. Meanwhile, the cost for correcting contigs, removing inter-allele links, and processing high-depth sequencing data were relatively low in HapHiC (**Fig. 4e,f**). Moreover, HapHiC demonstrated stable and excellent performance in terms of memory usage. We also validated the superior efficiency of HapHiC in numerous published genomes compared to ALLHiC and YaHS (**Fig. 4g,h** and **Supplementary Data 3**). Overall, HapHiC maintains a highly competitive execution speed and memory usage while being capable of dealing with more complex assemblies and providing superior scaffolding results.

### Examples of scaffolding published assemblies

We further validated the scaffolding performance of HapHiC in real cases (**Supplementary Data 4**). First, we analyzed several published autopolyploid assemblies and compared the results with those of ALLHiC.

*Saccharum spontaneum* AP85-441 (1*n*=4*x*=32) is the first published haplotype-resolved autotetraploid genome that scaffolded to chromosome level^16^. Due to its highly repetitive nature of genome, combined with the use of Illumina short reads and PacBio RS II data, the final assembly has a contig N50 of only 45 Kb and contains numerous collapsed contigs. In our tests, ALLHiC successfully separated contigs from different homologous chromosomes and produced chromosome or near-chromosome level scaffolds (**Supplementary Fig. 46b**). However, it still introduced noticeable misassignments between non-homologous chromosomes, consistent with previous simulation results. In contrast, HapHiC showed significantly fewer misassignments and accurately clustered contigs to 32 complete chromosomes (**Supplementary Fig. 46a**), greatly reducing the need for manual adjustment in Juicebox. Similarly, HapHiC also produced more accurate and contiguous results in scaffolding the autotetraploid genome assemblies of *M*. *sativa* XinJiangDaYe^17^ (**Supplementary Fig. 47**) and Zhongmu-4^29^ (**Supplementary Fig. 48**).

In 2022, the genome of *S. spontaneum* Np-X, another autotetraploid sugarcane with a different basic chromosome number (*x*=10), was published^20^. Despite its higher contiguity compared to AP85-441, with a contig N50 of 381 Kb, it contains a considerable number of chimeric contigs (**Fig. 5a** and **Supplementary Data 5**). Consequently, we performed assembly correction prior to scaffolding. Although both tools scaffolded contigs into chromosome-level pseudomolecules, ALLHiC produced more misassignments between non-homologous chromosomes (**Supplementary Fig. 49**). Furthermore, the contig N50 of ALLHiC scaffolds dramatically dropped to only 139 Kb after correction, while HapHiC experienced a much milder decrease of 8.7% (**Supplementary Table 12**). We randomly selected a subset of contigs from the first haplotype of chromosome 1 and classified them as non-chimeric contigs, chimeric contigs formed between homologous chromosomes, and chimeric contigs formed between non-homologous chromosomes (**Fig. 5a** and **Supplementary Data 5**). Among the chimeric contigs, HapHiC detected 12 out of 16 (75%) formed between homologous chromosomes and 5 out of 6 (83.3%) formed between non-homologous chromosomes. ALLHiC demonstrated higher sensitivity with a detection rate of 100%. However, it misidentified over 105 out of 112 (93.8%) non-chimeric contigs as chimeric contigs. In contrast, HapHiC exhibited superior stringency, as none of the non-chimeric contigs were mislabeled as chimeric contigs. Furthermore, analysis of the specific breakpoints revealed that ALLHiC tended to break contigs at the positions distant from misjoin points (**Fig. 5b** and **Supplementary Fig. 50**). These issues finally led to a significant reduction in the contig contiguity after ALLHiC correction. In conclusion, HapHiC adopted a more stringent strategy to maintain contig contiguity without sacrificing accuracy of chromosome assignment.

**Fig. 5.**
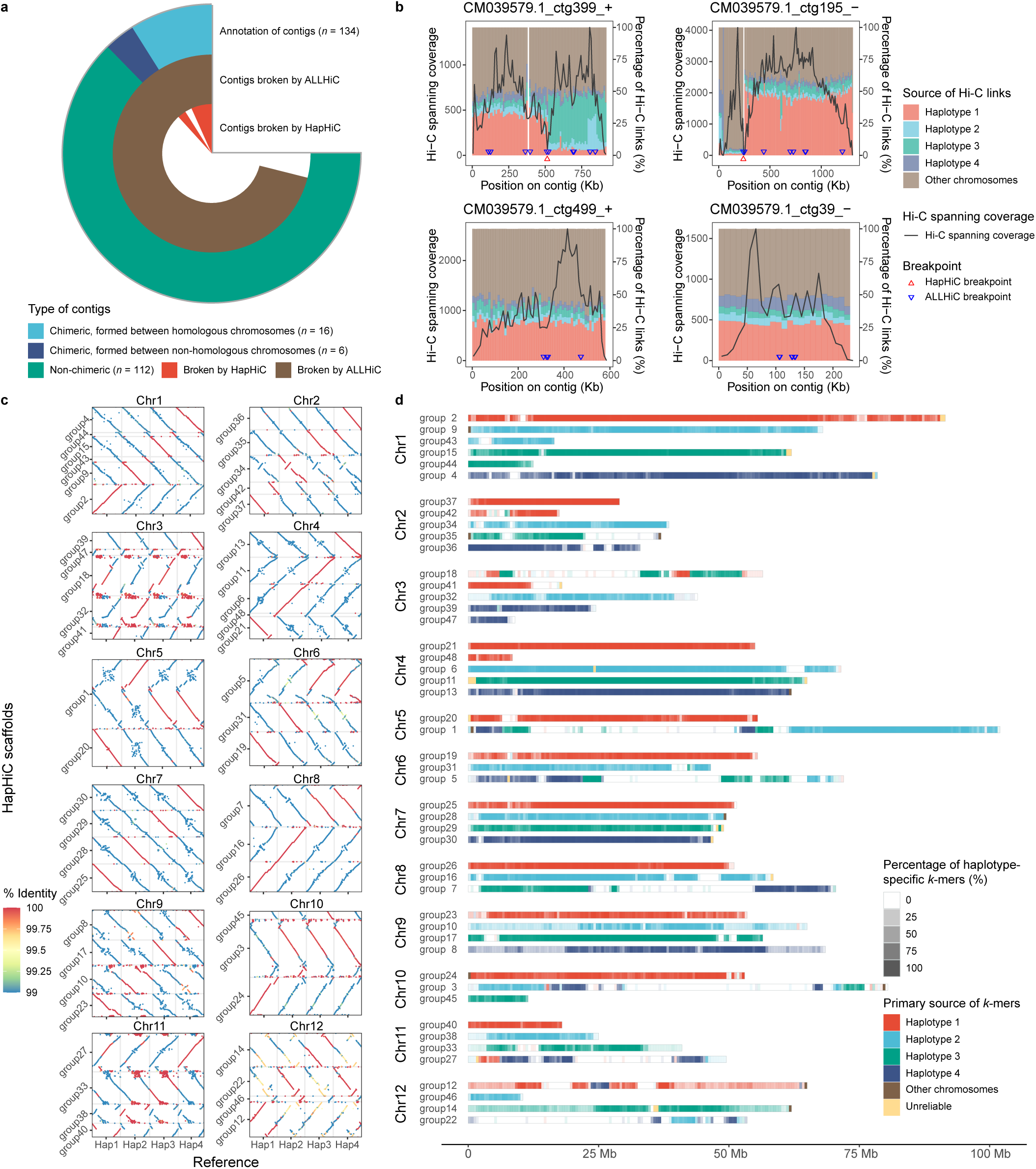
| Comparative analysis and examples of HapHiC in scaffolding published autotetraploid genomes. **a**, A comparison of assembly correction between HapHiC and ALLHiC for the *S*. *spontaneum* Np-X genome. **b**, Examples of assembly correction by HapHiC and ALLHiC, including a chimeric contig formed between homologous chromosomes (CM039579.1_ctg399_+), a chimeric contig formed between non-homologous chromosomes (CM039579.1_ctg195_−), and two non-chimeric contigs (CM039579.1_ctg499_+ and CM039579.1_ctg39_−). The line charts depict the Hi-C spanning coverages along contigs (left axes), while histograms represent the percentages of Hi-C links based on their sources along contigs (right axes). The source of each Hi-C link is determined by the mapping position of the other end of the read pair. Red and blue triangles indicate the breakpoints determined by HapHiC and ALLHiC, respectively. **c**, The dot plots illustrate the alignments between the HapHiC scaffolds and the haplotypes of potato C88 genome, with dot colors indicating the sequence identities of alignments. **d**, A *k*-mer-based analysis reveals the primary source of each position along the contigs from the potato C88 haplotypes. The color indicates the primary source of *k*-mers, while the degree of transparency represents the percentage of haplotype-specific *k*-mers.

We also conducted an analysis of the autotetraploid potato (*Solanum tuberosum*) C88 genome^30^. In contrast to wild plants, the domestication and breeding history of the cultivated potato has left footprints in its haplotypes, resulting in patchy distribution of large, nearly identical regions. These regions make conventional genome assembly and scaffolding methods unable to accurately represent the haplotypes of C88 genome, even when utilizing ONT ultra-long reads^15^. Therefore, the researchers incorporated additional genetic population information to assist in resolving the C88 haplotypes. Another similar case is the autotetraploid potato cultivar Otava^31^. To evaluate the effectiveness of HapHiC and ALLHiC in scaffolding such a complex genome, we first assembled the C88 genome using conventional methods without employing genetic population information. The total length of the assembled unitigs is 3.22 Gb, with an N50 of 730 Kb.

There are no large-scale regions of low divergence in the haplotypes of chromosomes 1, 4, 7, and 9 in the C88 genome^30^. This suggests that these haplotypes can be resolved more easily without relying on genetic population information. HapHiC effectively separated them into chromosome or near-chromosome scaffolds (**Fig. 5c,d** and **Supplementary Fig. 52a**). Additionally, all haplotypes of chromosome 2 and those haplotypes with evenly distributed unique polymorphic loci were accurately represented. Although some haplotypes were not correctly clustered by HapHiC, this issue can be attributed to the existence of large-scale regions of low divergence. In contrast, ALLHiC consistently misassigned contigs from different haplotypes into the same clusters for all C88 chromosomes (**Supplementary Fig. 51** and **Supplementary Fig. 52b**), indicating significantly reduced performance compared to HapHiC.

In addition to autopolyploids, HapHiC outperformed ALLHiC in allele-aware scaffolding of phased diploid assembly of the *Camellia sinensis* Tieguanyin genome^19^. HapHiC exhibited significantly higher scaffold contiguity and fewer misassignments between both homologous and non-homologous chromosomes (**Supplementary Fig. 53**). Furthermore, HapHiC can also scaffold haplotype-collapsed allopolyploid (**Supplementary Figs. 54-58**) and diploid assemblies (**Supplementary Figs. 59-73**). Importantly, HapHiC is not limited to plants. It has been successfully validated in scaffolding representative genomes from various taxa, including humans, birds, amphibians, fish, insects, mollusks, and annelids (**Supplementary Figs. 59** and **68-73**). In these cases, HapHiC achieved comparable or even better performance than YaHS.

The results of real cases demonstrate not only the robustness and reliability of HapHiC in scaffolding various assemblies, but also its potential in overcoming the challenges posed by more complex genomes.

### Application of HapHiC in constructing the haplotype-resolved genome of *M*. × ***giganteus***

*M*. × *giganteus* is widely recognized as a promising lignocellulosic bioenergy crop due to its perennial nature, rapid growth, high productivity, and low input requirements^32^. It is an allotriploid (2*n*=3*x*=57, ABB) formed through natural intrageneric hybridization between the diploid *Miscanthus sinensis* (AA) and the autotetraploid *Miscanthus sacchariflorus* (BBBB)^33^. Additionally, the common ancestor of the *Miscanthus* genus experienced a recent whole-genome duplication (WGD) event prior to this hybridization^34^, resulting in the hexaploidy nature of the *M*. × *giganteus* genome. Despite recent publication of genomes for several other *Miscanthus* species^34–37^, decoding the *M*. × *giganteus* genome is hindered by its complexity. With the help of HapHiC, here we present the first chromosome-level haplotype-resolved genome of *M*. × *giganteus*.

A total of 69.4 Gbp of PacBio HiFi reads and 684.1 Gbp of Hi-C reads were generated for genome assembly and scaffolding, respectively. After assembly and contamination filtration, we obtained phased unitigs with a total length of 6.11 GB, which represents 90% coverage of the genome size as determined by flow cytometry. The assembly was then scaffolded using HapHiC and ALLHiC separately. HapHiC outperformed ALLHiC with significantly fewer misassignments (**Supplementary Fig. 74**). Finally, we anchored contigs accounting for 98.3% of the total assembly onto 57 haplotype-resolved chromosomes based on the HapHiC scaffolds. The contig N50 reached 2.18 Mb, surpassing all existing genome assemblies within the *Miscanthus* genus (**Supplementary Table 13**).

The structural accuracy of the *M.* × *giganteus* genome was subsequently evaluated through gene synteny analysis. As previously mentioned, the A and B subgenomes of *M.* × *giganteus* originated from the genomes of *M*. *sinensis* and *M*. *sacchariflorus*, respectively. Therefore, it is expected that the A subgenome of *M*. × *giganteus* (MgiA) would be phylogenetically closer to the *M*. *sinensis* genome^34^ (MsiA) than to the B subgenomes of *M*. × *giganteus* (MgiB1, MgiB2). However, the gene synteny analysis yielded contradictory results, revealing that MgiA shares higher similarity with the B subgenomes, MgiB1 and MgiB2, than MsiA (**Fig. 6a**). The finding was also supported by the *Miscanthus lutarioriparius* genome^36^ (MluB in **Fig. 6a**), which serves as an alternative B genome of *M*. *sacchariflorus*^35^ with higher completeness and contiguity.

**Fig. 6.**
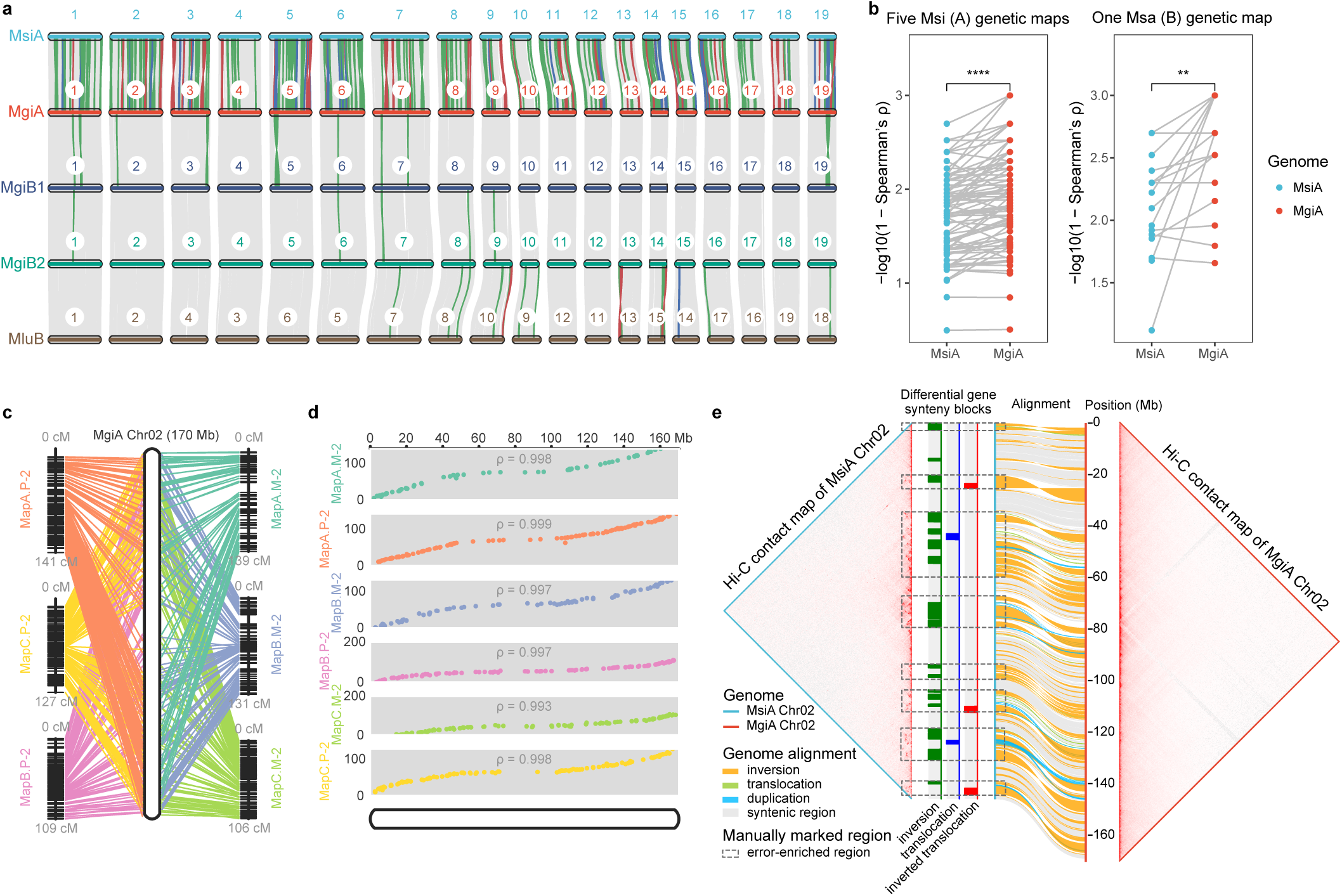
| Comparative genomic analysis of *M*. × *giganteus* and other *Miscanthus* species. **a**, Gene synteny analyses between each subgenome haplotypes of *M*. × *giganteus* (MgiA, MgiB1, and MgiB2) and the genomes of *M*. *sinensis* (MsiA) and *M*. *lutarioriparius* (MluB). Ribbons illustrate gene synteny blocks between orthologous chromosome pairs, with green, blue, and red ribbons representing inversions, translocations, and inverted translocations, respectively. **b**, A comparison between MsiA and MgiA chromosomes based on their alignments with five genetic maps of *M*. *sinensis* (*n* = 94) and one genetic map of *M*. *sacchariflorus* (*n* = 19). Spearman’s correlation coefficients (ρ) were calculated to quantify the agreements between genetic maps and genomes using ALLMAPS. P values were derived from two-sided Wilcoxon signed-rank tests on the raw ρ values. **c**, Alignments between the physical positions on the chromosome 2 of MgiA and the positions on the six genetic maps. **d**, A correlation analysis between the physical positions on the chromosome 2 of MgiA and the corresponding map positions using Spearman’s correlation coefficients. **e**, A comparison of Hi-C contact maps between the chromosomes 2 of MsiA and MgiA. Genome alignment between the two chromosomes is shown in ribbons, with yellow, green, and blue ribbons representing inversions, translocations, and duplications identified by SyRI. The inconsistent gene synteny blocks identified in the gene synteny analysis (**a**) are also shown in this plot. Dashed rectangles highlight error-enriched regions in MsiA chromosome 2.

To determine the authenticity of this observation, we compared the MgiA and MsiA structures using genetic maps and Hi-C contact maps. Five genetic maps of *M*. *sinensis* and one genetic map of *M*. *sacchariflorus*^38^ showed a significantly stronger correlation with MgiA than with MsiA (**Fig. 6b-d** and **Supplementary Data 6**). Additionally, the Hi-C contact maps revealed substantial errors within the MsiA genome, primarily concentrated in the divergence-enriched regions identified through gene synteny analysis and genome alignment (**Fig. 6e**). These findings strongly suggest that the A subgenome of *M.* × *giganteus* has a more accurate structural organization compared to the previously published *M*. *sinensis* genome. The construction of the high-quality genome of *M.* × *giganteus* not only facilitates its genetic breeding but also provides improved reference genomes for its hybridization parents, *M*. *sinensis* and *M*. *sacchariflorus*. This reaffirms the effectiveness and accuracy of HapHiC as an allele-aware scaffolding tool for handling such a complex polyploid genome.

## Discussion

The advancement of sequencing techniques and genome assemblers has ushered in a new era of haplotype-resolved genome research. To tackle the challenges presented by species diversity and varying genome characteristics, there is a pressing need for a robust and efficient allele-aware scaffolding tool with minimal restrictions. One such restriction is the reliance on a reference genome. Although this can be alternatively achieved by assembling and annotating a haplotype-collapsed genome as the reference^19^, it greatly increased the time and cost for genome construction. Moreover, our results have shown the drawbacks of using a reference genome. HapHiC overcomes this limitation by achieving allele-aware scaffolding without relying on reference genomes, demonstrating greater tolerance for assembly errors. Our simulations and real-case tests have demonstrated the superior reliability of HapHiC compared to other software, with broad support for a wide range of taxa and ploidies. Additionally, the entire pipeline requires less time and memory resources and reduces the need for parameter tuning and manual adjustment.

To effectively address a problem, it is crucial to thoroughly understand it. This study conducted exhaustive simulations and evaluations of the factors that could impede the allele-aware scaffolding of phased assemblies on various widely used Hi-C-based scaffolding tools. In addition to the factors mentioned in the ALLHiC paper^7^, our assessment also considered other factors such as contig length distribution, effective Hi-C sequencing depth, and ploidy. Notably, our analysis revealed that diagonally distributed Hi-C links between haplotypes results from switch errors in the initial genome assemblies rather than inherent attributes such as sequence divergence. These findings offer new insights into the challenges of allele-aware scaffolding and pave the way for the development of improved tools.

The formation of collapsed contigs primarily results from extremely low sequence divergence. To mitigate the adverse effects of collapsed contigs, HapHiC has implemented the rank-sum algorithm. However, large-scale collapsed regions still significantly impede subsequent allele-aware scaffolding, as demonstrated in the cultivated potato C88 genome^30^. Furthermore, unlike chimeric contigs, scaffolding tools typically do not correct collapsed contigs. Therefore, achieving a higher quality assembly remains a fundamental prerequisite for haplotype resolution. Otherwise, the resulting scaffolds will still suffer from the “garbage in, garbage out” phenomenon, which means that flawed input data will produce low-quality output. This holds true even when using a scaffolding tool with a high tolerance for assembly errors.

HapHiC still has some limitations. Its accurate clustering in HapHiC relies on prior knowledge of the chromosome number, as well as empirical preferences for length distribution of clusters and chromosomes. To address this, HapHiC provides a straightforward way to understand genome features or manually establish chromosome boundaries through fast sorting without clustering, similar to the intermediate result “0.assembly” in 3D-DNA^5^. Additionally, HapHiC identifies allelic contig pairs based on the distribution pattern of Hi-C links between them. Unlike ALLHiC, this function may not be effective when there are only a few links present. Fortunately, such few links typically do not adversely affect allele-aware chromosome scaffolding.

## Methods

### Overall allele-aware scaffolding strategy of HapHiC

Assembly errors are common in the phased assemblies of heterozygous genomes. Genome assemblers may misjoin nonadjacent sequences, forming chimeric contigs, or merge multiple similar regions into a consensus sequence, resulting in collapsed contigs (**Fig. 2a**). These errors often occur between homologous chromosomes, making chromosome assignment challenging. Scaffolding tools such as 3D-DNA^5^, SALSA2^6^, and YaHS^8^ determine scaffold or chromosome boundaries during or after the contig ordering and orientation. Although this approach does not require prior knowledge of the chromosome count, it exacerbates the adverse effects of assembly errors on chromosome assignment by disrupting the contig ordering and orientation. Therefore, HapHiC employs the same divide-and-conquer strategy as LACHESIS^3^ and ALLHiC^7^, addressing the chromosome assignment problem through clustering before the ordering and orientation of contigs within each chromosome (**Fig. 1**).

Additionally, HapHiC implements four optional preprocessing steps (**Fig. 1b**) to enhance clustering: (1) correcting chimeric contigs using Hi-C link spanning coverage; (2) filtering out low-information contigs, such as short contigs and those lacking Hi-C links; (3) discarding potential collapsed contigs and residual chimeric contigs; (4) removing Hi-C links between allelic contig pairs based on the distribution pattern of Hi-C links. These preprocesses enable HapHiC to perform allele-aware clustering and increase its tolerance towards assembly errors.

### Correcting chimeric contigs

Similar to other scaffolding tools, HapHiC detects misjoins by analyzing the spanning coverage of Hi-C reads at each contig position (**Supplementary Fig. 11**). To accurately determine breakpoints, this coverage is calculated by counting the number of Hi-C read pairs spanning each contig position with a bin size of 500 bp. HapHiC differs from other tools by applying stricter criteria to ensure contig contiguity. Specifically, HapHiC identifies a low-coverage region bounded by two large high-coverage regions as a reliable misjoin and breaks it. Low- and high-coverage regions are contiguous bins divided by one-fifth of the median coverage. By default, the threshold for a large region is the larger of either one-tenth of a contig or 5000 bp.

### Filtering out low-information contigs

Low-information contigs are defined as those that meet one or more of the following criteria: (1) a length shorter than N80, (2) fewer than five restriction sites, or (3) a Hi-C link density below one-fifth of the median value. The Hi-C link density of a contig is calculated by dividing the number of Hi-C links connected to all other contigs by the number of restriction sites within it. These contigs are filtered out before preliminary clustering because they are error-prone and can significantly reduce clustering efficiency.

### Discarding collapsed and chimeric contigs

First, contigs with a Hi-C link density exceeding 1.9 times the median value are identified as potential collapsed contigs and removed. Next, the rank-sum algorithm calculates a rank-sum value for each contig by measuring neighborhood density (**Supplementary Fig. 15**). Let 𝐺 = (𝑉, 𝐸) be a network, where 𝑉 represents the set of contigs as vertices and 𝐸 represents the set of Hi-C links as edges (**Supplementary Fig. 15a**). For any contig 𝑣 ∈ 𝑉, let 𝑁(𝑣, 𝑛) denote the set of contigs with the top 𝑛 Hi-C links connected to 𝑣 . For any two contigs 𝑢, 𝑤 ∈ 𝑁(𝑣, 𝑛), let 𝑟𝑎𝑛𝑘(𝑢, 𝑤) represent the minimum rank of the number of Hi-C links between 𝑢 and 𝑤 (**Supplementary Fig. 15b,c**). The final rank-sum value is given by:

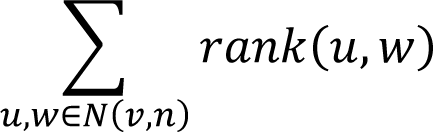

By default, 𝑛 is set to ten. The higher the value, the lower the neighborhood density. Because collapsed and chimeric contigs are respectively merged and misjoined from nonadjacent sequences, their neighborhood densities will be much lower than normal contigs, as reflected by the relatively high rank-sum values (**Supplementary Figs. 16** and 22). Consequently, contigs with a rank-sum value greater than the third quartile (Q3) plus 1.5 times the interquartile range (IQR) are considered remaining collapsed or uncorrected chimeric contigs and are discarded during preliminary clustering.

### Identifying allelic contig pairs and removing inter-allele Hi-C links

HapHiC eliminates the need for reference genomes by identifying allelic contigs based on the distribution pattern of Hi-C links (**Supplementary Fig. 33**). Similar to sequence alignment between allelic contigs, the coordinates of inter-allele Hi-C links are distributed along the diagonal with a slope of 1 or -1 (**Supplementary Figs. 30** and **33a,b**), which differs from the pattern within a chromosome. We introduce a “concordance ratio” to quantify the proportion of Hi-C links that conform to this distribution (**Supplementary Fig. 33b**). The algorithm is described in detail below.

Given a pair of contigs, we construct a coordinate system using the coordinates of 𝑛 Hi-C links connecting them, where 20 ≤ 𝑛 ≤ 200 (randomly selected if exceeding 200). We then use two sliding lines with slopes of 1 and -1 (i.e., 𝑦 = 𝑥 + 𝑏 and 𝑦 = −𝑥 + 𝑏, where 𝑏 is a variable intercept) to calculate the maximum number of coordinate pairs within a certain distance from the lines, denoted as 𝑚. The final concordance ratio is ^𝑚^. The distance is dynamically defined as 1/100 of the length of the shorter contig, with a minimum value of 5 Kb. The higher the value of the concordance ratio, the more it indicates that this pair of contigs conforms to the distribution pattern of Hi-C links between allelic contigs. By default, contig pairs with a concordance ratio greater than 0.2 are considered allelic (**Supplementary Fig. 34**).

Next, an undirected weighted graph is constructed, where vertices denote contigs, edges represent the allelic relationships of contigs, and the edge weights indicate the number of Hi-C links between contigs (**Supplementary Fig. 33**c). Maximum cliques are identified in the graph, and weak edges are removed to divide these cliques into subcliques with the known ploidy as the maximum number of vertices. To retain intra-haplotype links while removing unfavorable inter-haplotype Hi-C links, contigs from the same haplotypes are determined by solving the maximum weighted matching problem across subcliques using a Hungarian algorithm. Finally, Hi-C links from both non-maximal matches and allelic contig pairs are removed before preliminary clustering.

### Clustering

ALLHiC^7^ and LACHESIS^3^ use an agglomerative hierarchical clustering algorithm to cluster contigs into chromosomes. However, specifying the number of clusters (𝑘) to be the number of chromosomes in this method often fails to accurately separate homologous chromosomes in phased assemblies^16, 17^. In such cases, substantial parameter tuning on the 𝑘 values is necessary to improve the clustering results. On the contrary, HapHiC employs a random walk-based Markov cluster algorithm^21^ (MCL) for the initial clustering process. This robust and scalable unsupervised clustering algorithm has proven effective in constructing protein-protein interaction (PPI) networks^39^, clustering orthologous gene families^40^, and analyzing gene synteny^41^. Unlike agglomerative hierarchical clustering, Markov clustering does not specify *k* but regulates granularity with different inflation values. Various clustering results of different granularities can be achieved within a limited range of inflation values. HapHiC then determines the optimal inflation based on the known number of chromosomes, the actual number of clusters, and the length distribution of clusters.

First, the Markov clusters for each inflation value are sorted in descending order by their cluster length, which is the sum of contig lengths within them. Next, we calculate the length ratio for each adjacent cluster pair. Empirically, if the length ratio between adjacent clusters is far below 1, it could indicate unseparated contigs from different chromosomes. Consequently, we prefer the results with progressively decreasing cluster lengths, where the number of clusters meeting the criterion should be greater than or equal to the known number of chromosomes. The pseudocode of the algorithm can be found in **Supplementary Table 14**. By default, HapHiC initially uses a length ratio threshold of 0.75. If none of the clustering results meets this threshold, it will be gradually lowered to 0.5. This method efficiently reduces clustering errors and eliminates the need for manual tuning of 𝑘.

HapHiC optimizes the clustering step for assemblies with varying levels of contiguity. The power-law model of Hi-C links exhibits rapid decay within short distances, while showing gradual decay over longer ranges^7^. As a result, longer contigs are more susceptible to long-range background noise. This is because only their ends are adjacent to other contigs and provide useful information for clustering. Recent tools, such as EndHiC^42^, pin_hic^43^, and YaHS^8^, have addressed this issue primarily by utilizing the Hi-C links from contig ends only. In contrast, HapHiC also considers information from the inner part of each long contig. First, these contigs are split into bins, with a size dynamically defined as 1/30 of the average chromosome length and a maximum value of 2 Mb. These bins are then simply treated as regular contigs during the subsequent construction of the Hi-C link matrix and Markov clustering. The matrix is constructed using links from the ends of contigs or bins with a maximum end length of 500 Kb. After clustering, long contigs are placed in the best clusters based on the total length of bins in each cluster. On the other hand, assemblies of low contiguity often consist of too many contigs, thus remarkably slowing down Markov clustering. To overcome this, we optimize the clustering speed using sparse matrix data structures.

### Reassignment

In the reassignment step, previously filtered-out contigs are rescued and assigned to the most suitable clusters. Unlike ALLHiC rescue^7^, this process also allows for the reassignment of misassigned contigs. All contigs that meeting the minimum requirements for restriction sites and Hi-C links are traversed in descending order to determine their most suitable clusters. First, Hi-C link densities between each contig and MCL cluster are calculated, defined as the number of Hi-C links between a contig and a cluster, divided by the sum of their restriction site counts. The most suitable cluster for each contig is determined based on two criteria: (1) the Hi-C link density to the best cluster is more than four times the average value to other clusters; (2) the density ratio between to the second-best and the best cluster is less than 0.6. By default, reassignment is iterated for up to five rounds to converge, followed by a rescue process for unanchored contigs using only the first criterion.

If the number of clusters still exceeds the number of chromosomes, HapHiC performs an additional agglomerative hierarchical clustering^22^ with average-linkage to obtain the final chromosome-level clusters. To achieve this, a matrix is constructed by calculating the number of Hi-C links between each cluster pair, divided by the product of their restriction site counts. Here, multiplication is used for normalization instead of sum, which differs from the Hi-C link density described above. This approach performs better when cluster lengths differ significantly. The maximum value in the matrix is subtracted from the matrix to obtain the final distance matrix for clustering.

### Ordering and orientation

3D-DNA and ALLHiC have demonstrated excellent performance in contig ordering and orientation compared with other Hi-C-based scaffolding tools^7^. However, their disadvantage is that they are very slow when dealing with a large number of contigs.

The iterative contig ordering and orientation algorithm of 3D-DNA breaks newly connected scaffolds from their exact midpoints, resulting in the trouble that the density graph should be reconstructed from scratch in each iteration^5^. Besides, the intermediate results are read and written through text files. In HapHiC, we simplify the process and call it “fast sorting”. Specifically, we calculate the distance between the midpoint of each newly connected scaffold and the midpoints of all contigs in it to find the nearest one as the approximate midpoint of the scaffold. As a result, the density graph reconstruction in HapHiC can retrieve necessary data directly from memory, eliminating the need for file reading. This improvement makes the iterative ordering and orientation process significantly faster than the original one without compromising the results.

ALLHiC starts contig ordering iterations from randomly shuffled contigs using a genetic algorithm^7^. When handling numerous contigs, the algorithm can be very slow to converge. On the other hand, although the process is parallelized for each cluster, but the Hi-C links between all contigs are stored and processed using a single huge CLM file, leading to a high memory usage. In HapHiC, we use the result of fast sorting as the initial configuration and subsequently employing the ALLHiC algorithm for further optimization. Additionally, the CLM files is split into cluster-specific files, retaining only the Hi-C links within each cluster. This integrated approach allows HapHiC to order and orient contigs more accurately and efficiently than ALLHiC.

### Pseudomolecule Building

In pseudomolecule building, the contigs in each cluster are joined together based on the results of contig clustering, ordering, and orientation. By default, the adjacent contigs are separated with 100-bp Ns as gaps. The sequences and scaffolding information of these pseudomolecules are finally output in the FASTA format and AGP format, respectively.

### Mapping and filtering of Hi-C reads

Fastp^44^ (version 0.21.0) was used to remove adaptors, trim low-quality sequences, and evaluate the quality of raw Hi-C reads. To evaluate the performance of 3D-DNA, Hi-C reads were aligned to assemblies and filtered using the Juicer pipeline^45^. For other scaffolding tools or purposes, Hi-C reads were processed using a uniform method. First, the reads were aligned to assemblies or published genomes using BWA-MEM^46^ (version 0.7.17-r1198-dirty) with the parameter “-5SP”. Next, PCR duplicates, secondary and supplementary alignments were filtered out using SAMBLASTER^47^ (version 0.1.26) and SAMtools^48^ (version 1.11). Finally, a custom script “filter_bam.py” was employed to remove alignments with a mapping quality of zero and an edit distance greater than 2.

### Simulation of datasets with various adverse factors

In the ALLHiC paper, the authors simulated a haplotype-resolved diploid genome by merging both the genome sequences and Hi-C data of two rice subspecies, *indica* and *japonica*^7^. However, this approach has some limitations. First, in a real diploid, there are background interactions between homologous chromosomes, because they are located in the same nucleus. The artificial merging of data from different subspecies leads to an underestimation of Hi-C links between homologous chromosomes, making it easier to separate them even than those between non-homologous chromosomes (**Supplementary Fig. 2**). Second, the conclusions drawn from experiments on diploids may not apply to polyploids. Third, using only a pair of homologous chromosomes for analysis ignores misassignments between non-homologous chromosomes. Therefore, we constructed a ground truth by manually removing obvious assembly errors based on the published haplotype-resolved autotetraploid genome of *M*. *sativa* XinJiangDaYe^17^ (**Supplementary Fig. 1**). We used all 32 chromosomes to construct a ground truth genome of 2.0 Gb.

Using this template, we simulated a series of assemblies with various adverse factors through a pipeline of multiple custom scripts (**Supplementary Fig. 3**). The script “sim_contig.py” fragmented the genome into assemblies with different N50 values (**Supplementary Fig. 4**) and coefficients of variation (CVs) of length (**Supplementary Fig. 6**). Since the contig N50 of the *M*. *sativa* genome is less than 500 Kb, we retained the gaps represented by Ns in the fragments to simulate higher contiguity. For concision, we always refer to these fragments as contigs, regardless of whether there are gaps in them. Based on these assemblies, we used the script “sim_chimeric_contigs.py” to simulate assemblies with different proportions of chimeric contigs (**Supplementary Fig. 10**). Among them, chimeric contigs simulated between homologous chromosomes, non-homologous chromosomes, and within chromosomes were generated at a ratio of 7:2:1, respectively. To simulate different effective sequencing depths, we used “samtools view -s” with 12345 as the seed for random sampling (**Supplementary Fig. 8**).

We simulated haplotypes by introducing single nucleotide polymorphisms (SNPs), insertions, and deletions to the first haplotype in the ground truth genome using the script “sim_haplotypes.py”. By adjusting the proportion of variations and number of haplotypes generated, we obtained genomes with different sequence divergence levels (**Supplementary Fig. 27**) and ploidies. The corresponding Hi-C reads were generated by sim3C^49^, with the parameter “--trans-rate” dynamically adjusted according to different ploidies to maintain a fixed intensity of chromosome interaction. To simulate reality, we introduced SNPs, insertions, and deletions at a ratio of 10:0.5:0.5, with a 2:1 ratio of transitions to transversions for SNPs (**Supplementary Fig. 26**). Furthermore, these variations were unevenly distributed in bins with a length of 100 Kb and a CV of 0.5.

Based on these simulated haplotypes and Hi-C reads, we further generated collapsed contigs (**Supplementary Fig. 20**) and switch errors (**Supplementary Fig. 31**). We used the script “sim_collapsed_regions.py” to create two-, three-, and four-haplotype collapsed contigs in a ratio of 7:2:1. Switch errors were introduced by randomly shuffling variations between haplotypes using the script “sim_switch_errors.py”. We also incorporated all the custom scripts above to introduce 5% each of chimeric, collapsed contigs, and switch errors into assemblies with ploidy ranging from 2 to 16. Each contig in simulated assemblies recorded information such as source chromosome, haplotype, position, and error type for subsequent result evaluation.

Three reference genomes of rice (IRGSP-1.0^24^), *Arabidopsis* (TAIR10.1^25^), and human (CHM13v2.0_noY^1^) were used as ground truths to evaluate the accuracy of Hi-C-based scaffolding tools in contig ordering and orientation. Assemblies with varying contig N50 values were also simulated using the script “sim_contigs.py”. HapHiC, ALLHiC, and LACHESIS perform ordering and orientation for individual chromosomes (clusters), differing from the whole-genome scaffolding of SALSA2, 3D-DNA, and YaHS. Thus, contigs in each simulated assembly were partitioned into corresponding chromosomes and ordered and oriented separately, making the evaluation independent of chromosome assignment. Assembly correction of scaffolding tools was disabled for this experiment.

### Scaffolding performance evaluation

We initially evaluated the performance of Hi-C-based scaffolding tools in chromosome assignment using metrics including contiguity, anchoring rate, misassignment rates, and the number of scaffolds using a custom script “result_statistics.py”.

In simulated assemblies, all chromosomes were fragmented. Thus, only sequences output by scaffolding tools containing multiple contigs are true scaffolds, while individual contigs are classified as unanchored sequences. The anchoring rate is the ratio of the length of contigs in scaffolds to the total length of all contigs. Ideally, the anchoring rate is 100%, and the number of scaffolds equals the number of chromosomes.

We did not use metrics such as N50 to assess scaffold contiguity because incorrectly separated chromosomes could artificially inflate N50 values. Instead, we designed an indicator with a maximum value of one that is independent of the anchoring rate. First, we calculated the cumulative lengths of contigs based on their source chromosomes for each scaffold and identified the longest source chromosome as the dominant chromosome. Subsequently, we divided this length by the total length of dominant chromosome anchored to all scaffolds to obtain a ratio. A ratio of one signifies that the scaffold entirely comprises contigs from a specific chromosome, and all contigs from this chromosome are distributed only within this scaffold. Finally, we calculated the average value of this ratio among all scaffolds to obtain the final contiguity. A contiguity of one indicates that all anchored chromosome sequences correspond perfectly to scaffolds in a one-to-one relationship.

For contigs in scaffolds that do not originate from the dominant chromosome, we categorize them as either misassignments between homologous chromosomes or between non-homologous chromosomes, depending on their actual relationship with the dominant chromosome. The misassignment rate is calculated as the proportion of their length to the total length of the anchored genome. If all contigs in the scaffolds originate from the dominant chromosome, the misassignment rate is zero.

Chimeric and collapsed contigs are excluded from statistics because they cannot be considered totally correct when placed in any scaffold. During ALLHiC pruning, the genome of a closely related species, *Medicago truncatula* (MtrunA17r5.0-ANR^50^), was used as a reference. However, chromosomes 4 and 8 of *M*. *truncatula* have some structural differences compared to those of *M*. *sativa*^17^. Therefore, we also calculated the contiguity and misassignment rate after excluding these two chromosomes. Since we manually tuned the parameter 𝑘 for ALLHiC, we calculated Δ𝑘 as the difference between optimized 𝑘 and default 𝑘.

We also evaluated the accuracy of Hi-C-based scaffolding tools in contig ordering and orientation for assemblies with varying contig N50 values. In preliminary comparisons between HapHiC and ALLHiC, we counted the number of generations for convergence and calculated scores using the “optimize” program of ALLHiC^7^. For a more objective comparison, two metrics were used to evaluate the accuracy of all scaffolding tools. Lin’s concordance correlation coefficients^26^ (CCCs) were calculated using a custom script “draw_tour_file.py” to measure large-scale consistency between results and reference genomes. “Costs” were also calculated using a modified DERANGE II program^27^ with the parameters including linear (-L), signed (-S), and a look-ahead value of three. The weights for inversions, transpositions, and transversions were set to one, one, and two, respectively, to simulate the number of steps needed to achieve optimal results in Juicebox^28^. Costs for both the original order and reverse complementary order were calculated for each scaffolding result, and the minimum of the two values were considered the final cost. For SALSA2 and YaHS, which can output multiple scaffolds for each chromosome, we joined these scaffolds as the result of ordering and orientation. However, for 3D-DNA, the intermediate result “0.assembly” was used to ensure contig completeness and result comparability.

### Measurement of execution time and peak memory usage

All tasks were executed on a server running CentOS Linux (release 7.6.1810). The server is equipped with two Intel Xeon Gold 6132 CPUs (a total of 28 cores at 2.6 GHz) and 192 gibibytes (GiB) of memory. The CPU time, wall time, and peak memory usage of each task were measured using the PBS Professional job scheduler (PBS Pro, version 18.1.4). A custom script, “pbsperf.py,” was utilized to summarize the records and convert the units to minutes (min) and GiB. However, the measurements of peak memory usage may not be precise because the scheduler measures them at intervals. If a task completes within several seconds, PBS Pro may record a peak memory usage of zero. All steps in ALLHiC were executed in parallel using GNU parallel^51^ (version 20210922) to optimize the wall time if possible.

### Validation of HapHiC in real cases

HapHiC was further validated using real cases. We compared its scaffolding performance and resource usage with those of ALLHiC in published haplotype-resolved autotetrapolyploid and diploid genomes. Additionally, we compared HapHiC with YaHS in haplotype-collapsed allotetraploid and diploid genomes of various taxa. The information of all species used in the validation is listed in **Supplementary Data 4**. Apart from the potato C88 genome, which was assembled using hifiasm^12^ (version 0.19.0-r534), the corresponding assemblies of other genomes were generated by breaking the “N” gaps and randomly shuffling the ordering and orientation of contigs using custom scripts “split_fasta.py” and “shuffle_fasta.py”, respectively. All scaffolding results were visualized using Juicebox^28^ (version 1.11.08) for comparisons.

For the potato C88 assembly, we also used a 𝑘-mer-based method to analyze the scaffolds output by HapHiC and ALLHiC using a script “haplotype_kmers.py”. First, we generated 201-mers from the published haplotype-resolved reference genome of potato C88. These 201-mers were then annotated based on their source chromosomes and used to classify each region of HapHiC and ALLHiC scaffolds with a bin size of 500 Kb. Additionally, we aligned these scaffolds to the reference genome using unimap (https://github.com/lh3/unimap, version 0.1-r41) and visualized the alignments with a modified version of paf2dotplot (https://github.com/zengxiaofei/paf2dotplot).

### Sampling, library construction, and genome sequencing of *M*. × *giganteus*

Young leaves of a *M*. × *giganteus* plant were collected at Hunan Agriculture University in Changsha, Hunan Province, P.R. China and immediately frozen in liquid nitrogen. For HiFi sequencing, DNA was extracted from the leaves using a modified CTAB protocol. The DNA was then qualified and quantified using a NanoDrop 2000 Spectrophotometer (NanoDrop Technologies, Wilmington, DE, USA), a Qubit 3.0 Fluorometer (Life Technologies, Carlsbad, CA, USA), and 0.8% agarose gel electrophoresis. Three SMRTbell libraries were constructed using sheared DNA and the SMRTbell Express Template Prep Kit 2.0 (Pacific Biosciences, Menlo Park, CA, USA). The libraries were size-selected with a minimum length of ∼15 Kb using the BluePippin (Sage Science, Beverly, MA, USA) and sequenced on the PacBio Sequel II System under the circular consensus sequencing (CCS) mode for 30-h movies using 2.0 Chemistry. A total of 69.4 Gb of HiFi reads were generated.

For Hi-C sequencing, the leaves were fixed with formaldehyde to cross-link chromatin. After cell lysis, the cross-linked chromatin was digested using the MboI restriction enzyme. Sticky ends were repaired, labeled with biotin, and ligated to form chimeric molecules. Proteins were then digested from the chromatin using protease, and DNA was purified using a QIAamp DNA Mini Kit (Qiagen, Hilden, NRW, Germany) and sheared into fragments of 400-600 bp. Biotin-labeled fragments were enriched using streptavidin-coated magnetic beads (Vazyme, Nanjing, JS, P.R. China) for library construction. Hi-C sequencing was performed on the BGI MGISEQ-2000 platform under the PE150 mode, generating a total of 684.1 Gb of Hi-C reads.

### *De novo* assembly and comparative analysis of *M*. × *giganteus* genome

The genome of *M*. × *giganteus* was assembled using hifiasm^12^ (version 0.13-r308) with HiFi reads. The parameter “-l0” was employed to disable duplication purging, resulting in the primary unitigs (p_utg) with a size of 6.13 Gb and an N50 of 2.18 Mb. After removing organellar and exogenous DNA sequences from these unitigs, a draft assembly of 6.11 Gb with an N50 of 2.19 Mb was obtained. To identify the source of diagonally distributed inter-allele Hi-C links, the genome was also assembled using HiCanu^23^ (version 2.1.1) for comparison with the hifiasm assembly. The hifiasm assembly was scaffolded onto 57 chromosome-level pseudomolecules using HapHiC and ALLHiC separately, both with default parameters. After manual curation in Juicebox^28^ (version 1.11.08), the final chromosome-level haplotype-resolved genome of *M*. × *giganteus* was generated based on the HapHiC scaffolds.

The genes in each haplotype of *M*. × *giganteus* were simply annotated by mapping the coding sequences of the *M*. *sinensis* genome using GMAP^52^ (version 2019-12-01) with the parameter “-n 1”. MCscan^41^ in JCVI utility libraries (version 1.1.18) was used to perform gene synteny comparisons between the subgenomes of *Miscanthus* species and to draw the karyotype plot. Genome alignment between chromosomes of *M*. × *giganteus* and *M*. *sinensis* was performed using Minimap2^53^ (version 2.26-r1175). Structural variations were identified using SyRI^54^ (version 1.6.3) and visualized using plotsr^55^ (version 1.1.1). To compare the structural accuracy of the *M*. × *giganteus* and *M*. *sinensis* genomes, five genetic maps of *M*. *sinensis* and one genetic map of *M*. *sacchariflorus* were collected^38^. The genetic markers of each map were aligned to the two genomes using BWA-ALN and BWA-SAMSE (version 0.7.17-r1198-dirty) with default parameters. The agreements between the genetic maps and the genomes were analyzed and visualized using ALLMAPS^2^ in JCVI utility libraries with the markers shared by all genetic maps.

### Program versions and command lines

All program versions and command lines used in this research are available in **Supplementary Information.**

### Statistics analysis

Two-sided Wilcoxon signed-rank tests were performed to compare scaffold contiguity values, misassignment rates, ALLHiC scores, Lin’s concordance correlation coefficients (CCCs), DERANGE costs, and time and memory usage. The function “wilcox.test” in R language (version 4.0.2) was used with the “paired” parameter set to “TRUE”. The results of statistical tests were visualized using the ggpubr package in R. The “geom_smooth” function in ggplot2 was used to fit curves with the formula “y ∼ x” and the method “loess”. Lin’s CCCs were calculated between the corresponding positions on the reference chromosomes and scaffolds to measure their agreements using the following formula:

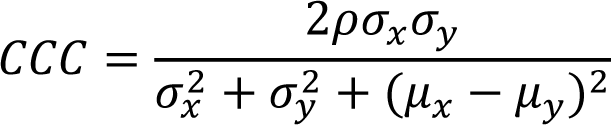

where 𝜌 is the Pearson correlation coefficient between the two variables, 𝜎_𝑥_ and 𝜎_𝑦_are the standard deviations of the two variables, and 𝜇_𝑥_ and 𝜇_𝑦_ are the means of the two variables. Additionally, Spearman’s correlation coefficients were calculated to quantify the agreements between genetic maps and genomes using ALLMAPS^2^.

### Data availability

All raw sequencing data and the final chromosome-level haplotype-resolved genome of *M*. × *giganteus* will be publicly available after publication. All published raw sequencing data and genome assemblies used for HapHiC validation are listed in **Supplementary Data 4**.

### Code availability

HapHiC and all custom scripts for dataset simulation are available on GitHub at https://github.com/zengxiaofei/HapHiC. The source code of modified ALLHiC can be found at https://github.com/zengxiaofei/allhic.

## Acknowledgements

This work was supported by the National Natural Science Foundation of China (32100459), and the fellowship of China Postdoctoral Science Foundation (2020M672695) to Xiaofei Zeng. Additional funding was provided by the Shenzhen Municipal Science and Technology Innovation Commission Foundation (JCYJ20220530114415036, JCYJ20210324104800001), and the National Natural Science Foundation of China (32070625) to G.C.; and the National Natural Science Foundation of China (32222019) to Xingtan Zhang. The Center for Computational Science and Engineering at Southern University of Science and Technology also provided support for this work. We thank Dr. Dong Zhang and Dr. Dafu Ru from Lanzhou University, Mr. Zhigui Bao from the Max Planck Institute for Biology Tübingen, and Dr. Shilin Zhu from Huazhong Agricultural University for their valuable suggestions and contributions to HapHiC.

## Author contributions

Xiaofei Zeng designed the algorithms, implemented HapHiC, analyzed the genome of *M*. × *giganteus*, and wrote the manuscript. Z.Y. and S.Y. managed and provided the plant materials of *M*. × *giganteus*. Xiaofei Zeng, Y.D., Y.L., Z.Z., S.C., and H.Z. benchmarked HapHiC and other scaffolding tools. G.C., Xingtan Zhang, and Y.W. provided suggestions for the algorithms. G.C. and Xingtan Zhang revised the manuscript.

## References

1. Nurk, S. et al. The complete sequence of a human genome. Science 376, 44–53 (2022).

2. Tang, H. et al. ALLMAPS: robust scaffold ordering based on multiple maps. Genome Biology 16, 3 (2015).

3. Burton, J.N. et al. Chromosome-scale scaffolding of *de novo* genome assemblies based on chromatin interactions. Nature Biotechnology 31, 1119–1125 (2013).

4. Putnam, N.H. et al. Chromosome-scale shotgun assembly using an in vitro method for long-range linkage. Genome Research 26, 342–350 (2016).

5. Dudchenko, O. et al. De novo assembly of the *Aedes aegypti* genome using Hi-C yields chromosome-length scaffolds. Science 356, 92–95 (2017).

6. Ghurye, J. et al. Integrating Hi-C links with assembly graphs for chromosome-scale assembly. PLOS Computational Biology 15, e1007273 (2019).

7. Zhang, X., Zhang, S., Zhao, Q., Ming, R. & Tang, H. Assembly of allele-aware, chromosomal-scale autopolyploid genomes based on Hi-C data. Nature Plants 5, 833–845 (2019).

8. Zhou, C., McCarthy, S.A. & Durbin, R. YaHS: yet another Hi-C scaffolding tool. *Bioinformatics (Oxford*, England) 39 (2022).

9. Lieberman-Aiden, E. et al. Comprehensive Mapping of Long-Range Interactions Reveals Folding Principles of the Human Genome. Science 326, 289–293 (2009).

10. Yuan, Y., Scheben, A., Edwards, D. & Chan, T.-F. Toward haplotype studies in polyploid plants to assist breeding. Molecular plant 14, 1969–1972 (2021).

11. Koren, S. et al. *De novo* assembly of haplotype-resolved genomes with trio binning. Nature Biotechnology 36, 1174–1182 (2018).

12. Cheng, H. et al. Haplotype-resolved assembly of diploid genomes without parental data. Nature Biotechnology 40, 1332–1335 (2022).

13. Meyer, R.S., DuVal, A.E. & Jensen, H.R. Patterns and processes in crop domestication: an historical review and quantitative analysis of 203 global food crops. New Phytologist 196, 29–48 (2012).

14. Huang, X., Huang, S., Han, B. & Li, J. The integrated genomics of crop domestication and breeding. Cell 185, 2828–2839 (2022).

15. Cheng, H., Asri, M., Lucas, J., Koren, S. & Li, H. Scalable telomere-to-telomere assembly for diploid and polyploid genomes with double graph. *arXiv preprint arXiv:2306.03399* (2023).

16. Zhang, J. et al. Allele-defined genome of the autopolyploid sugarcane *Saccharum spontaneum* L. Nature Genetics 50, 1565–1573 (2018).

17. Chen, H. et al. Allele-aware chromosome-level genome assembly and efficient transgene-free genome editing for the autotetraploid cultivated alfalfa. Nature Communications 11, 2494 (2020).

18. Wang, P. et al. Genetic basis of high aroma and stress tolerance in the oolong tea cultivar genome. Horticulture Research 8 (2021).

19. Zhang, X. et al. Haplotype-resolved genome assembly provides insights into evolutionary history of the tea plant *Camellia sinensis*. Nature Genetics 53, 1250–1259 (2021).

20. Zhang, Q. et al. Genomic insights into the recent chromosome reduction of autopolyploid sugarcane *Saccharum spontaneum*. Nature Genetics 54, 885–896 (2022).

21. Dongen, S.V. Graph Clustering Via a Discrete Uncoupling Process. SIAM Journal on Matrix Analysis and Applications 30, 121–141 (2008).

22. Pedregosa, F. et al. Scikit-learn: Machine learning in Python. the Journal of machine Learning research 12, 2825–2830 (2011).

23. Nurk, S. et al. HiCanu: accurate assembly of segmental duplications, satellites, and allelic variants from high-fidelity long reads. Genome Research 30, 1291–1305 (2020).

24. Kawahara, Y. et al. Improvement of the *Oryza sativa* Nipponbare reference genome using next generation sequence and optical map data. Rice 6, 4 (2013).

25. Berardini, T.Z. et al. The arabidopsis information resource: Making and mining the “gold standard” annotated reference plant genome. genesis **53**, 474–485 (2015).

26. Lawrence, I.K.L. A Concordance Correlation Coefficient to Evaluate Reproducibility. Biometrics 45, 255–268 (1989).

27. Blanchette, M., Kunisawa, T. & Sankoff, D. Parametric genome rearrangement. Gene 172, GC11–GC17 (1996).

28. Durand, N.C. et al. Juicebox provides a visualization system for Hi-C contact maps with unlimited zoom. Cell systems 3, 99–101 (2016).

29. Long, R. et al. Genome Assembly of Alfalfa Cultivar Zhongmu-4 and Identification of SNPs Associated with Agronomic Traits. Genomics, Proteomics & Bioinformatics 20, 14–28 (2022).

30. Bao, Z. et al. Genome architecture and tetrasomic inheritance of autotetraploid potato. Molecular plant 15, 1211–1226 (2022).

31. Sun, H. et al. Chromosome-scale and haplotype-resolved genome assembly of a tetraploid potato cultivar. Nature Genetics 54, 342–348 (2022).

32. Heaton, E.A. et al. in Advances in Botanical Research, Vol. 56. (eds. J.-C. Kader & M. Delseny) 75–137 (Academic Press, 2010).

33. Chramiec-Głąbik, A., Grabowska-Joachimiak, A., Sliwinska, E., Legutko, J. & Kula, A. Cytogenetic analysis of *Miscanthus* × *giganteus* and its parent forms. Caryologia 65, 234–242 (2012).

34. Mitros, T. et al. Genome biology of the paleotetraploid perennial biomass crop *Miscanthus*. Nature Communications 11, 5442 (2020).

35. De Vega, J., Donnison, I., Dyer, S. & Farrar, K. Draft genome assembly of the biofuel grass crop *Miscanthus sacchariflorus*. F1000Research 10 (2021).

36. Miao, J. et al. Chromosome-scale assembly and analysis of biomass crop *Miscanthus lutarioriparius* genome. Nature Communications 12, 2458 (2021).

37. Zhang, G. et al. The reference genome of *Miscanthus floridulus* illuminates the evolution of Saccharinae. Nature Plants 7, 608–618 (2021).

38. Dong, H. et al. Winter hardiness of *Miscanthus* (II): Genetic mapping for overwintering ability and adaptation traits in three interconnected *Miscanthus* populations. GCB Bioenergy 11, 706–726 (2019).

39. Brohée, S. & van Helden, J. Evaluation of clustering algorithms for protein-protein interaction networks. BMC Bioinformatics 7, 488 (2006).

40. Li, L., Stoeckert, C.J. & Roos, D.S. OrthoMCL: Identification of Ortholog Groups for Eukaryotic Genomes. Genome Research 13, 2178–2189 (2003).

41. Tang, H. et al. Unraveling ancient hexaploidy through multiply-aligned angiosperm gene maps. Genome Research 18, 1944–1954 (2008).

42. Wang, S. et al. EndHiC: assemble large contigs into chromosome-level scaffolds using the Hi-C links from contig ends. BMC Bioinformatics 23, 528 (2022).

43. Guan, D. et al. Efficient iterative Hi-C scaffolder based on N-best neighbors. BMC Bioinformatics 22, 569 (2021).

44. Chen, S., Zhou, Y., Chen, Y. & Gu, J. fastp: an ultra-fast all-in-one FASTQ preprocessor. *Bioinformatics (Oxford*, England) 34, i884–i890 (2018).

45. Durand, N.C. et al. Juicer provides a one-click system for analyzing loop-resolution Hi-C experiments. Cell systems 3, 95–98 (2016).

46. Li, H. Aligning sequence reads, clone sequences and assembly contigs with BWA-MEM. *arXiv preprint arXiv:1303.*3997 (2013).

47. Faust, G.G. & Hall, I.M. SAMBLASTER: fast duplicate marking and structural variant read extraction. *Bioinformatics (Oxford*, England) 30, 2503–2505 (2014).

48. Li, H. et al. The Sequence Alignment/Map format and SAMtools. *Bioinformatics (Oxford*, England) 25, 2078–2079 (2009).

49. DeMaere, M.Z. & Darling, A.E. Sim3C: simulation of Hi-C and Meta3C proximity ligation sequencing technologies. GigaScience 7 (2017).

50. Pecrix, Y. et al. Whole-genome landscape of *Medicago truncatula* symbiotic genes. Nature Plants 4, 1017–1025 (2018).

51. Tange, O. GNU Parallel 2018. Zenodo 10.5281/zenodo.1146014 (2018).

52. Wu, T.D. & Watanabe, C.K. GMAP: a genomic mapping and alignment program for mRNA and EST sequences. *Bioinformatics (Oxford*, England) 21, 1859–1875 (2005).

53. Li, H. Minimap2: pairwise alignment for nucleotide sequences. *Bioinformatics (Oxford*, England) 34, 3094–3100 (2018).

54. Goel, M., Sun, H., Jiao, W.-B. & Schneeberger, K. SyRI: finding genomic rearrangements and local sequence differences from whole-genome assemblies. Genome Biology 20, 277 (2019).

55. Goel, M. & Schneeberger, K. plotsr: visualizing structural similarities and rearrangements between multiple genomes. *Bioinformatics (Oxford*, England) 38, 2922–2926 (2022).

