## Supplementary Information for "Chromosome-level scaffolding of haplotype-resolved assemblies using Hi-C data without reference genomes"

### Software Commands

#### 1 Simulation of datasets for evaluating scaffolding tools

##### 1.1 Custom scripts

All custom Python scripts used in this section are available in the "simulation" directory of the HapHiC GitHub repository (https://github.com/zengxiaofei/HapHiC).

##### 1.2 Simulation of assemblies with different N50 values and coefficients of variation

We used a custom script, "sim_contig.py", to simulate assemblies with different N50 values and coefficients of variation (CVs) of length based on the chromosome-level genome of ground truth (chr_genome.fa):

```
$ python3 sim_contigs.py <chr_genome.fa> <N50> <CV>
```

##### 1.3 Simulation of Hi-C data with different effective sequencing depths

We used SAMtools (version 1.11) for random sampling to simulate Hi-C data with different effective sequencing depths. The parameter "-s" sets the seed and the fraction of reads to be sampled. For example, we used "-s 12345.092" to achieve random sampling of 9.2% of filtered reads with a seed of 12345, resulting in ~1X of effective Hi-C data:

```
$ samtools view <filtered_bam> -s 12345.092 -o <filtered_bam_1X>
```

Similarly, we used a custom script "sample_mnd.py" to randomly sample Hi-C reads from "merged_nodups.txt" file for 3D-DNA:

```
$ python3 sample_mnd.py <merged_nodups_mapq1.txt> <npairs> <fraction> >
<merged_nodups_mapq1_sampled.txt>
```

##### 1.4 Simulation of assemblies with different proportions of chimeric contigs

We used a custom script, "sim_chimeric_contigs.py", to simulate assemblies with different proportions of chimeric contigs based on the contig-level genome assemblies (asm.fa):

```
$ python3 sim_chimeric_contigs.py <asm.fa> --chimeric_ratio <ratio> --
inner_chrom_weight 0.1 --inter_nonhomo_weight 0.2 --inter_homo_weight
0.7
```

where chimeric contigs were generated between homologous chromosomes, non-homologous chromosomes, and within chromosomes at a ratio of 7:2:1, respectively.

##### 1.5 Simulation of datasets with different sequence divergences and ploidies

We used a custom script, "sim_haplotypes.py", to simulate haplotypes with different sequence divergences and ploidies based on the chromosome-level genome of the first haplotype in the ground truth (first_hap.fa):

```
$ python3 sim_haplotypes.py <first_hap.fa> <div> <ploidy>
```

By default, SNPs, insertions, and deletions were generated at a ratio of 10:0.5:0.5, with a 2:1 ratio of transitions to transversions for SNPs. The command produced a FASTA file of chromosome-level haplotypes (haps.fa) and a file containing allele information (allele_info.txt). Contig-level assemblies can then be simulated based on these haplotypes using "sim_contigs.py", as described in Section **1.2**.

The corresponding Hi-C reads were generated by sim3C:

```
$ sim3C <haps.fa> <sim.fq> --profile ./mycom.txt -n <npairs> --
efficiency 0.9 --trans-rate <rate> --anti-rate 0 -l 150 -e MboI -m hic
--seed <seed> --machine-profile HiSeqXtruSeqL150
```

where "mycom.txt" is the community profile file indicating that all chromosomes are located in the same cell with equal abundance and copy numbers set to 1. The parameters "-n" and "--trans-rate" were dynamically adjusted according to different ploidies to maintain a fixed intensity of intra- and inter-chromosome Hi-C links, as listed in Supplementary Table 9. To save time, the command was executed using ten parallel jobs with different seeds.

##### 1.6 Simulation of assemblies with different proportions of collapsed contigs

We used a custom script, "sim_collapsed_regions.py", to simulate different proportions and lengths of collapsed regions. This was performed based on the chromosome-level genome of the first haplotype in the ground truth (first_hap.fa) and the allele information file (allele_info.txt):

```
$ python3 sim_collapsed_regions.py <first_hap.fa> <allele_info.txt> --
collapsed_len <len> --collapsed_ratio <ratio>
```

By default, collapsed regions formed by two, three, and four haplotypes were generated in a ratio of 7:2:1. This command produced a FASTA file as the template and another FASTA file containing simulated collapsed regions. The latter file was subsequently fragmented into contig-level assemblies using "sim_contigs.py", as described in Section **1.2**.

##### 1.7 Simulation of assemblies with different switch error rates

We used a custom script, "sim_switch_errors.py", to simulate haplotypes with different switch error rates. This was performed based on the chromosome-level genome of the first haplotype in the ground truth (first_hap.fa) and the allele information file (allele_info.txt):

```
$ python3 sim_switch_errors.py <first_hap.fa> <allele_info.txt> --rate
<rate>
```

This command randomly shuffled variations between haplotypes and generated a new allele information file (new_allele_info.txt). The resulting FASTA files were subsequently fragmented into contig-level assemblies using “sim_contigs.py”, as described in Section **1.2**.

##### 1.8 Simulation of assemblies with different ploidies and three types of assembly errors

We have developed a pipeline that incorporates custom scripts to introduce 5% each of chimeric, collapsed contigs and switch errors in assemblies with different ploidies.

(1) The script “sim_switch_error.py” was used to simulate haplotypes with a 5% switch error rate:

```
# simulate switch errors and generate new_allele_info.txt
$ python3 sim_switch_errors.py <first_hap.fa> <allele_info.txt> --rate
0.05
```

(2) Based on the newly generated allele information file, we simulated 5% of collapsed contigs using “sim_collapsed_regions.py”:

```
# simulate collapsed regions and generate a FASTA file as the template
and a FASTA files (template.fa) containing simulated collapsed regions
(collapsed.fa)
$ python3 sim_collapsed_regions.py <first_hap.fa> <new_allele_info.txt>
# fragment collapsed.fa into contig-level assemblies (ctg_asm.fa)
$ python3 sim_contigs.py <collapsed.fa> 500000 0.2
```

(3) The script “sim_chimeric_contigs.py” was used to generate 5% of chimeric contigs in the fragmented assemblies:

```
# simulate chimeric contigs
$ python3 sim_chimeric_contigs.py <ctg_asm.fa> --chimeric_ratio 0.05 --
inner_chrom_weight 0.1 --inter_nonhomo_weight 0.2 --inter_homo_weight
0.7
```

(4) Corresponding Hi-C reads were simulated by sim3C, based on the template FASTA file generated in step (2):

```
# simulate Hi-C reads based on template.fa
$ sim3C <template.fa> <sim.fq> --profile ./mycom.txt -n <npairs> --
efficiency 0.9 --trans-rate <rate> --anti-rate 0 -l 150 -e MboI -m hic
--seed <seed> --machine-profile HiSeqXtruSeqL150
```

##### 1.9 Simulation of clusters for evaluating the reassignment step in HapHiC

We used a script, “sim_for_reassignment.py”, to simulate a series of clusters with different misassignment rates, anchoring rates, and contiguity levels. The commands are as follows:

```
# (1) simulate misassignments between homologous chromosomes
$ python3 sim_for_reassignment.py <asm.fa> <rate> --error_type
inter_homo --output_groups
```

```
# (2) simulate misassignments between non-homologous chromosomes
$ python3 sim_for_reassignment.py <asm.fa> <rate> --error_type
inter_nonhomo --output_groups
```

```
# (3) simulate different contiguity levels
$ python3 sim_for_reassignment.py <asm.fa> <contiguity> --error_type
contiguity --output_groups
```

```
# (4) simulate different anchoring rates
$ python3 sim_for_reassignment.py <asm.fa> <rate> --error_type
anchoring_rate --output_groups
```

##### 1.10 Simulation of datasets for evaluating contig ordering and orientation in each scaffolding tool

We performed ordering and orientation on each chromosome in the reference genomes of rice (*Oryza sativa*, IRGSP-1.0), *Arabidopsis* (*Arabidopsis thaliana*, TAIR10.1), and human (*Homo sapiens*, CHM13v2.0_noY). To achieve this, we need to prepare chromosome-specific genome assemblies and mapped Hi-C reads for each scaffolding tool.

These chromosome-level genomes were fragmented into contig-level assemblies (asm.fa) using “sim_contigs.py”, as described in Section **1.2**. Chromosome-specific contigs were extracted based on sequence IDs, which had recorded their source chromosomes.

For 3D-DNA, we split “merged_nodups.txt” files containing mapped Hi-C reads into chromosome-specific subfiles using “split_mnd.py”:

```
$ python3 split_mnd.py <merged_nodups.txt> Chr1 Chr2 Chr3 Chr4 Chr5
[chrn ...]
```

For LACHESIS, SALSA2, and YaHS, we split BAM files containing mapped Hi-C reads into chromosome-specific subfiles using “split_bam.py”:

```
$ python3 split_bam.py <filtered_HiC.bam> Chr1 Chr2 Chr3 Chr4 Chr5
[chrn ...]
```

ALLHiC and HapHiC record mapped Hi-C reads using CLM files. ALLHiC converts all mapped Hi-C reads into a single CLM file, while HapHiC splits the CLM file into chromosome-specific subfiles. We used HapHiC to prepare both the single CLM file and chromosome-specific subfiles for HapHiC and ALLHiC:

```
$ haphic cluster <asm.fa> <filtered_HiC.bam> <nchrs> --skip_clustering
```

In addition, ALLHiC and HapHiC perform ordering and orientation on each cluster (chromosome). Therefore, we simulated chromosome-specific clusters (referred to as "counts_RE.txt" in ALLHiC) using a script "sim_group_files.py" for these two scaffolding tools:

```
$ python3 sim_group_files.py <asm.fa>
```

##### 1.11 Simulation of contigs for published genomes (real cases)

For published genomes, we generated contig-level assemblies by breaking the "N" gaps and randomly shuffling the ordering and orientation of contigs using custom scripts, namely "fasta_split.py" and "shuffle_fasta.py". The commands are as follows:

```
# get contig-level assembly (asm.fa) by breaking the "N" gaps in
chromosome-level reference genome (ref.fa)
$ python3 split_fasta.py <ref.fa>

# shuffle the ordering and orientation of contigs in asm.fa
$ python3 shuffle_fasta.py <asm.fa>
```

#### 2 Quality control, mapping and filtering of Hi-C reads

##### 2.1 Quality control of Hi-C reads

We used fastp (version 0.21.0) to remove adaptors, trim low-quality sequences, and evaluate the quality of raw reads for each Hi-C library:

```
$ fastp -i <raw_1.fq> -I <raw_2.fq> -o <1.fq> -O <2.fq> --threads 28 --
json <libraryID.json> --html <libraryID.html>
```

##### 2.2 General purpose Hi-C read mapping and filtering

We used BWA-MEM (version 0.7.17-r1198-dirty) with the "-5SP" parameter to align Hi-C reads for general purpose. PCR duplicates were identified and marked using samblaster (version 0.1.26). Secondary and supplementary alignments were filtered out using SAMtools (version 1.11) with the flag "3340". The commands are as follows:

```
$ bwa index <asm.fa>
$ bwa mem -5SP <asm.fa> <1.fq> <2.fq> -t 28 | samblaster | samtools
view -@ 14 -S -h -b -F 3340 -o <HiC.bam>
```

Then, we removed the alignments with a mapping quality of zero (MAPQ = 0) and an edit distance greater than 2 (NM>=3) using a custom script "filter_bam.py", which is available in the "utils" directory of the HapHiC GitHub repository (https://github.com/zengxiaofei/HapHiC).

```
$ python3 filter_bam.py <HiC.bam> 1 --NM 3 --threads 14 | samtools view
-b -@ 14 -o <filtered_HiC.bam>
```

##### 2.3 Hi-C read mapping and filtering for 3D-DNA scaffolding

We used the Juicer pipeline (https://github.com/aidenlab/juicer, commit: 827fe44) to align and filter Hi-C reads for 3D-DNA as follows:

```
# (1) prepare scripts
ln -s /path/to/juicer2/CPU scripts

# (2) prepare reads
mkdir fastq && cd fastq
ln -s <1.fq> reads_R1.fastq.gz && ln -s <2.fq> reads_R2.fastq.gz &&
cd ..

# (3) prepare genome
mkdir references && cd references
ln -s <asm.fa> genome.fa && bwa index genome.fa && cd ..

# (4) prepare restriction sites
$ python2 /path/to/juicer2/misc/generate_site_positions.py MboI
genome ../references/genome.fa
$ awk 'BEGIN{OFS="\t"}{print $1, $NF}' genome_MboI.txt >
genome.chrom.sizes

# (5) run Juicer pipeline
$ bash /path/to/juicer2/CPU/juicer.sh -g genome -z references/genome.fa
-p restriction_sites/genome.chrom.sizes -y
restriction_sites/genome_MboI.txt -D `pwd` -s MboI -t 28 --assembly
```

#### 3 Hi-C-based scaffolding

##### 3.1 HapHiC

(1) HapHiC was typically executed with default optional parameters:

```
$ haphic pipeline <asm.fa> <filtered_HiC.bam> <nchrs> --RE <RE>
```

where the parameter “--RE” was set based on the restriction site used in Hi-C library construction. If the restriction site was unclear, the parameter was left as default (GATC). Commands for special cases are listed separately as follows.

(2) We evaluated the performance of HapHiC in scaffolding assemblies with contig N50 values ranging from 2 Mb to 25 Kb. For scaffolding the assembly with an extremely low contig N50 of 25 Kb, we used a lower contig length threshold of N70 for Markov clustering to save time and set lower thresholds for number of Hi-C links and link density to optimize the anchoring rate:

```
$ haphic pipeline <asm.fa> <filtered_HiC.bam> <nchrs> --Nx 70 --
min_link_density 0.00002 --min_links 1
```

(3) For scaffolding assemblies with different effective Hi-C sequencing depths, we used

lower thresholds for the number of Hi-C links and link density to optimize the anchoring rate. For effective depths ranging from 1X to 0.02X, the thresholds for number of Hi-C links were set to 20, 10, 4, 2, 1, 1, and 1, while the thresholds for Hi-C link density were set to 0.0001, 0.00005, 0.00002, 0.00001, 0.000005, 0.000003, and 0.000001 (which is 0.1 to 0.002 times the default values):

```
$ haphic pipeline <asm.fa> <filtered_HiC.bam> <nchrs> --
min_link_density <density> --min_links <links>
```

(4) For scaffolding assemblies with simulated chimeric contigs, HapHiC was executed with an additional parameter, "--correct_nrounds", which is required for assembly correction:

```
$ haphic pipeline <asm.fa> <filtered_HiC.bam> <nchrs> --correct_nrounds
2
```

(5) For scaffolding assemblies with simulated switch errors, HapHiC was executed with an additional parameter, "--remove_allelic_links", which is required for removing inter-allele Hi-C links:

```
$ haphic pipeline <asm.fa> <filtered_HiC.bam> <nchrs> --
remove_allelic_links <nhaps>
```

(6) To evaluate the performance of the rank-sum method in identifying chimeric or collapsed contigs, we calculated the rank-sum value of each contig using HapHiC:

```
$ haphic cluster <asm.fa> <filtered_HiC.bam> <nchrs> --Nx 100 --
bin_size 0 --min_RE_sites 0 --density_lower 0 --density_upper 1 --
rank_sum_upper 0 --skip_clustering --verbose 2> <log.txt>
```

(7) To evaluate the performance of the Hi-C link density method in identifying collapsed contigs, we calculated the link density of each contig using HapHiC:

```
$ haphic cluster <asm.fa> <filtered_HiC.bam> <nchrs> --Nx 100 --
bin_size 0 --min_RE_sites 0 --density_lower 0 --density_upper 0 --
rank_sum_upper 1 --skip_clustering --verbose 2> <log.txt>
```

(8) To evaluate the performance of the concordance ratio method in identifying allelic contig pairs, we calculated the concordance ratio of each contig using HapHiC:

```
$ haphic cluster <asm.fa> <filtered_HiC.bam> <nchrs> --Nx 100 --
bin_size 0 --min_RE_sites 0 --density_lower 0 --density_upper 1 --
rank_sum_upper 1 --remove_allelic_links 4 --skip_clustering --verbose
2> <log.txt>
```

(9) To evaluate the performance of reassignment, we executed HapHiC using the simulated clusters as described in Section **1.9**:

```
$ haphic reassign <asm.fa> <full_links.pkl> <simulated_clusters.txt> --
nclusters <nchrs>
```

(10) To evaluate the performance and efficiency of ordering and orientation, we executed HapHiC using the simulated datasets as described in Section **1.10**:

```
$ haphic sort <asm_chr.fa> <HT_links.pkl> <split_clm_dir>
<group_chr.txt>
```

where "asm_chr.fa" and "group_chr.txt" represent the chromosome-specific contigs and clusters, respectively. "HT_links.pkl" is a pickle file containing Hi-C links between semi-contigs, while "split_clm_dir" is a directory containing chromosome-specific CLM files. Both are generated by "haphic cluster". This command generates "*.tour.sav" files as the results of fast sorting and "*.tour" files as the final ordering and orientation results after ALLHiC optimization.

To evaluate the efficiency (execution speed and memory usage) of the entire HapHiC pipeline without ALLHiC optimization (fast sorting only):

```
$ haphic pipeline <asm.fa> <filtered_HiC.bam> <nchrs> --skip_allhic
```

(11) Given the extremely low contig contiguity of the *Saccharum spontaneum* AP85-441 genome assembly (contig N50 of ~45 Kb) and the presence of numerous collapsed contigs, we added the parameters "–Nx 70", "–density_upper 0.9 --rank_sum_upper 0.8", and "–min_RE_sites 2" to save time, filter out potential collapsed contigs, and optimize the anchoring rate, respectively:

```
$ haphic pipeline <asm.fa> <filtered_HiC.bam> 32 --RE AAGCTT --Nx 70 --
density_upper 0.9 --rank_sum_upper 0.8 --min_RE_sites 2
```

(12) For scaffolding the genome assembly of *Medicago sativa* XinJiangDaYe, we added the parameters "--min_links 5 --min_link_density 0.00001" to optimize the anchoring rate:

```
$ haphic pipeline <asm.fa> <filtered_HiC.bam> 32 --min_links 5 --
min_link_density 0.00001
```

(13) Given the extremely low contig contiguity of the *M. sativa* Zhongmu-4 genome assembly (contig N50 of ~94 Kb) and the presence of numerous collapsed contigs, we added the parameters "–Nx 70", "–density_upper 0.8 --rank_sum_upper 0.8", and "–min_RE_sites 2 --min_links 5" to save time, filter out potential collapsed contigs, and optimize the anchoring rate, respectively:

```
$ haphic pipeline <asm.fa> <filtered_HiC.bam> 32 --RE AAGCTT --Nx 70 --
density_upper 0.8 --rank_sum_upper 0.8 --min_RE_sites 2 --min_links 5
```

(14) Considering the presence of numerous collapsed and chimeric contigs in the genome assembly of *S. spontaneum* Np-X, we added "--correct_nrounds 2" to correct chimeric contigs and "--density_upper 0.9 --rank_sum_upper 0.8" for filtering of collapsed contigs:

```
$ haphic pipeline <asm.fa> <filtered_HiC.bam> 40 --RE AAGCTT --
correct_nrounds 2 --density_upper 0.9 --rank_sum_upper 0.8
```

(15) Considering the presence of large, nearly identical regions in the potato C88 genome where Hi-C links are not evenly distributed. we added "--normalize_by_nlinks --remove_concentrated_links" to normalize the Hi-C links between contigs:

```
$ haphic pipeline <asm.fa> <filtered_HiC.bam> 48 --RE AAGCTT --
normalize_by_nlinks --remove_concentrated_links
```

##### 3.2 ALLHiC

(1) We evaluated the performance of ALLHiC (version 0.9.13) in scaffolding assemblies under various adverse conditions. The chromosome assignment in ALLHiC was typically performed in three different modes with default parameters.

(i) Using a reference genome for pruning and separating homologous groups (referred to as "ALLHiC sep" in figures):

```
# generate Allele.ctg.table for pruning
$ gmap_build -D . -d DB <asm.fa>
# use gmapl instead of gmap for large genomes
$ gmap -D . -d DB -t 28 -f 2 -n <nhaps> <cds.fa> > <gmap.gff3>
$ perl /path/to/ALLHiC/scripts/gmap2AlleleTable.pl <gmap.gff3>

# separate homologous groups
$ samtools sort <filtered_HiC.bam> -o <sorted_filtered_HiC.bam> -@ 28
$ samtools index <sorted_filtered_HiC.bam>
$ perl /path/to/ALLHiC/scripts/partition_gmap.py -r <asm.fa> -g
Allele.ctg.table -b <sorted_filtered_HiC.bam> -t 28

# pruning for each separated homologous group
$ ALLHiC_prune -i <Allele.ctg.table> -b <sorted_filtered_HiC_group.bam>
-r <asm_group.fa>

# partitioning (clustering) for each separated homologous group
$ ALLHiC_partition -b <pruning.bam> -r <asm_group.fa> -e <RE> -k
<nclusters>

# rescue for each separated homologous group
$ ALLHiC_rescue -b <sorted_filtered_HiC_group.bam> -r <asm_group.fa> -c
<clusters_group.txt> -i <counts_RE_group.txt>

# global rescue (when anchoring rate is less than 1)
# generate counts_RE.txt file for all contigs (counts_RE_all.txt)
$ allhic extract <filtered_HiC_group.bam> <asm.fa> --RE <RE>
$ ALLHiC_rescue -r <asm.fa> -b <filtered_HiC_group.bam> -c
<clusters_all.txt> -i <counts_RE_all.txt>
```

In the scaffolding of autotetraploid genome assemblies for *M. sativa* (including XinJiangDaYe, Zhongmu-4, and simulated assemblies derived from XinJiangDaYe), the genome of a closely related species, *Medicago truncatula* (MtrunA17r5.0-ANR51), was used as a reference. Considering that chromosomes 4 and 8 have structural differences compared to those in *M. sativa*, the parameter "nclusters" was set to 8 for these chromosomes during the partitioning step. For the rest of the chromosomes, this parameter was set to 4.

(ii) Using a reference genome for pruning without separating homologous groups (referred to as "ALLHiC" in figures):

```
# generate Allele.ctg.table for pruning
$ gmap_build -D . -d DB <asm.fa>
# use gmapl instead of gmap for large genomes
$ gmap -D . -d DB -t 28 -f 2 -n <nhaps> <cds.fa> > <gmap.gff3>
$ perl /path/to/ALLHiC/scripts/gmap2AlleleTable.pl <gmap.gff3>

# pruning
$ ALLHiC_prune -i <Allele.ctg.table> -b <filtered_HiC.bam> -r <asm.fa>

# partitioning (clustering)
$ ALLHiC_partition -b <pruning.bam> -r <asm.fa> -e <RE> -k <nclusters>

# rescue (when anchoring rate is less than 1)
$ ALLHiC_rescue -b <filtered_HiC.bam> -r <asm.fa> -c
<pruning_clusters.txt> -i <pruning_counts_RE.txt>
```

(iii) Executing ALLHiC without pruning or separating homologous groups (referred to as "ALLHiC no pruning" in figures):

```
# partitioning (clustering)
$ ALLHiC_partition -b <filtered_HiC.bam> -r <asm.fa> -e <RE> -k
<nclusters>

# rescue (when anchoring rate is less than 1)
$ ALLHiC_rescue -b <filtered_HiC.bam> -r <asm.fa> -c <clusters_all.txt>
-i <counts_RE_all.txt>
```

(2) For the processes of ordering, orientation, and building final scaffolds:

```
# order and orient contigs for each cluster/chromosome
$ allhic optimize <counts_RE_chr.txt> <HiC.clm>

# build final scaffolds/pseudomolecules in the directory containing
*.tour files
$ ALLHiC_build <asm.fa>
```

(3) For scaffolding assemblies with different effective Hi-C sequencing depths, we used lower thresholds for link density to optimize anchoring rate. For effective depths ranging from 1X to 0.02X, the thresholds for Hi-C link density were set to 0.0001, 0.0005, 0.0002, 0.0001, 0.00005, 0.00003, and 0.00002 in the rescue process (which is 0.1 to 0.002 times the default values):

```
# (i) Using a reference genome for pruning and separating homologous
groups
```

```
# rescue for each separated homologous group
$ ALLHiC_rescue -r <asm.fa> -b <filtered_HiC_group.bam> -c
<clusters_all.txt> -i <counts_RE_all.txt> -m <density>
```

```
# global rescue
$ ALLHiC_rescue -r <asm.fa> -b <filtered_HiC_group.bam> -c
<clusters_all.txt> -i <counts_RE_all.txt> -m <density>
```

```
# (ii) Using a reference genome for pruning without separating
homologous groups
$ ALLHiC_rescue -b <filtered_HiC.bam> -r <asm.fa> -c
<pruning_clusters.txt> -i <pruning_counts_RE.txt> -m <density>
```

```
# (iii) Executing ALLHiC without pruning or separating homologous
groups
$ ALLHiC_rescue -b <filtered_HiC.bam> -r <asm.fa> -c <clusters_all.txt>
-i <counts_RE_all.txt> -m <density>
```

(4) When scaffolding assemblies with chimeric contigs, we performed additional assembly correction before the ALLHiC pipeline:

```
$ samtools sort <filtered_HiC.bam> -o <sorted_filtered_HiC.bam> -@ 28
$ samtools index <sorted_filtered_HiC.bam>
$ ALLHiC_corrector -m <sorted_filtered_HiC.bam> -r <asm.fa> -o
<asm_corrected.fa> -t 28
```

Subsequently, Hi-C reads were realigned to "asm_corrected.fa" using the method described in Section **2.2**.

(5) When scaffolding the genome assembly of *M. sativa* XinJiangDaYe, we added "-m 100" in the partitioning step to optimize the results:

```
$ ALLHiC_partition -b <pruning.bam> -r <asm_group.fa> -e <RE> -k
<nclusters> -m 100
```

##### 3.3 LACHESIS

We executed LACHESIS (https://github.com/shendurelab/LACHESIS, commit: 2e27abb) without using a reference genome:

```
$ Lachesis params.ini
```

where the "params.ini" file contained several modifications from the default "test_case.ini" file provided by LACHESIS:

```
# use an arbitrary species name but avoid using string like "human" to
represent a species without a reference genome
SPECIES = alfalfa
OUTPUT_DIR = out
DRAFT_ASSEMBLY_FASTA = <asm.fa>
SAM_DIR = ./bam
```

```
SAM_FILES = <filtered_HiC.bam>
RE_SITE_SEQ = <RE>
USE_REFERENCE = 0
CLUSTER_N = <nchrs>
CLUSTER_DRAW_HEATMAP = 0
CLUSTER_DRAW_DOTPLOT = 0
REPORT_DRAW_HEATMAP = 0
```

##### 3.4 SALSA2

SALSA2 (https://github.com/marbl/SALSA, commit: ed76685) was executed without an assembly graph.

(1) Assembly correction was typically disabled. Given that the BAM file "filtered_HiC.bam" is already sorted by name, the sorting step was skipped:

```
$ bedToBed -i <filtered_HiC.bam> > alignment.bed
$ samtools faidx <asm.fa>
$ python2 run_pipeline.py -a <asm.fa> -l <asm.fa.fai> -b alignment.bed
-e <RE> -m no -o output
```

(2) For scaffolding assemblies with chimeric contigs, assembly correction was enabled:

```
$ python2 run_pipeline.py -a <asm.fa> -l <asm.fa.fai> -b alignment.bed
-e <RE> -m yes -o output
```

##### 3.5 3D-DNA

(1) 3D-DNA (https://github.com/aidenlab/3d-dna, commit: 529ccf4) was typically executed using default parameters without assembly correction:

```
$ bash run-asm-pipeline.sh -r 0 <asm.fa> <merged_nodups.txt>
```

(2) For scaffolding assemblies with different proportions of chimeric contigs, we performed two rounds of assembly correction. The repeat coverage threshold for the misjoin editor was set to 6 to prevent error messages. For scaffolding assemblies with 5% each of chimeric, collapsed contigs, and switch errors, we set the repeat coverage threshold for the misjoin editor to 6 and 10 for diploid and triploid genomes, respectively:

```
$ bash run-asm-pipeline.sh -r 2 --editor-repeat-coverage <cov> <asm.fa>
<merged_nodups.txt>
```

(3) For scaffolding assemblies with low contig N50 values (50 Kb and 25 Kb), and high proportions of collapsed contigs and switch errors (25%), we set a lower coarse stringency value of 30 for the polisher to prevent error messages:

```
$ bash run-asm-pipeline.sh -r 0 --polisher-coarse-stringency 30
<asm.fa> <merged_nodups.txt>
```

##### 3.6 YaHS

(1) YaHS (https://github.com/c-zhou/yahs, commit: 42b8421) was typically executed without assembly correction and memory check:

```
$ samtools faidx <asm.fa>
$ yahs <asm.fa> <filtered_HiC.bam> -e <RE> -q 1 --no-contig-ec --no-mem-check
```

(2) When encountering difficulties in scaffolding low-contiguity assemblies with YaHS, we specified different ranges of resolutions. For example:

```
$ yahs <asm.fa> <filtered_HiC.bam> -e <RE> -q 1 --no-contig-ec --no-mem-check -r 1000,2000,5000,10000,20000,50000,100000,200000,500000,1000000,2000000,5000000,10000000,20000000,50000000,100000000,200000000,500000000
```

(3) Given that YaHS can only handle up to 45,000 contigs, we used the parameter "-l" to specify the minimum contig length for scaffolding when the number of contigs exceeded this limit. For example:

```
$ yahs <asm.fa> <filtered_HiC.bam> -e <RE> -q 1 --no-contig-ec -l 5000 --no-mem-check
```

(4) For scaffolding assemblies with chimeric contigs, assembly correction was enabled:

```
$ yahs <asm.fa> <filtered_HiC.bam> -e <RE> -q 1 --no-mem-check
```

(5) In case of an out-of-memory error, we manually released the cached memory and then executed YaHS without the parameter "--no-mem-check":

```
$ sync && echo 1 > /proc/sys/vm/drop_caches
$ yahs <asm.fa> <filtered_HiC.bam> -e <RE> -q 1 --no-contig-ec
```

##### 3.7 Visualization in Juicebox

To visualize the scaffolding results of HapHiC, ALLHiC, and YaHS in Juicebox, we prepared "*.assembly" and "*.hic" files based on the AGP files (*.agp) output by these scaffolding tools using matlock (https://github.com/phasegenomics/matlock), juicebox scripts (https://github.com/phasegenomics/juicebox_scripts, commit: 4d3d297) and 3D-DNA (https://github.com/aidenlab/3d-dna, commit: 529ccf4):

```
# (1) generate .mnd files (merged nodups)
$ /path/to/matlock bam2 juicer <filtered_HiC.bam> > out.links.mnd
$ sort -k2,2 -k6,6 out.links.mnd > out.sorted.links.mnd

# (2) generate .assembly file
$ /path/to/juicebox_scripts/agp2assembly.py <scaffolds_final.agp> <scaffolds_final.assembly>

# (3) generate .hic file
$ bash /path/to/3d-dna/visualize/run-assembly-visualizer.sh -p false <scaffolds_final.assembly> out.sorted.links.mnd
```

Subsequently, the generated "*.hic" and "*.assembly" files were imported into Juicebox (version 1.11.08) for visualization.

#### 4 Performance evaluation of scaffolding

##### 4.1 Custom scripts

All custom Python scripts used in this section are available in the "simulation" directory of the HapHiC GitHub repository (https://github.com/zengxiaofei/HapHiC).

##### 4.2 Performance evaluation of chromosome assignment

HapHiC and ALLHiC provide the results of chromosome assignment in "group*.txt" files (also known as "counts_RE.txt" in ALLHiC). For other scaffolding tools, we converted their results to this format using different custom scripts:

```
# (1) for LACHESIS
$ python3 convert_lachesis_result_to_groups.py <clusters.by_name.txt>
<asm.fa>
```

```
# (2) for 3D-DNA
$ python3 convert_assembly_to_groups.py <genome.final.assembly>
```

```
# (3) for SALSA2 and YaHS
$ python3 convert_agp_to_groups.py <scaffolds_final.agp>
```

Subsequently, we evaluated the performance of Hi-C-based scaffolding tools in chromosome assignment using metrics such as contiguity, anchoring rate, misassignment rates, and the number of scaffolds. This was accomplished using a custom script named "result_statistics.py":

```
$ python3 result_statistics.py <asm.fa> <counts_RE_group1.txt>
<counts_RE_group2.txt> [<counts_RE_group3.txt> ...]
```

##### 4.3 Performance evaluation of ordering and orientation

HapHiC and ALLHiC provide the results of ordering and orientation in "*.tour" files. For other scaffolding tools, we converted their results to this format using different custom scripts:

```
# (1) for LACHESIS
$ python3 convert_lachesis_ordering_to_tour.py <asm_chr.fa> <chr_name>
<group0.ordering> <group1.ordering> [<group2.ordering> ...]
```

```
# (2) for 3D-DNA
$ python3 convert_assembly_to_tour.py <genome.0.assembly> <chr_name>
```

```
# (3) for SALSA2 and YaHS
$ python3 convert_agp_to_tour.py <scaffolds_final.agp> <chr_name>
```

Subsequently, the ordering and orientation results of each scaffolding tools were visualized using a custom script, “draw_tour_file.py”. This script calculates Lin’s concordance correlation coefficients (CCCs) and DERANGE costs for each input “*.tour” file:

```
$ python3 draw_tour_file.py <asm_chr.fa> <chr.tour> <program_name>
<N50> --CCC --derange
```

#### 4.4 Performance evaluation of assembly correction

We evaluated the performance of assembly correction on simulated chimeric contigs for each scaffolding tool. The breakpoints were extracted and analyzed using different custom scripts:

```
# (1) for HapHiC
$ python3 get_haphic_break_points.py <asm.fa> <asm_corrected.fa> <N50>

# (2) for ALLHiC
$ python3 get_allhic_break_points.py <asm.fa> <asm_corrected.fa> <N50>

# (3) for 3D-DNA
$ python3 get_3d_dna_break_points.py <asm.fa> <genome.FINAL.assembly>
<N50>

# (4) for SALSA2
$ python3 get_salsa_break_points.py <asm.fa> <scaffolds_final.agp>

# (5) for YaHS
$ python3 get_yahs_break_points.py <asm.fa> <scaffolds_final.agp>
```

We also compared the performance of assembly correction between HapHiC and ALLHiC in the genome assembly of *S. spontaneum* Np-X:

```
$ python3 correction_analysis.py <asm.fa> <filtered_HiC.bam>
<ctg_annoation_list.txt> <scaffolds_HapHiC.agp>
<scaffolds_ALLHiC.agp> > correction_statistics.txt
```

where “ctg_annoation_list.txt” is a list file that contains manually annotated chimeric and non-chimeric contigs.

#### 4.5 Performance evaluation of the rank-sum method in identifying chimeric and collapsed contigs

We evaluated the performance of the rank-sum method in identifying chimeric and collapsed contigs using the “log.txt” files generated in Section **3.1** (6). The custom scripts “chimeric_contig_statistics.py” and “collapsed_contig_statistics.py” were utilized for this purpose:

```
# (1) for chimeric contigs
$ python3 chimeric_contig_statistics.py <asm.fa> <log.txt>
<program_name> <N50>

# (2) for collapsed contigs
```

```
$ python3 collapsed_contig_statistics.py <asm.fa> <log.txt>
<program_name> <N50> --method rank_sum
```

##### 4.6 Performance evaluation of the Hi-C link density method in identifying collapsed contigs

We evaluated the performance of the Hi-C link density method in identifying collapsed contigs using the “log.txt” files generated in Section **3.1** (7). The custom script “collapsed_contig_statistics.py” was utilized for this purpose:

```
$ python3 collapsed_contig_statistics.py <asm.fa> <log.txt>
<program_name> <N50> --method link_density
```

##### 4.7 Performance evaluation of the concordance ratio method and the pruning method in identifying allelic contig pairs

We evaluated the performance of the concordance ratio method and pruning method in identifying allelic contig pairs using the “log.txt” files generated in Section **3.1** (8). The custom script “allelic_contig_statistics.py” was utilized for this purpose:

```
$ python3 allelic_contig_statistics.py <log.txt> <Allele.ctg.table>
<N50>
```

#### 5 Genome assembly

##### 5.1 Genome assembly of *Miscanthus* × *giganteus*

The genome of *M.* × *giganteus* was assembled using hifiasm (version 0.13-r308). The parameter “-l0” was employed to generate primary unitigs (p_utg):

```
$ hifiasm -o Mgi -l0 -t 28 <hifi.fq>
```

In addition, the genome of *M.* × *giganteus* was assembled using HiCanu (version 2.1.1) for comparison with the hifiasm assembly, in order to explain the source of diagonally distributed inter-allele Hi-C links:

```
$ canu -p Mgi -d Mgi_minOverlapLength_200 \
    genomeSize=2g \
    -pacbio-hifi <hifi.fq> \
    minOverlapLength=200
```

##### 5.2 Genome assembly of potato C88

The genome of potato (*Solanum tuberosum*) C88 was assembled using hifiasm (version 0.19.0-r534) to generate primary unitigs (p_utg):

```
$ hifiasm <hifi.fq> -l 0 --primary -t 64
```

#### 6 Comparative genomics analyses

##### 6.1 Custom scripts

All custom Python scripts used in this section are available in the "simulation" directory of the HapHiC GitHub repository (https://github.com/zengxiaofei/HapHiC).

##### 6.2 *K*-mer analysis

We performed a *k*-mer analysis on each potato C88 scaffold using a custom script, "haplotype_kmers.py", based on the published reference genome (C88.v1):

```
$ python3 haplotype_kmers.py <C88.v1.fa> <scaffold_*.fa>
```

where "scaffold_*.fa" represents the FASTA file of each scaffold and the default *k*-mer size was set to 201 bp.

##### 6.3 Gene synteny analysis

Initially, we simply identified coding sequences in each subgenome of *M.* × *giganteus* through GMAP (version 2019-12-01) mapping, using the coding sequences of *Miscanthus sinensis* as queries:

```
$ gmap_build -D . -d DB <subgenome.fa>
$ gmap -D . -d DB -t 28 -f 2 -n 1 ref.cds > <subgenome.gff>
# extract coding sequences using gffread from cufflinks (version 2.2.1)
$ gffread Msisub.gff -g genome.fa -x <subgenome.cds>
```

Subsequently, we performed gene synteny comparisons between the subgenomes of *Miscanthus* species and generated a karyotype plot using MCscan in JCVI utility libraries (version 1.1.18):

```
# prepare BED file for subgenomes
$ python3 -m jcvi.formats.gff bed --type=<type> --key=<key>
<subgenome.gff> -o <subgenome.bed>

# perform synteny analysis between subgenome pairs
$ python3 -m jcvi.compara.catalog ortholog <subgenomeA> <subgenomeB> --
no_strip_names --cscore 0.99
$ python3 -m jcvi.compara.synteny screen --minspan=30 --simple
subgenomeA.subgenomeB.anchors subgenomeA.subgenomeB.anchors.new

# retain synteny blocks between orthologous chromosomes
$ python3 filter_homologous_anchors.py
subgenomeA.subgenomeB.anchors.simple <subgenomeA.bed> <subgenomeB.bed>
"Chr01A,Chr02A,Chr03A,..." "Chr01B,Chr02B,Chr03B,..." >
subgenomeA.subgenomeB.anchors.simple.ortho

# draw karyotype plot
$ python3 -m jcvi.graphics.karyotype seqids layout
```

##### 6.4 Genome alignment and structural variation identification

(1) We performed a sequence alignment between potato C88 scaffolds and each haplotype of C88.v1 using unimap (https://github.com/lh3/unimap, version 0.1-r41):

```
# sequence alignment for each haplotype of C88.v1
$ unimap -d <hap*.umi> <C88.v1.hap*.fa> -t 28
$ unimap -c <hap*.umi> <scaffolds.fa> -x asm5 --cs -N 50 --secondary=no
-t 28 > <scaffolds_hap*.paf>
```

```
# concatenate all PAF files into a single file
$ cat scaffolds_hap1.paf scaffolds_hap2.paf scaffolds_hap3.paf
scaffolds_hap4.paf > scaffolds_allhaps.paf
```

Subsequently, we extracted the corresponding alignments from the PAF files for each chromosome and recalculate the number of matching bases (Column 10) based on the "de" tag (Column 21). For example:

```
$ grep "chr7_" ${aln} | egrep "group25|group28|group29|group30" | cut -
f 1-12,21 | sed 's/de:f://g' | awk '{print
$1"\t"$2"\t"$3"\t"$4"\t"$5"\t"$6"\t"$7"\t"$8"\t"$9"\t"(1-
$13)*$11"\t"$11"\t"$12}' > haphic_chr7.paf
```

Finally, we visualized the output PAF files using a modified version of paf2dotplot (https://github.com/zengxiaofei/paf2dotplot):

```
$ paf2dotplot.r -s -c 0.99 -p 4 <chr*.paf>
```

(2) We performed a sequence alignment and identified structural variations between chromosome 2 of *M.* × *giganteus* and *M. sinensis* using Minimap2 (version 2.26-r1175) and SyRI (version 1.6.3):

```
# sequence alignment
$ minimap2 -ax asm5 --eqx Msiref_Chr02.fa Msisub_Chr02.fa -t 28 >
Msisub_vs_Msiref_Chr02.sam
```

```
# structural variation identification
$ syri -c Msisub_vs_Msiref_Chr02.sam -r Msiref_Chr02.fa -q
Msisub_Chr02.fa -k -F S
```

```
# extract SVs identified by JCVI
$ python3 extract_SVs_from_simple.py Msi.Msisub.anchors.simple.ortho
Msi.bed
```

```
# visualization
$ plotsr --sr syri.out --genomes genomes.txt -s 100000 --tracks
tracks.txt -o plotsr_Msiref_ref.pdf -W 6 -H 3 -S 0.5
```

##### 6.5 Analysis of genetic maps

Initially, we aligned the genetic markers of each map to the A subgenomes of *M.* × *giganteus* (MgiA) and the *M. sinensis* genome (MsiA) using BWA-ALN and BWA-SAMSE (version 0.7.17-r1198-dirty) with default parameters:

```
$ bwa index <genome*.fa>
$ bwa aln <genome*.fa> <map*.fa> > <map*_genome*.sai> -t 28
```

```
$ bwa samse <genome*.fa> <map*_genome*.sai> <map*.fa> >
<map*_genome*.sam>
$ samtools view -b <map*_genome*.sam> -o <map*_genome*.bam>
```

After alignment, each pair of BAM files generated using the same genetic map were converted to a pair of CSV files, retaining only shared genetic markers mapped to the same linkage groups:

```
$ python3 bam_to_csv_shared.py --sam_LG_only <map*_genome_MgiA.bam>
<map*_genome_MsiA.bam>
```

Subsequently, we visualized the agreements between the genetic maps and each genome using ALLMAPS in JCVI utility libraries:

```
$ python3 -m jcvi.assembly.allmaps merge MapA_M_genome*.csv
MapA_P_genome*.csv MapB_M_genome*.csv MapB_P_genome*.csv
MapC_M_genome*.csv MapC_P_genome*.csv -o combined.bed
$ python3 mock_agp_file.py <genome*.fa> > combined.chr.agp
$ cp combined.chr.agp combined.agp
$ cp combined.bed combined.lifted.bed
$ python3 -m jcvi.assembly.allmaps plotall combined.bed
```

where the custom script “mock_agp_file.py” is available in the “utils” directory of the HapHiC GitHub repository at https://github.com/zengxiaofei/HapHiC.

#### Supplementary Tables

**Supplementary Table 1 |** Statistics of simulated assemblies with different N50 values

| Simulation (N50/CV) | Total length (bp) | Number of contigs | Max length (bp) | Avg length (bp) | Actual N50 (bp) |
|---|---|---|---|---|---|
| 2 Mb/0.2 | 2,044,810,672 | 1,028 | 3,571,011 | 1,989,115 | 2,064,317 |
| 1 Mb/0.2 | 2,044,810,672 | 2,049 | 1,785,505 | 997,955 | 1,037,798 |
| 800 Kb/0.2 | 2,044,810,672 | 2,558 | 1,350,484 | 799,379 | 832,577 |
| 500 Kb/0.2 | 2,044,810,672 | 4,084 | 892,752 | 500,688 | 520,506 |
| 300 Kb/0.2 | 2,044,810,672 | 6,826 | 535,651 | 299,562 | 311,687 |
| 200 Kb/0.2 | 2,044,810,672 | 10,256 | 357,101 | 199,377 | 207,685 |
| 100 Kb/0.2 | 2,044,810,672 | 20,470 | 178,550 | 99,893 | 103,997 |
| 50 Kb/0.2 | 2,044,810,672 | 40,928 | 89,617 | 49,961 | 52,025 |
| 25 Kb/0.2 | 2,044,810,672 | 81,856 | 45,829 | 24,981 | 25,978 |

**Supplementary Table 2 |** Statistics of simulated assemblies with different coefficients of variation (CVs)

| Simulation (N50/CV) | N10 (bp) | N30 (bp) | N50 (bp) | N70 (bp) | N90 (bp) |
|---|---|---|---|---|---|
| 500 Kb/0.2 | 647,202 | 573,137 | 520,506 | 466,757 | 397,161 |
| 500 Kb/0.3 | 670,526 | 572,006 | 534,413 | 429,894 | 333,633 |
| 500 Kb/0.4 | 716,123 | 595,095 | 507,578 | 417,377 | 301,438 |
| 500 Kb/0.6 | 760,920 | 611,098 | 500,829 | 397,033 | 255,800 |
| 500 Kb/0.8 | 819,645 | 637,888 | 512,337 | 398,322 | 241,392 |
| 500 Kb/1 | 821,080 | 630,147 | 499,628 | 381,094 | 224,603 |
| 500 Kb/1.5 | 848,800 | 636,684 | 501,865 | 373,211 | 213,587 |
| 500 Kb/3 | 868,450 | 641,236 | 498,918 | 362,597 | 206,077 |

**Supplementary Table 3 | Parameters for simulating different effective Hi-C sequencing depths**

| Sampling method | Simulation (depth) | Fraction | Number of read pairs | Number of read bases | Calculated depth |
|---|---|---|---|---|---|
| sample_mnd.py (For 3D-DNA) | All | NA | 97,500,525 | 19,692,884,346 | 9.6X |
| | 1X | 0.104 | 10,140,054 | 2,048,066,632 | 1.0X |
| | 0.5X | 0.0519 | 5,060,277 | 1,022,089,650 | 0.5X |
| | 0.2X | 0.0208 | 2,028,010 | 409,614,990 | 0.2X |
| | 0.1X | 0.0104 | 1,014,005 | 204,807,554 | 0.1X |
| | 0.05X | 0.00519 | 506,027 | 102,209,804 | 0.05X |
| | 0.03X | 0.00312 | 304,201 | 61,441,704 | 0.03X |
| | 0.02X | 0.00208 | 202,801 | 40,962,706 | 0.02X |
| SAMtools (For other scaffolding tools) | All | NA | 110,443,496 | 22,324,670,268 | 10.9X |
| | 1X | 0.092 | 10,161,564 | 2,054,060,928 | 1.0X |
| | 0.5X | 0. 0458 | 5,062,902 | 1,023,406,738 | 0.5X |
| | 0.2X | 0. 0183 | 2,023,015 | 408,930,034 | 0.2X |
| | 0.1X | 0. 00916 | 1,012,324 | 204,631,670 | 0.1X |
| | 0.05X | 0. 00458 | 506,602 | 102,407,568 | 0.05X |
| | 0.03X | 0. 00275 | 304,633 | 61,578,334 | 0.03X |
| | 0.02X | 0. 00183 | 202,516 | 40,937,902 | 0.02X |

**Supplementary Table 4 |** Statistics of simulated assemblies with different lengths and proportions of chimeric contigs

| Simulation (proportion/length) | Total length (bp) | Total length of chimeric contigs (bp) | Number of chimeric contigs | Avg length of chimeric contigs (Kb) | Calculated proportion |
|---|---|---|---|---|---|
| 20%/2 Mb | 2,044,810,672 | 400,417,814 | 204 | 1962.8 | 19.6% |
| 20%/1 Mb | 2,044,810,672 | 403,812,563 | 407 | 992.2 | 19.7% |
| 20%/800 Kb | 2,044,810,672 | 408,663,191 | 510 | 801.3 | 20.0% |
| 20%/500 Kb | 2,044,810,672 | 406,041,196 | 815 | 498.2 | 19.9% |
| 20%/300 Kb | 2,044,810,672 | 407,366,972 | 1,364 | 298.7 | 19.9% |
| 20%/200 Kb | 2,044,810,672 | 410,395,141 | 2,050 | 200.2 | 20.1% |
| 20%/100 Kb | 2,044,810,672 | 409,065,751 | 4,092 | 100.0 | 20.0% |
| 20%/50 Kb | 2,044,810,672 | 408,112,589 | 8,184 | 49.9 | 20.0% |
| 5%/500 Kb | 2,044,810,672 | 101,078,754 | 202 | 500.4 | 4.9% |
| 10%/500 Kb | 2,044,810,672 | 201,539,322 | 406 | 496.4 | 9.9% |
| 15%/500 Kb | 2,044,810,672 | 304,899,629 | 611 | 499.0 | 14.9% |
| 20%/500 Kb | 2,044,810,672 | 406,041,196 | 815 | 498.2 | 19.9% |
| 25%/500 Kb | 2,044,810,672 | 507,655,379 | 1,020 | 497.7 | 24.8% |
| 30%/500 Kb | 2,044,810,672 | 608,293,353 | 1,224 | 497.0 | 29.7% |
| 35%/500 Kb | 2,044,810,672 | 710,821,147 | 1,427 | 498.1 | 34.8% |
| 40%/500 Kb | 2,044,810,672 | 815,148,811 | 1,632 | 499.5 | 39.9% |

**Supplementary Table 5 |** Statistics of simulated assemblies with different proportions of collapsed contigs

| Simulation (proportion/length) | Total length (bp) | Total length of collapsed contigs (bp) | Number of collapsed contigs | Avg length of collapsed contigs (Kb) | Calculated proportion |
|---|---|---|---|---|---|
| 20%/2 Mb | 1,618,461,908 | 326,000,509 | 163 | 2000.0 | 20.1% |
| 20%/1 Mb | 1,636,461,918 | 327,000,255 | 328 | 997.0 | 20.0% |
| 20%/800 Kb | 1,639,262,554 | 327,200,663 | 410 | 798.1 | 20.0% |
| 20%/500 Kb | 1,636,461,582 | 327,500,111 | 658 | 497.7 | 20.0% |
| 20%/300 Kb | 1,639,162,248 | 327,600,641 | 1,095 | 299.2 | 20.0% |
| 20%/200 Kb | 1,642,462,159 | 327,799,923 | 1,640 | 199.9 | 20.0% |
| 20%/100 Kb | 1,642,161,390 | 327,800,153 | 3,285 | 100.0 | 20.0% |
| 20%/50 Kb | 1,641,511,968 | 327,850,417 | 6,570 | 49.9 | 20.0% |
| 5%/500 Kb | 1,957,462,001 | 97,999,972 | 198 | 495.0 | 5.0% |
| 10%/500 Kb | 1,837,461,910 | 184,000,064 | 370 | 497.3 | 10.0% |
| 15%/500 Kb | 1,733,962,036 | 260,000,265 | 523 | 497.1 | 15.0% |
| 17.5%/500 Kb | 1,685,961,813 | 294,499,854 | 589 | 500.0 | 17.5% |
| 20%/500 Kb | 1,636,461,582 | 327,500,111 | 658 | 497.7 | 20.0% |
| 22.5%/500 Kb | 1,594,462,108 | 359,000,004 | 718 | 500.0 | 22.5% |
| 25%/500 Kb | 1,553,461,670 | 388,500,153 | 777 | 500.0 | 25.0% |

**Supplementary Table 6 |** Statistics of simulated haplotypes at different sequence divergence levels

| Simulation (sequence divergence) | Calculated sequence divergence between haplotypes (number of variations divided by average length of two haplotypes) | | | | | |
|---|---|---|---|---|---|---|
| | Hap1 vs Hap2 | Hap1 vs Hap3 | Hap1 vs Hap4 | Hap2 vs Hap3 | Hap2 vs Hap4 | Hap3 vs Hap4 |
| 2% | 2.00% | 1.99% | 2.00% | 1.99% | 2.01% | 1.99% |
| 1.5% | 1.50% | 1.49% | 1.50% | 1.50% | 1.51% | 1.50% |
| 1% | 1.00% | 1.00% | 1.00% | 1.00% | 1.01% | 1.00% |
| 0.5% | 0.50% | 0.50% | 0.50% | 0.50% | 0.50% | 0.50% |
| 0.4% | 0.40% | 0.40% | 0.40% | 0.40% | 0.40% | 0.40% |
| 0.3% | 0.30% | 0.30% | 0.30% | 0.30% | 0.30% | 0.30% |
| 0.2% | 0.20% | 0.20% | 0.20% | 0.20% | 0.20% | 0.20% |
| 0.1% | 0.10% | 0.10% | 0.10% | 0.10% | 0.10% | 0.10% |

**Supplementary Table 7 |** Statistics of simulated haplotypes with different switch error rates

| Simulation (switch error rate) | Calculated switch error rate between haplotypes (number of switched variations divided by variations between two haplotypes) | | | | | | |
|---|---|---|---|---|---|---|---|
| | Hap1 vs Hap2 | Hap1 vs Hap3 | Hap1 vs Hap4 | Hap2 vs Hap3 | Hap2 vs Hap4 | Hap3 vs Hap4 | Total |
| 2.5% | 0.42% | 0.42% | 0.42% | 0.42% | 0.41% | 0.41% | 2.50% |
| 5% | 0.84% | 0.83% | 0.84% | 0.83% | 0.83% | 0.83% | 4.99% |
| 7.5% | 1.26% | 1.25% | 1.25% | 1.25% | 1.24% | 1.24% | 7.49% |
| 10% | 1.67% | 1.66% | 1.67% | 1.66% | 1.66% | 1.66% | 9.99% |
| 15% | 2.52% | 2.50% | 2.50% | 2.49% | 2.49% | 2.48% | 14.98% |
| 20% | 3.36% | 3.33% | 3.34% | 3.32% | 3.33% | 3.31% | 19.98% |
| 25% | 4.19% | 4.16% | 4.17% | 4.15% | 4.17% | 4.14% | 24.97% |

**Supplementary Table 8 |** Statistics of simulated assemblies at different ploidy levels

| Simulation (ploidy) | Total length (bp) | Number of contigs | Max length (bp) | Avg length (bp) | N50 (bp) |
|---|---|---|---|---|---|
| 1 | 524,615,222 | 1,051 | 892,752 | 499,158 | 515,805 |
| 2 | 1,049,230,598 | 2,099 | 892,752 | 499,872 | 519,610 |
| 3 | 1,573,846,741 | 3,144 | 892,752 | 500,587 | 522,203 |
| 4 | 2,098,461,825 | 4,194 | 892,752 | 500,349 | 520,439 |
| 6 | 3,147,691,937 | 6,304 | 892,752 | 499,317 | 519,824 |
| 8 | 4,196,923,741 | 8,414 | 920,319 | 498,802 | 520,056 |
| 10 | 5,246,154,432 | 10,530 | 920,345 | 498,210 | 519,137 |
| 12 | 6,295,384,748 | 12,626 | 957,377 | 498,605 | 519,888 |
| 16 | 8,393,845,927 | 16,833 | 957,294 | 498,654 | 519,914 |

**Supplementary Table 9 |** Sim3C parameters for simulating Hi-C reads of genomes at different ploidy levels

| Simulation (ploidy) | Number of read pairs | Trans rate |
|---|---|---|
| 1 | 87,500,000 | 0.104203152 |
| 2 | 175,000,000 | 0.199530516 |
| 3 | 262,500,000 | 0.276520509 |
| 4 | 350,000,000 | 0.34 |
| 6 | 525,000,000 | 0.438529089 |
| 8 | 700,000,000 | 0.511461318 |
| 10 | 875,000,000 | 0.567624683 |
| 12 | 1,050,000,000 | 0.612206217 |
| 16 | 1,400,000,000 | 0.678504085 |

**Supplementary Table 10 |** Statistics of simulated assemblies with different ploidies and 5% each of chimeric, collapsed, and switch errors

| Simulation (ploidy) | Total length (bp) | Number of contigs | Max length (bp) | Avg length (bp) | N50 (bp) |
|---|---|---|---|---|---|
| 2 | 999,730,885 | 2,121 | 917,743 | 471,349 | 506,006 |
| 3 | 1,481,346,483 | 3,168 | 892,752 | 467,597 | 500,023 |
| 4 | 1,957,461,928 | 4,247 | 952,262 | 460,905 | 500,008 |
| 6 | 2,938,691,961 | 6,584 | 946,643 | 446,338 | 500,002 |
| 8 | 3,919,423,846 | 9,071 | 913,634 | 432,083 | 499,999 |
| 10 | 4,902,653,840 | 12,067 | 944,770 | 406,286 | 499,996 |
| 12 | 5,883,384,546 | 15,449 | 1,005,068 | 380,826 | 499,995 |
| 16 | 7,852,345,231 | 24,867 | 579,692 | 315,773 | 499,991 |

**Supplementary Table 11 |** Statistics of simulated assemblies for rice (IRGSP-1.0), *Arabidopsis* (TAIR10.1), and human (CHM13v2.0_noY) with varying contig N50 values

| Simulation (ploidy) | N50 | Avg contigs per chromosome |
|---|---|---|
| Rice | 2 Mb | 15.6 |
| | 500 Kb | 62.5 |
| | 100 Kb | 310.8 |
| | 25 Kb | 1246.4 |
| *Arabidopsis* | 2 Mb | 12 |
| | 500 Kb | 48 |
| | 100 Kb | 239.2 |
| | 25 Kb | 952.4 |
| Human | 8 Mb | 16.7 |
| | 2 Mb | 66.7 |
| | 500 Kb | 265.9 |
| | 100 Kb | 1329.7 |

**Supplementary Table 12 |** Comparison between corrected and uncorrected genome assemblies of *S. spontaneum* Np-X for HapHiC and ALLHiC

| Contig length | Published assembly (before correction) | Assembly corrected by HapHiC | Assembly corrected by ALLHiC |
|---|---|---|---|
| Max length | 3,761,197 | 3,761,197 (-0%) | 1,779,478 (-52.67%) |
| N10 | 998,139 | 978,242 (-1.99%) | 314,000 (-68.54%) |
| N20 | 737,538 | 686,238 (-6.96%) | 241,636 (-67.24%) |
| N30 | 582,565 | 545,211 (-6.41%) | 195,608 (-66.42%) |
| N40 | 476,428 | 438,675 (-7.92%) | 161,904 (-66.02%) |
| N50 | 381,898 | 348,775 (-8.67%) | 138,999 (-63.60%) |
| N60 | 296,647 | 272,959 (-7.99%) | 118,448 (-60.07%) |
| N70 | 223,057 | 206,651 (-7.36%) | 98,999 (-55.62%) |
| N80 | 152,457 | 143,688 (-5.75%) | 80,999 (-46.87%) |
| N90 | 80,422 | 77,553 (-3.57%) | 55,709 (-30.73%) |

**Supplementary Table 13 |** Comparison of genome assemblies between *M.* × *giganteus* and other published *Miscanthus* species

| Species | Total length of assembly (Gb) | Number of contigs | Contig N50 (bp) | Is chromosome-level | Is haplotype-resolved |
|---|---|---|---|---|---|
| *M.* × *giganteus* | 6.11 | 5,823 | 2,176,999 | Yes | Yes |
| *M. sinensis* | 1.85 | 170,175 | 33,106 | Yes | No |
| *M. sacchariflorus* | 1.24 | 1,189,594 | 2,242 | No | No |
| *M. lutarioriparius* | 2.08 | 2,621 | 1,588,283 | Yes | No |
| *M. floridulus* | 2.68 | 7,351 | 673,655 | Yes | No |

**Supplementary Table 14 |** Pseudocode for optimal recommendation algorithm

| Optimal inflation recommendation |
| --- |
| $n$: the number of Markov clusters for a given inflation value<br>$k$: the known number of chromosomes<br>$r$: the length ratio threshold of adjacent clusters<br>$L$: a reversely sorted list to store Markov cluster length for a given inflation value<br>$D$: a dictionary with inflation values as keys and the number of clusters meeting the threshold $r$ as values<br>**for** float $i \in$ closed interval [min_inflation, max_inflation] **do**<br> **if** $n < k$ **then**<br> **continue**<br> **for** integer $j \in$ closed interval [1, $n - 1$] **do**<br> **if** $L[j]\ /\ L[j-1] < r$ **then**<br> **if** $j \geq k$ **then**<br> $D[i] = j$<br> **break**<br> $D[i] = n$<br>**return** min($D$, key=lambda $i$ : $D[i]$) |

#### Supplementary Figures

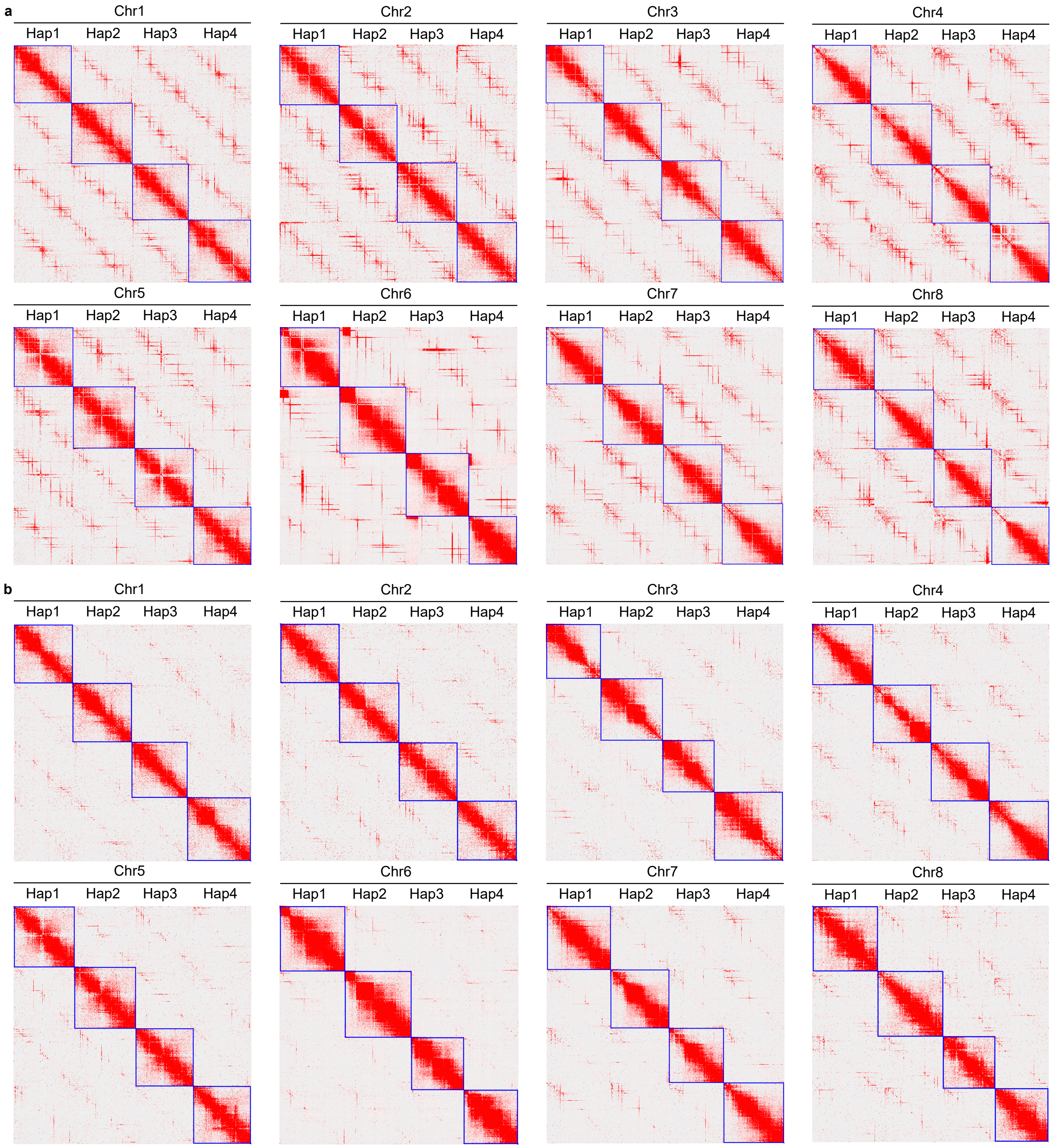


**Supplementary Fig. 1 |** Generation of the ground truth for evaluating scaffolding tools using the published haplotype-resolved autotetraploid genome of *M. sativa* XinJiangDaYe. **a**, The Hi-C contact maps of published *M. sativa* XinJiangDaYe genome. **b**, The Hi-C contact maps of the ground truth, generated after the manual removal of obvious assembly errors.

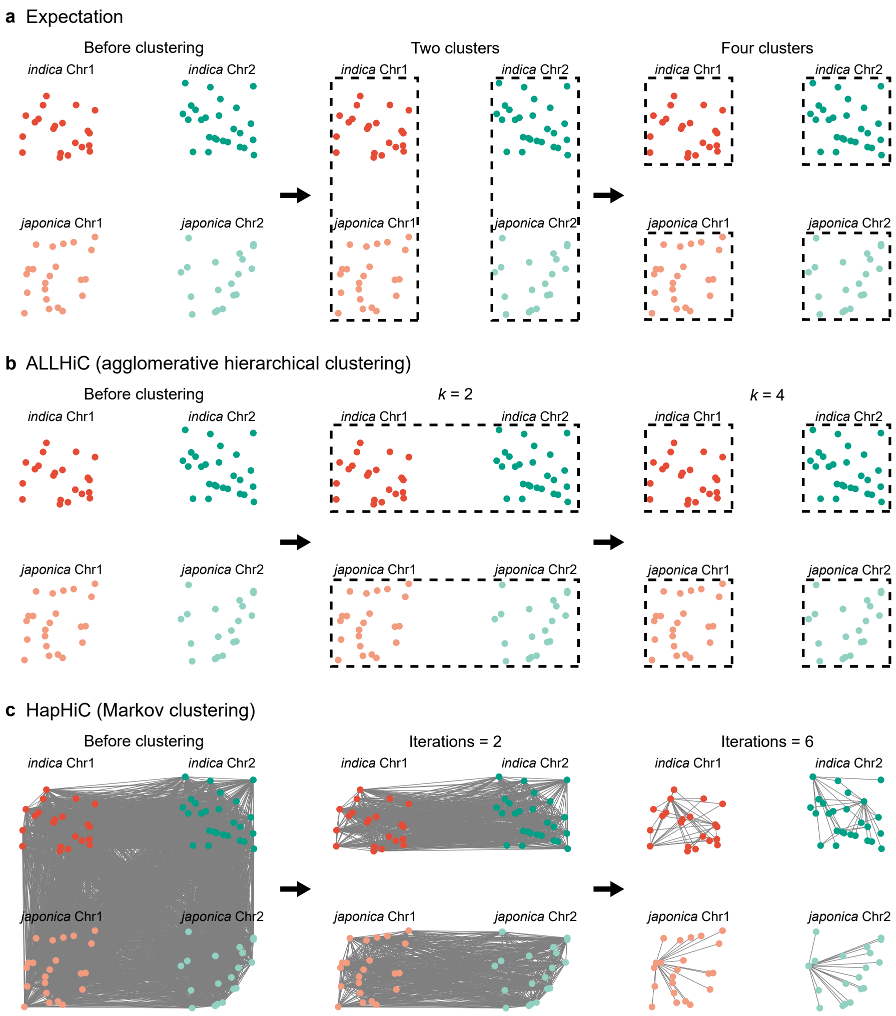


**Supplementary Fig. 2 |** Artificial merging of data from two rice subspecies, *indica* and *japonica*, leads to an underestimation of Hi-C links between homologous chromosomes. **a**, Ideally, the contigs between homologous chromosomes should be more difficult to separate than those between non-homologous chromosomes. **b**, However, the artificial merging of data from *indica* and *japonica* makes the contigs between homologous chromosomes easier to separate than those between non-homologous chromosomes in ALLHiC. **c**, This phenomenon can also be observed in HapHiC.

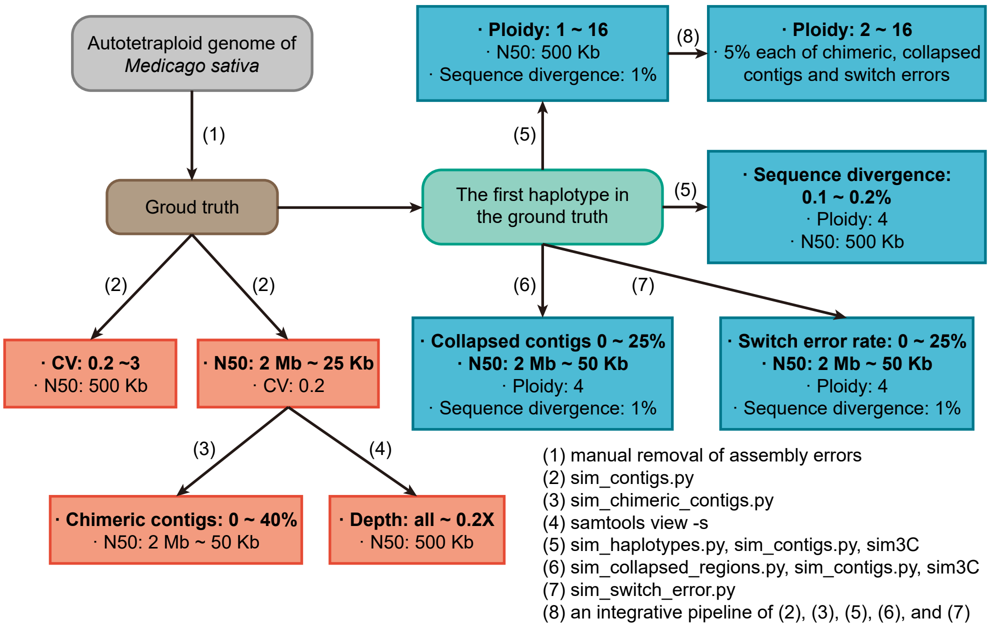


**Supplementary Fig. 3 |** Workflow for simulating datasets with various adverse factors. Rounded rectangles represent chromosome-level genomes in FASTA format, while rectangles denote simulated assemblies and/or Hi-C data. Arrows depict the simulation processes.

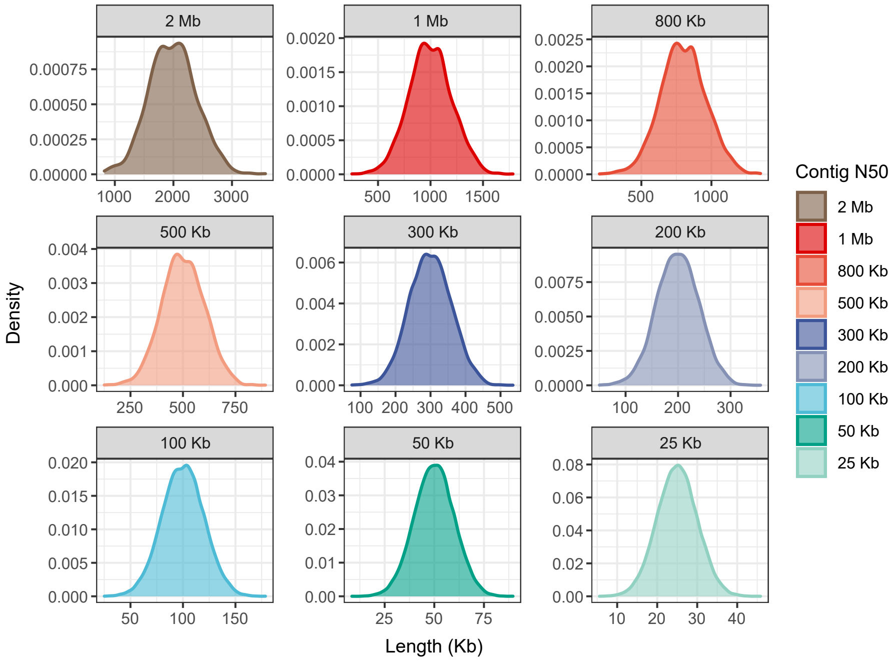


**Supplementary Fig. 4 |** Distribution of contig lengths in simulated assemblies, with the contig N50 values ranging from 2 Mb to 25 Kb.

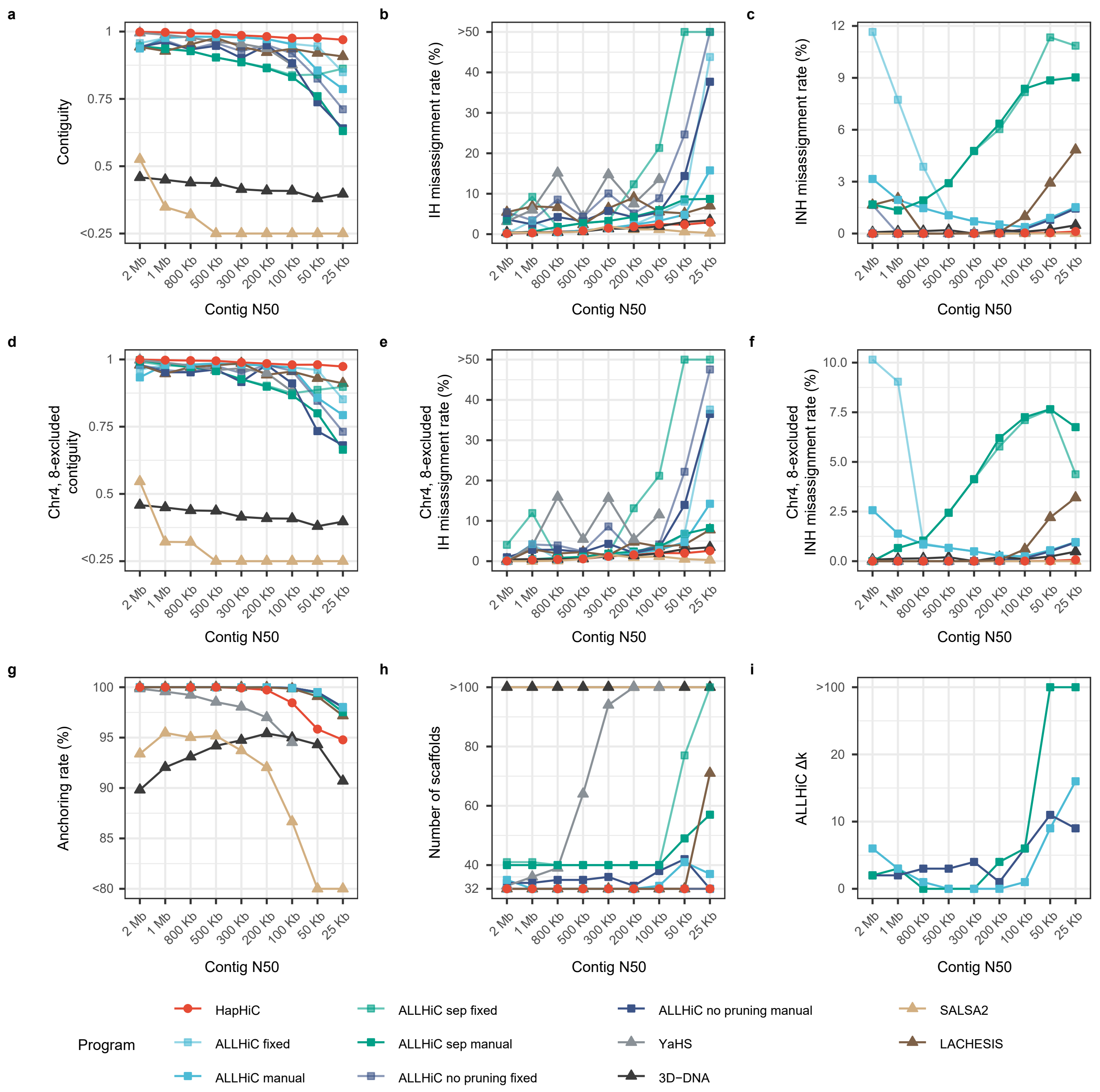


**Supplementary Fig. 5 |** Performance evaluation of Hi-C-based scaffolding tools in chromosome assignment on assemblies with varying contig N50 values. The evaluation metrics include contiguity (**a**), misassignment rate between homologous chromosomes (**b**), misassignment rate between non-homologous chromosomes (**c**), anchoring rate (**g**), number of scaffolds (**h**), and $\Delta k$ of manual parameter tuning for ALLHiC (**i**). During ALLHiC pruning, the genome of a closely related species, *M. truncatula* (MtrunA17r5.0-ANR), was used as a reference. However, chromosomes 4 and 8 of *M. truncatula* have some structural differences compared to those of *M. sativa*. Therefore, we also calculated the contiguity and misassignment rate after excluding these two chromosomes (**d-f**). The parameters for these scaffolding tools are provided in Section **3 Hi-C-based scaffolding** of **Software Commands**.

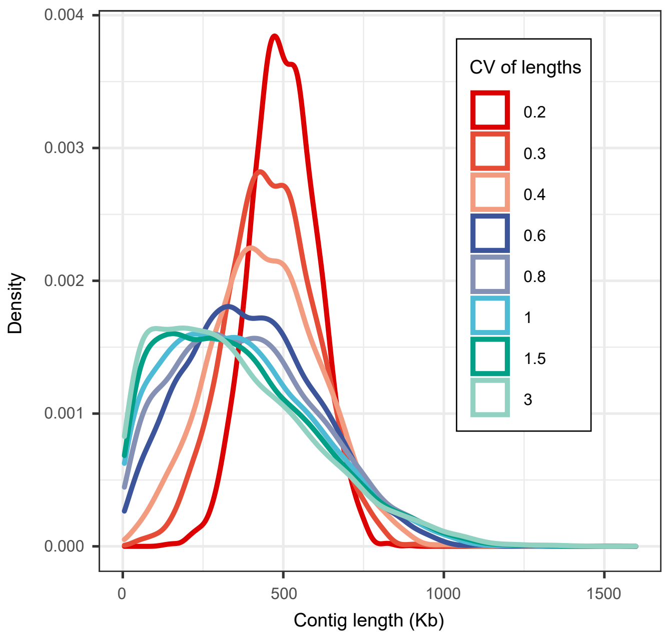


**Supplementary Fig. 6 |** Distribution of contig lengths in simulated assemblies, with the coefficients of variation (CVs) of contig length ranging from 0.2 to 3.

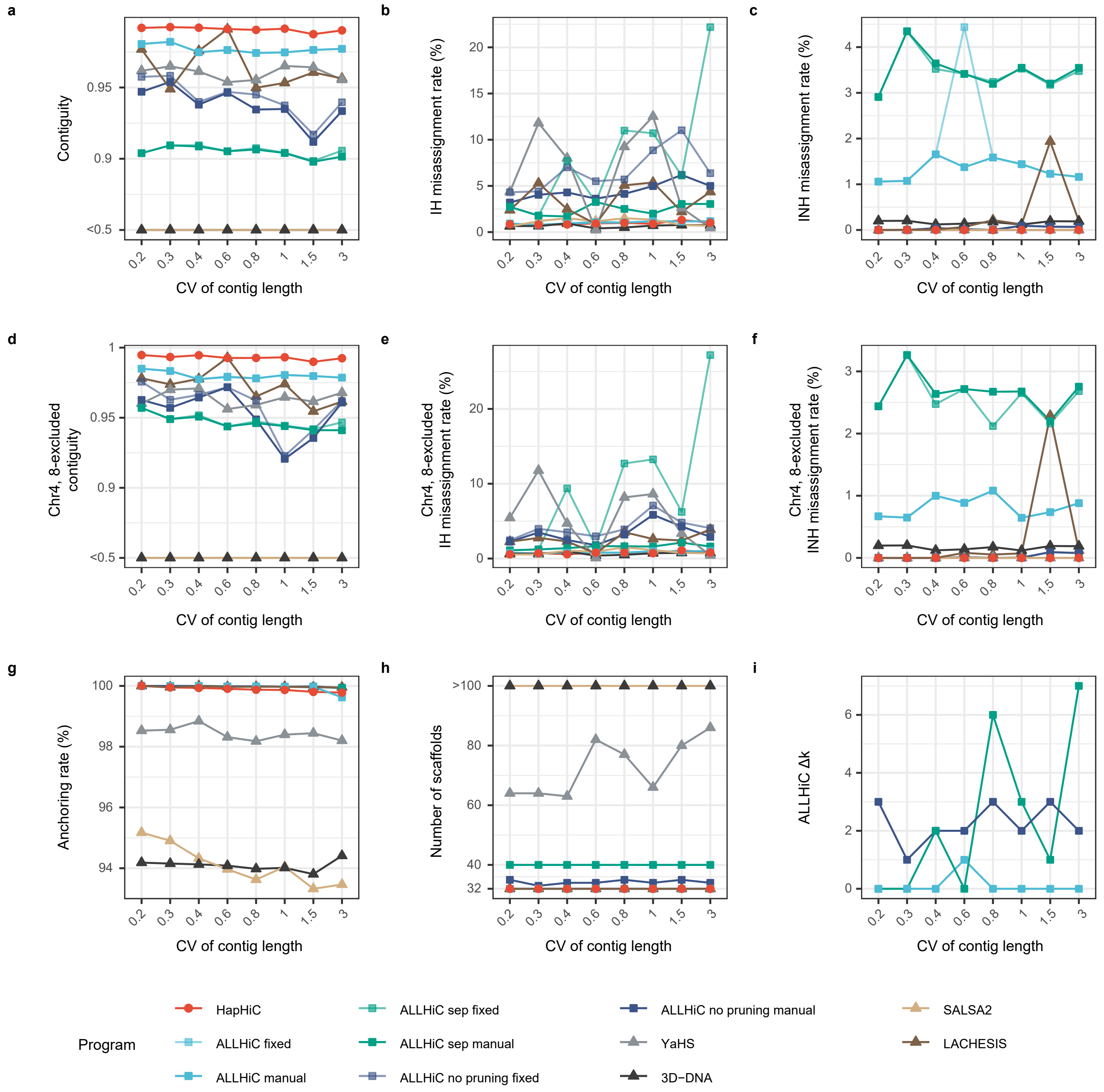


**Supplementary Fig. 7 |** Performance evaluation of Hi-C-based scaffolding tools in chromosome assignment on assemblies with varying coefficients of variation (CVs) of contig length. The evaluation metrics include contiguity (**a**), misassignment rate between homologous chromosomes (**b**), misassignment rate between non-homologous chromosomes (**c**), anchoring rate (**g**), number of scaffolds (**h**), and $\Delta k$ of manual parameter tuning for ALLHiC (**i**). During ALLHiC pruning, the genome of a closely related species, *M. truncatula* (MtrunA17r5.0-ANR), was used as a reference. However, chromosomes 4 and 8 of *M. truncatula* have some structural differences compared to those of *M. sativa*. Therefore, we also calculated the contiguity and misassignment rate after excluding these two chromosomes (**d-f**). The parameters for these scaffolding tools are provided in Section **3 Hi-C-based scaffolding** of **Software Commands**.

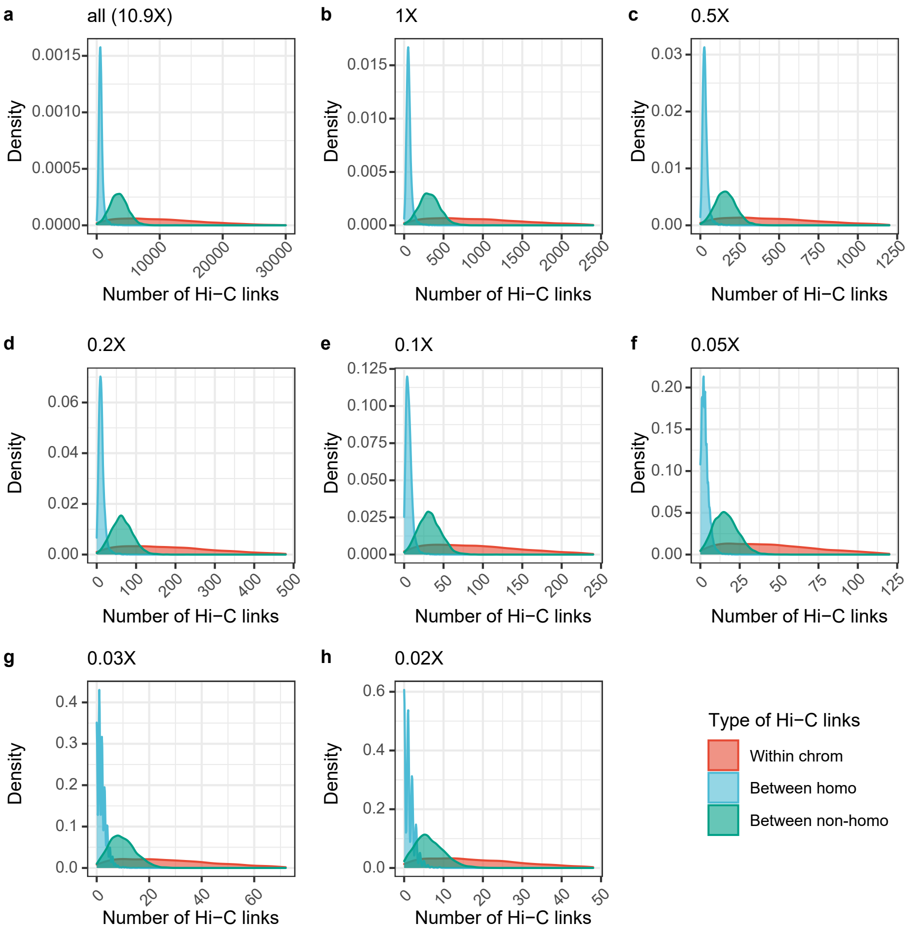


**Supplementary Fig. 8 |** Distribution of the number of Hi-C links for contigs at varying effective sequencing depths ranging from 10.9X to 0.02X (**a-h**).

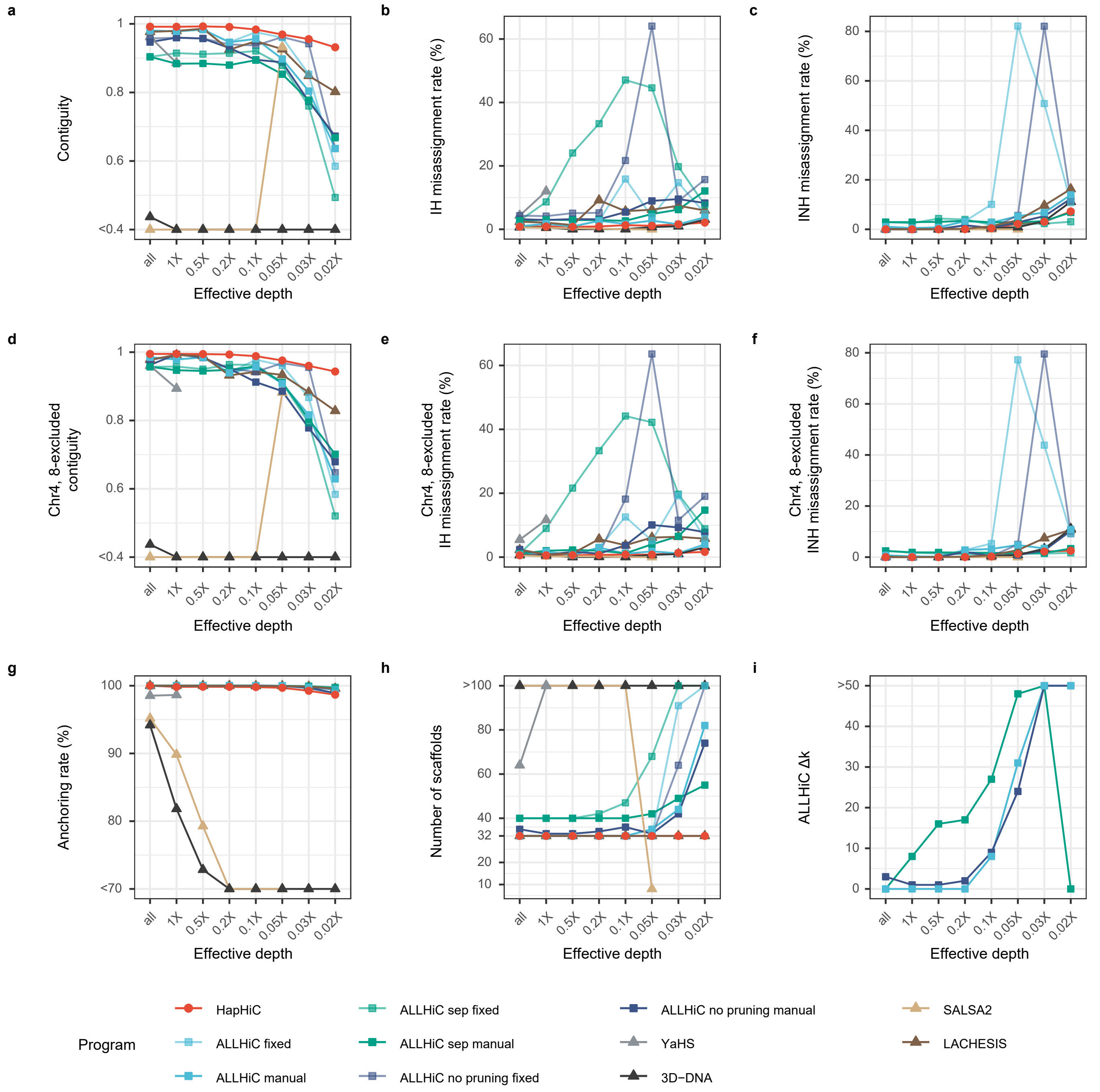


**Supplementary Fig. 9 |** Performance evaluation of Hi-C-based scaffolding tools in chromosome assignment on assemblies with varying effective Hi-C sequencing depths. The evaluation metrics include contiguity (**a**), misassignment rate between homologous chromosomes (**b**), misassignment rate between non-homologous chromosomes (**c**), anchoring rate (**g**), number of scaffolds (**h**), and $\Delta k$ of manual parameter tuning for ALLHiC (**i**). During ALLHiC pruning, the genome of a closely related species, *M. truncatula* (MtrunA17r5.0-ANR), was used as a reference. However, chromosomes 4 and 8 of *M. truncatula* have some structural differences compared to those of *M. sativa*. Therefore, we also calculated the contiguity and misassignment rate after excluding these two chromosomes (**d-f**). The parameters for these scaffolding tools are provided in Section **3 Hi-C-based scaffolding** of **Software Commands**.

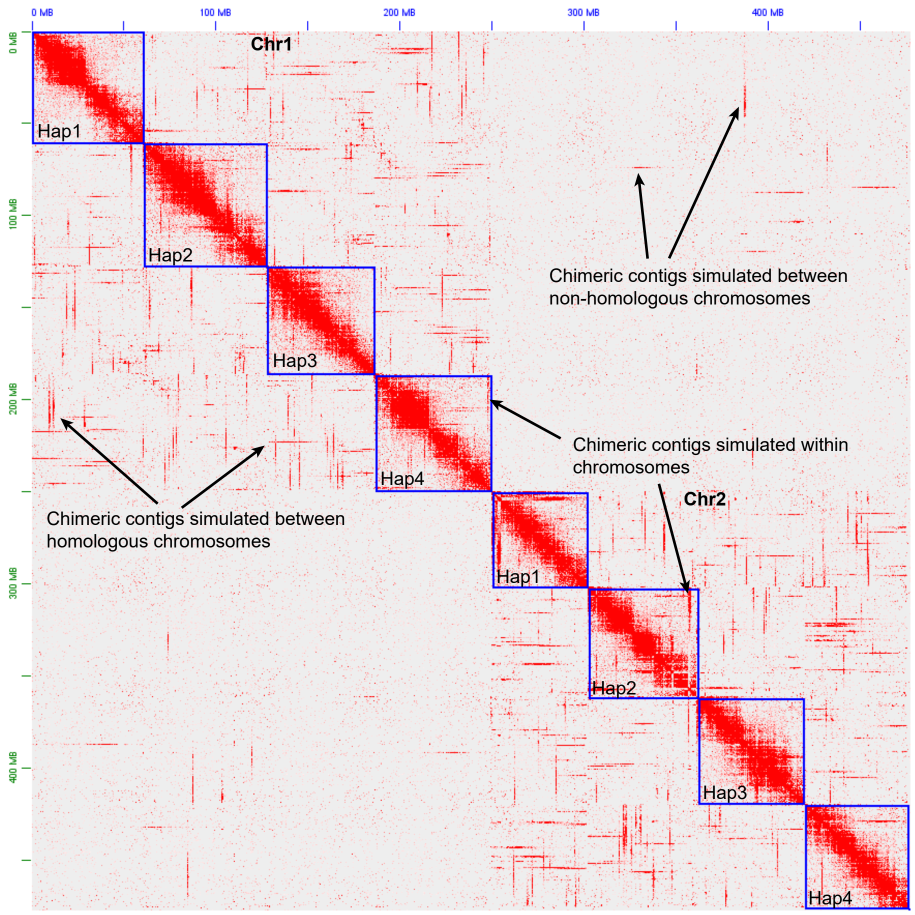


**Supplementary Fig. 10 |** Hi-C contact map of simulated chimeric contigs. Arrows indicate examples of simulated chimeric contigs.

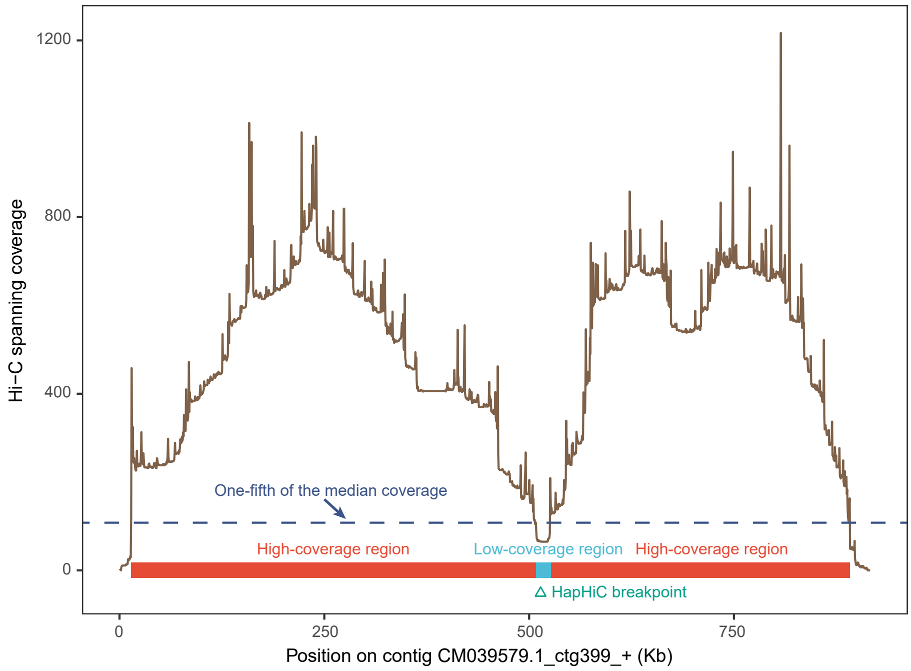


**Supplementary Fig. 11 |** Assembly correction method in HapHiC. The brown line chart represents the Hi-C spanning coverage along contig CM039579.1_ctg399_+. The dashed blue line illustrates the coverage threshold, which is set at one-fifth of the median coverage by default. Regions of high and low coverage are denoted by red and blue rectangles, respectively. The green triangle indicates the breakpoint as determined by HapHiC.

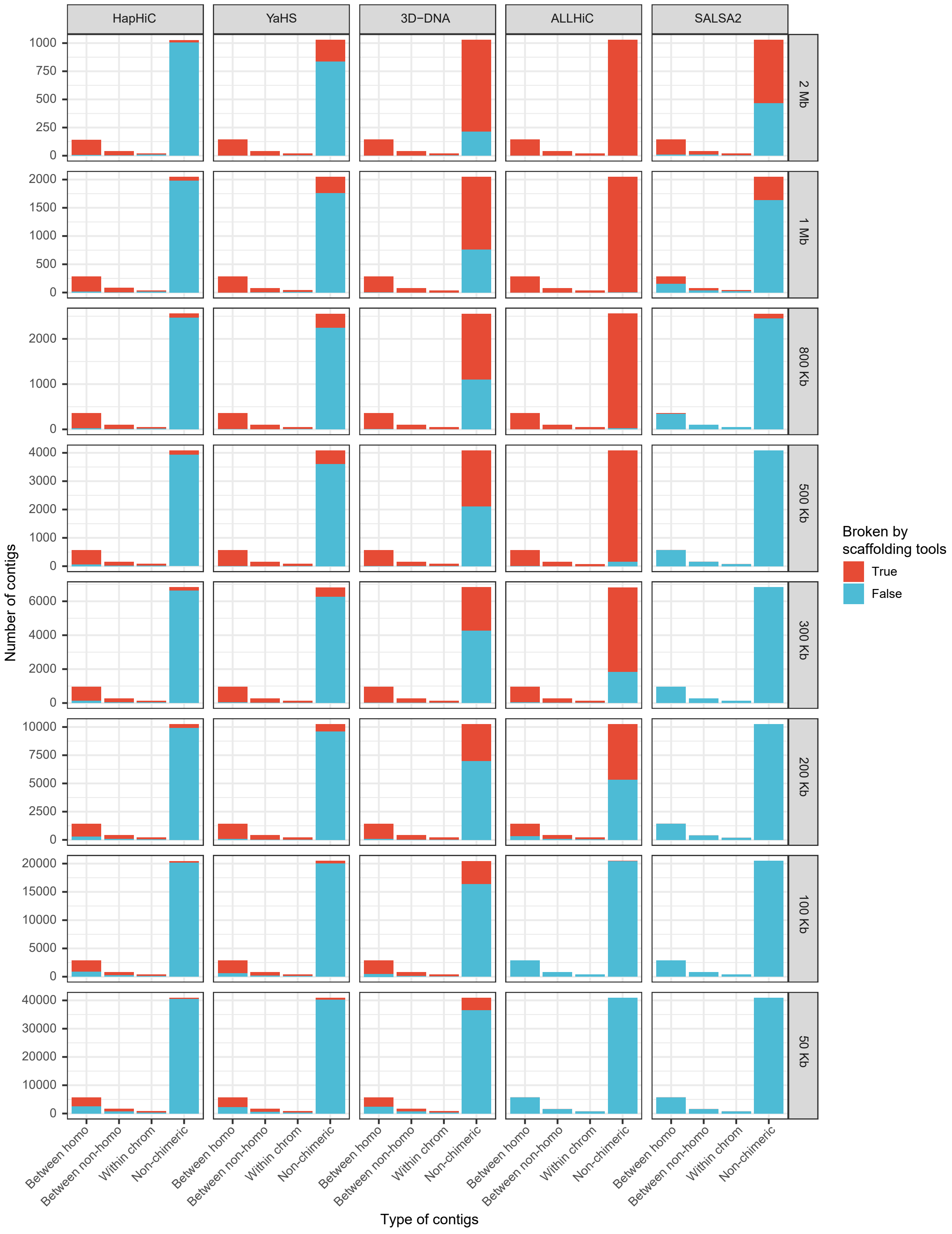


**Supplementary Fig. 12 |** Performance evaluation of Hi-C-based scaffolding tools in identifying chimeric contigs. The chimeric contigs are classified based on their simulated misjoins: between homologous chromosomes (between homo), between non-homologous chromosomes (between non-homo), and within the same chromosome (within chrom). Ideally, all simulated chimeric contigs should be identified and broken by Hi-C-based scaffolding tools (indicated in red), while non-chimeric contigs should remain intact (indicated in blue).

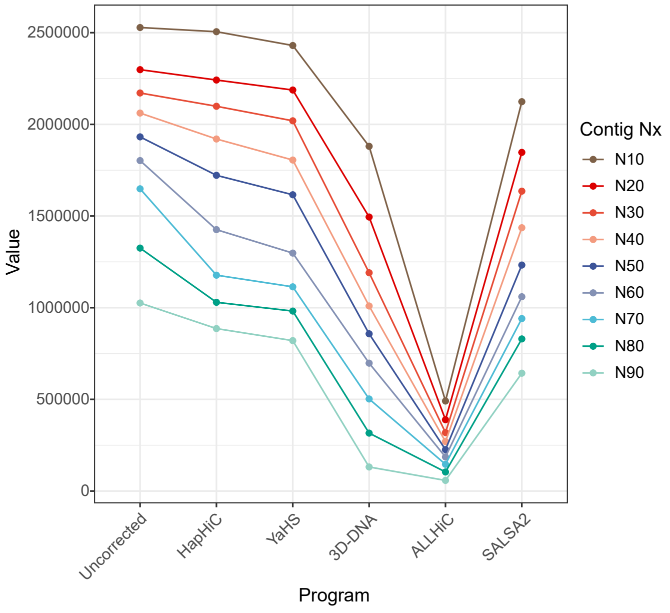


**Supplementary Fig. 13 |** Comparison of Contig N10 to N90 values of assemblies before and after assembly correction by each Hi-C-based scaffolding tool.

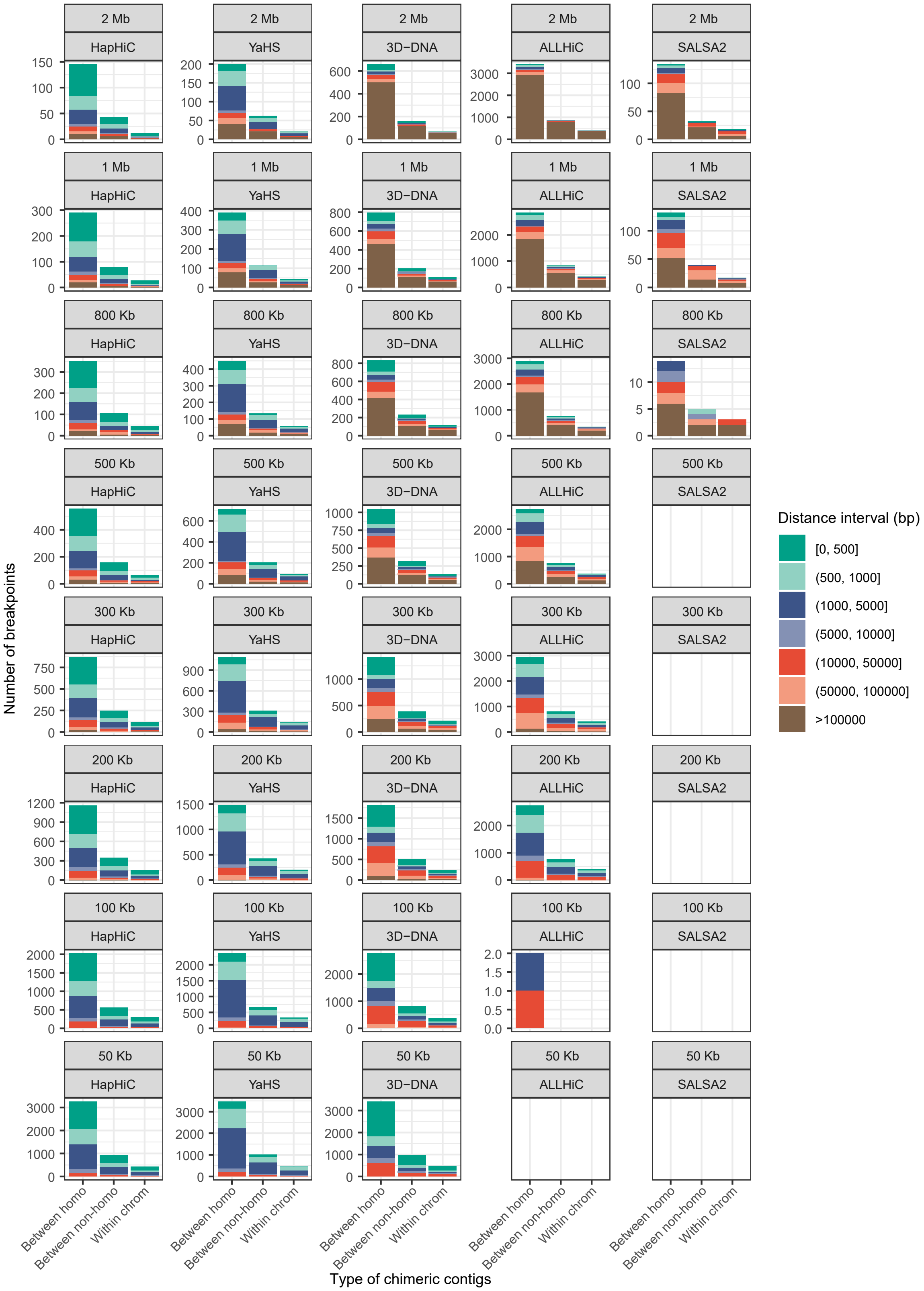


**Supplementary Fig. 14 |** Performance evaluation of Hi-C-based scaffolding tools in determining breakpoints for chimeric contigs. The chimeric contigs are classified based on their simulated misjoins: between homologous chromosomes (between homo), between non-homologous chromosomes (between non-homo), and within the same chromosome (within chrom). A closer distance between the determined breakpoint and the actual simulated misjoin indicates better results.

**a**

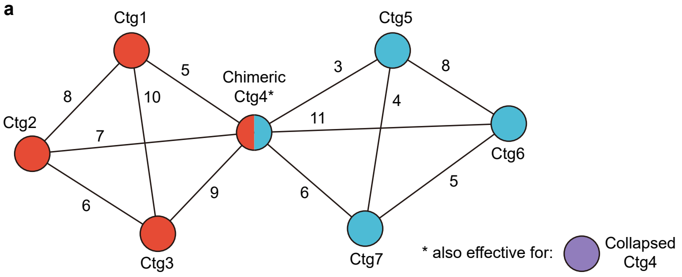


**b** Ranking of each contig relative to others

| From \ To | Ctg1 | Ctg2 | Ctg3 | Ctg4 | Ctg5 | Ctg6 | Ctg7 |
|---|---|---|---|---|---|---|---|
| Ctg1 | - | **2** | **1** | **3** | 4 | 5 | 6 |
| Ctg2 | **1** | - | **3** | **2** | 4 | 5 | 6 |
| Ctg3 | **1** | **3** | - | **2** | 4 | 5 | 6 |
| Ctg4 | 5 | 3 | **2** | - | 6 | **1** | 4 |
| Ctg5 | 4 | 5 | 6 | **3** | - | **1** | **2** |
| Ctg6 | 4 | 5 | 6 | **1** | **2** | - | **3** |
| Ctg7 | 4 | 5 | 6 | **1** | **3** | **2** | - |

**c** Calculation of rank-sum values ($n$ = 3)

| Contig | #1 | #2 | #3 | min(#1→#2, #2→#1) + min(#2→#3, #3→#2) + min(#1→#3, #3→#1) |
|---|---|---|---|---|
| Ctg1 | Ctg3 | Ctg2 | Ctg4 | min(3, 3) + min(2, 3) + min(2, 2) = 7 |
| Ctg2 | Ctg1 | Ctg4 | Ctg3 | min(3, 5) + min(2, 2) + min(1, 1) = 6 |
| Ctg3 | Ctg1 | Ctg4 | Ctg2 | min(3, 5) + min(3, 2) + min(2, 1) = 6 |
| Ctg4 | Ctg6 | Ctg3 | Ctg2 | min(6, 5) + min(3, 3) + min(5, 5) = 13 |
| Ctg5 | Ctg6 | Ctg7 | Ctg4 | min(3, 2) + min(1, 4) + min(1, 1) = 4 |
| Ctg6 | Ctg4 | Ctg5 | Ctg7 | min(6, 3) + min(2, 3) + min(4, 1) = 7 |
| Ctg7 | Ctg4 | Ctg6 | Ctg5 | min(1, 1) + min(2, 1) + min(6, 3) = 5 |

**Supplementary Fig. 15 |** Rank-sum algorithm for identifying chimeric and collapsed contigs in HapHiC. **a**, A network graph illustrates contigs connected via Hi-C links. Red and blue circles (vertices) represent contigs from different haplotypes. Bicolor and purple circles symbolize chimeric and collapsed contigs, respectively. Edges connecting these circles indicate Hi-C links between contigs, with the number of Hi-C links displayed adjacent to the edges. **b**, The ranking of each contig relative to others based on the number of Hi-C links. Gray numbers indicate the absence of direct connections between contigs. **c**, The calculation of rank-sum values. Different colors are used to trace the origin of the ranks.

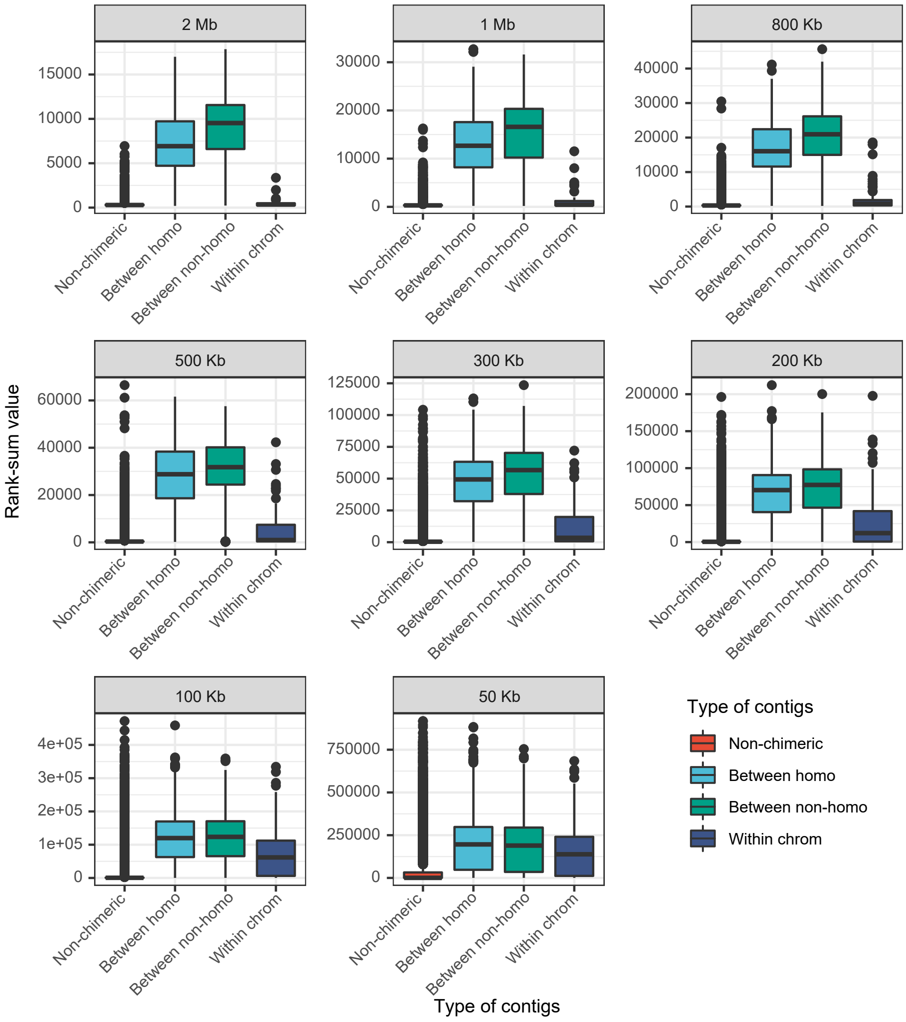


**Supplementary Fig. 16 |** Rank-sum values of chimeric and non-chimeric contigs of varying lengths, ranging from 2 Mb to 50 Kb. The chimeric contigs are classified based on their simulated misjoins: between homologous chromosomes (between homo), between non-homologous chromosomes (between non-homo), and within the same chromosome (within chrom).

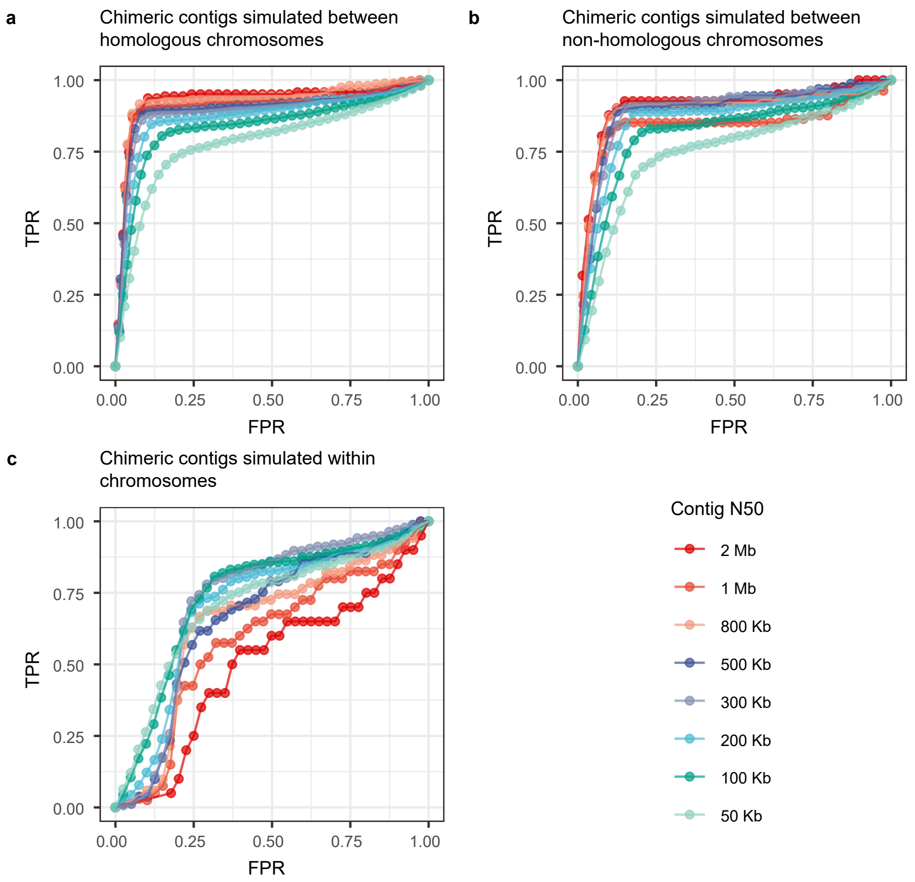


**Supplementary Fig. 17 |** Receiver operating characteristics (ROC) curves demonstrate the performance of the rank-sum algorithm in identifying chimeric contigs of varying lengths. The chimeric contigs are classified based on their simulated misjoins: between homologous chromosomes (**a**), between non-homologous chromosomes (**b**), and within the same chromosome (**c**).

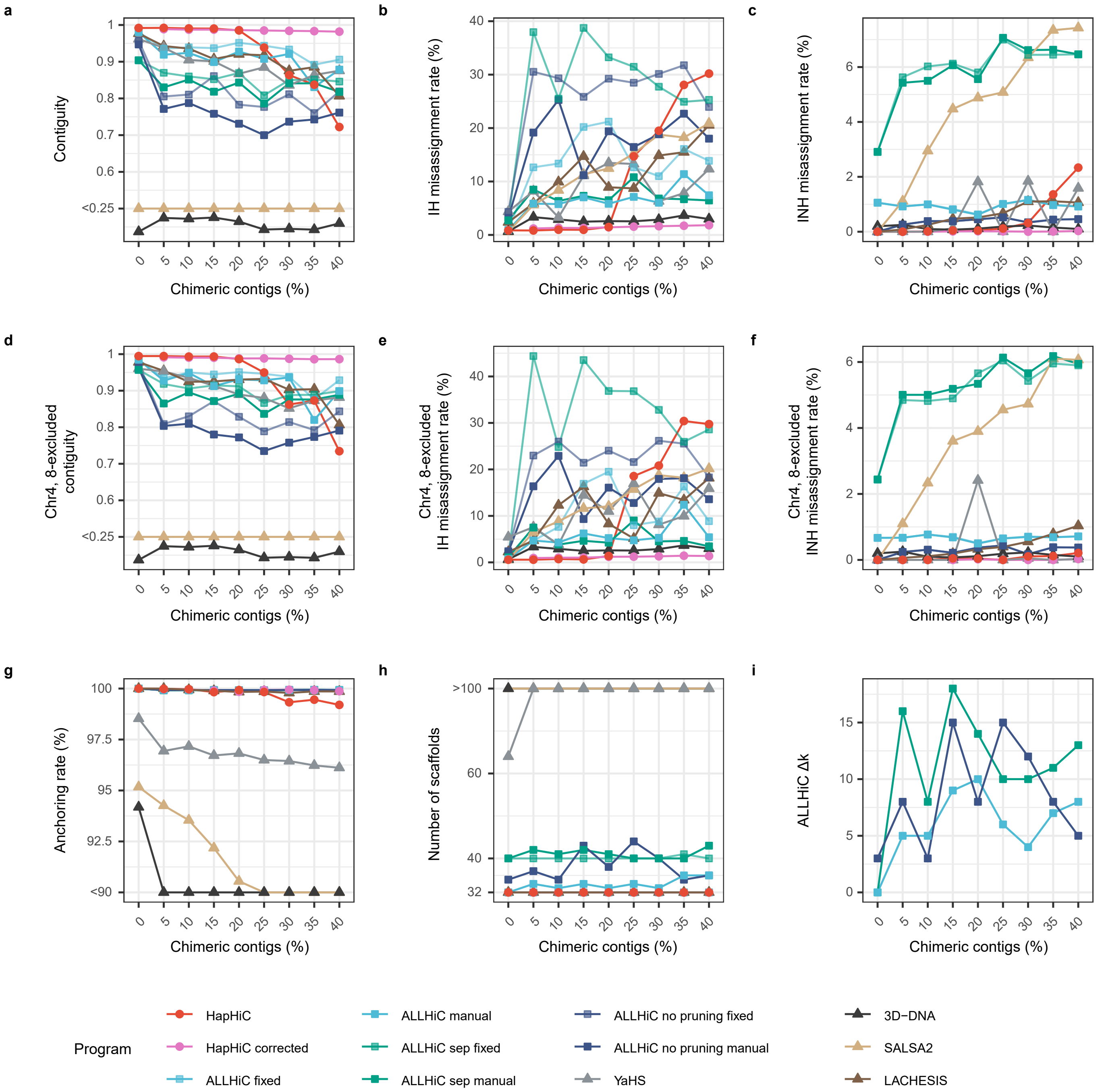


**Supplementary Fig. 18 |** Performance evaluation of Hi-C-based scaffolding tools in chromosome assignment on assemblies with varying proportions of chimeric contigs. The evaluation metrics include contiguity (**a**), misassignment rate between homologous chromosomes (**b**), misassignment rate between non-homologous chromosomes (**c**), anchoring rate (**g**), number of scaffolds (**h**), and Δ*k* of manual parameter tuning for ALLHiC (**i**). During ALLHiC pruning, the genome of a closely related species, *M. truncatula* (MtrunA17r5.0-ANR), was used as a reference. However, chromosomes 4 and 8 of *M. truncatula* have some structural differences compared to those of *M. sativa*. Therefore, we also calculated the contiguity and misassignment rate after excluding these two chromosomes (**d-f**). The parameters for these scaffolding tools are provided in Section **3 Hi-C-based scaffolding** of **Software Commands**.

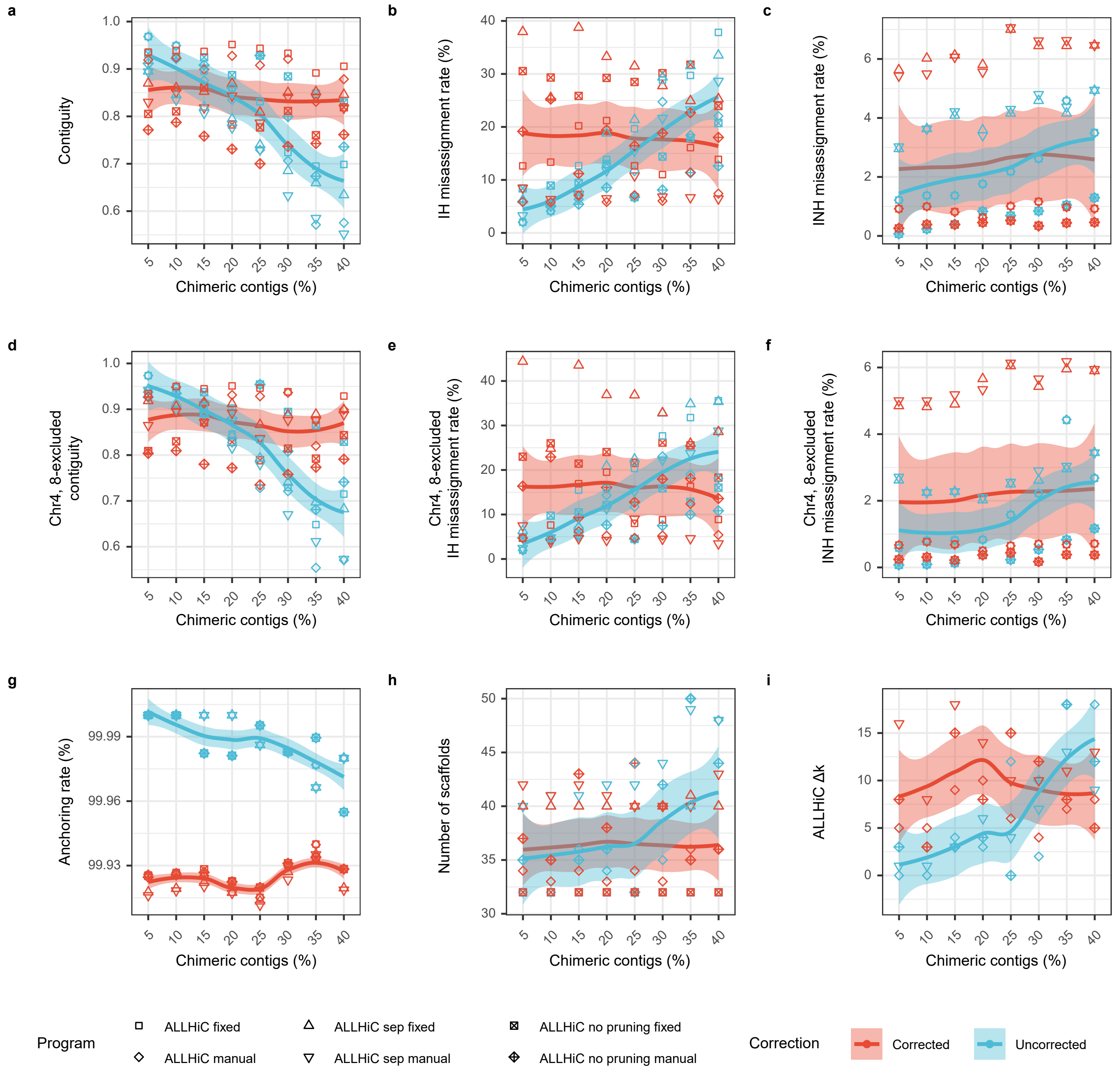


**Supplementary Fig. 19 |** Performance evaluation of ALLHiC with and without assembly correction in chromosome assignment on assemblies with varying proportions of chimeric contigs. The evaluation metrics include contiguity (**a**), misassignment rate between homologous chromosomes (**b**), misassignment rate between non-homologous chromosomes (**c**), anchoring rate (**g**), number of scaffolds (**h**), and $\Delta k$ of manual parameter tuning for ALLHiC (**i**). During ALLHiC pruning, the genome of a closely related species, *M. truncatula* (MtrunA17r5.0-ANR), was used as a reference. However, chromosomes 4 and 8 of *M. truncatula* have some structural differences compared to those of *M. sativa*. Therefore, we also calculated the contiguity and misassignment rate after excluding these two chromosomes (**d**-**f**). The parameters for these scaffolding tools are provided in Section **3 Hi-C-based scaffolding** of **Software Commands**.

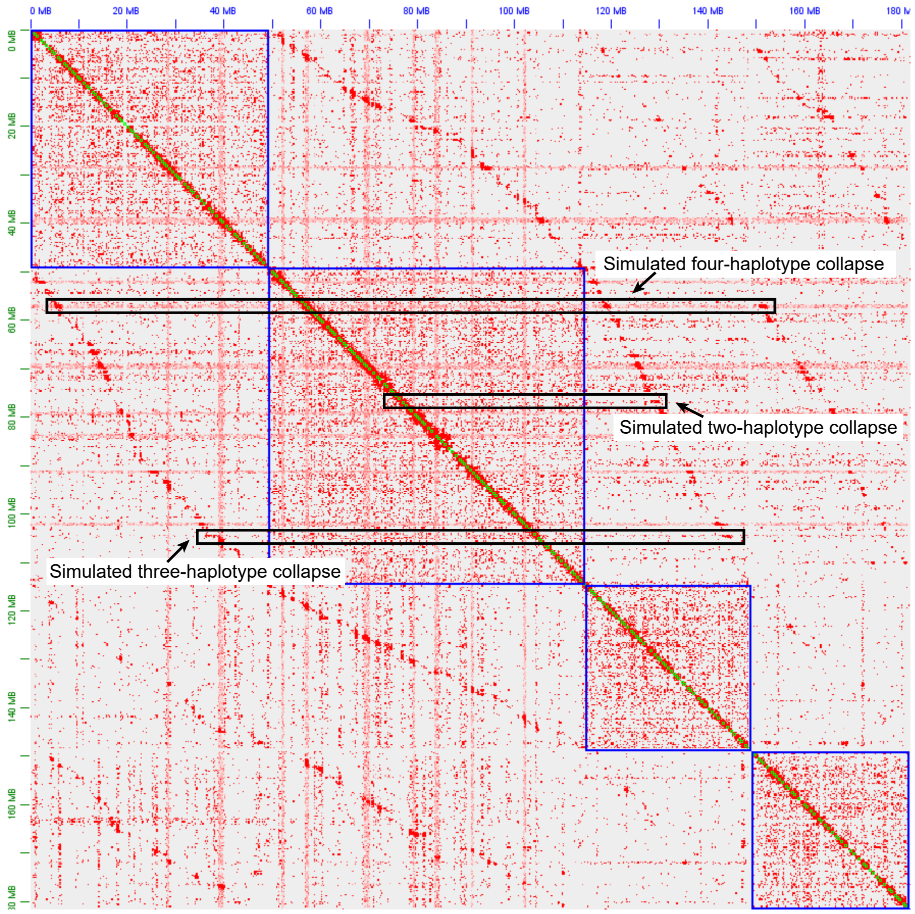


**Supplementary Fig. 20 |** Hi-C contact map of simulated collapsed contigs. Arrows indicate examples of collapsed chimeric contigs.

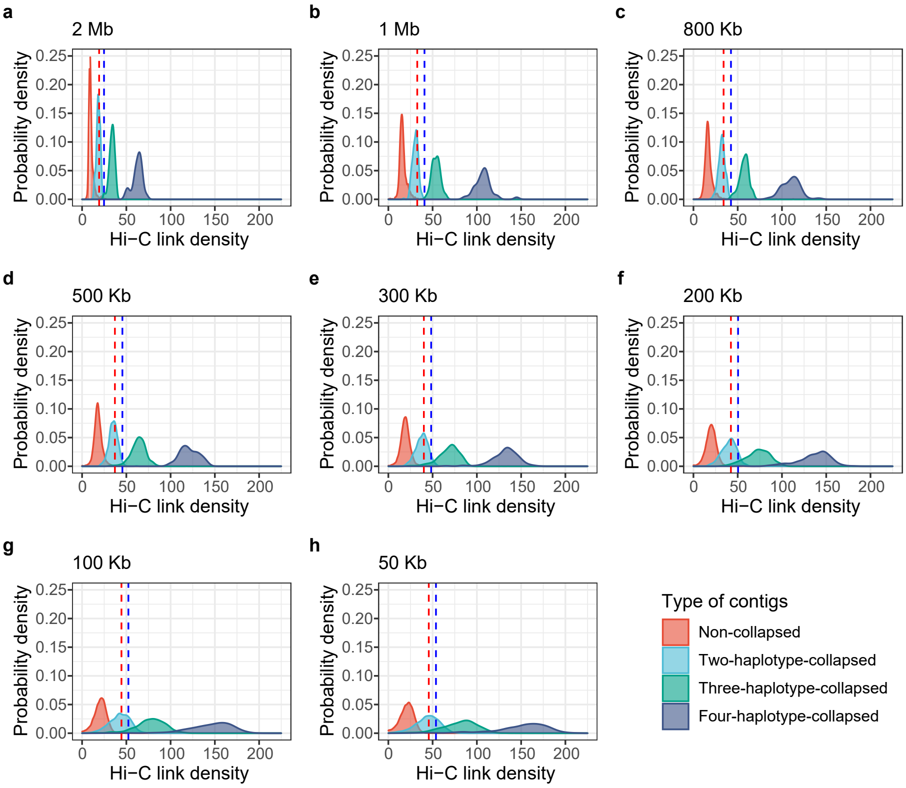


**Supplementary Fig. 21 |** Distribution of Hi-C link density for collapsed and non-collapsed contigs, with the contig lengths ranging from 2 Mb to 50 Kb (**a**-**h**). The collapsed contigs are classified based on the number of haplotypes contributing to the collapse: two haplotypes (two-haplotype-collapsed), three haplotypes (three-haplotype-collapsed), and four haplotypes (four-haplotype-collapsed).

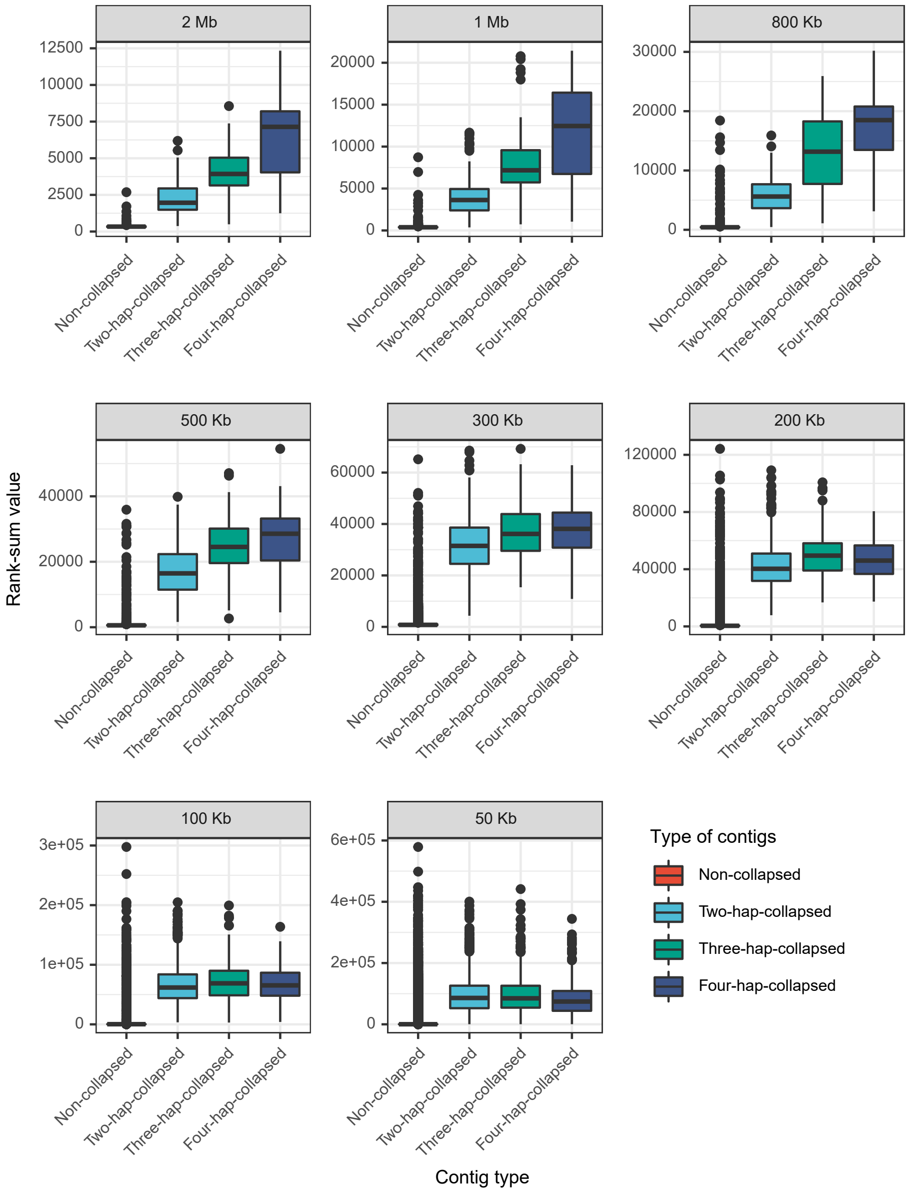


**Supplementary Fig. 22 |** Rank-sum values of collapsed and non-collapsed contigs of varying lengths, ranging from 2 Mb to 50 Kb. The collapsed contigs are classified based on the number of haplotypes contributing to the collapse: two haplotypes (two-haplotype-collapsed), three haplotypes (three-haplotype-collapsed), and four haplotypes (four-haplotype-collapsed).

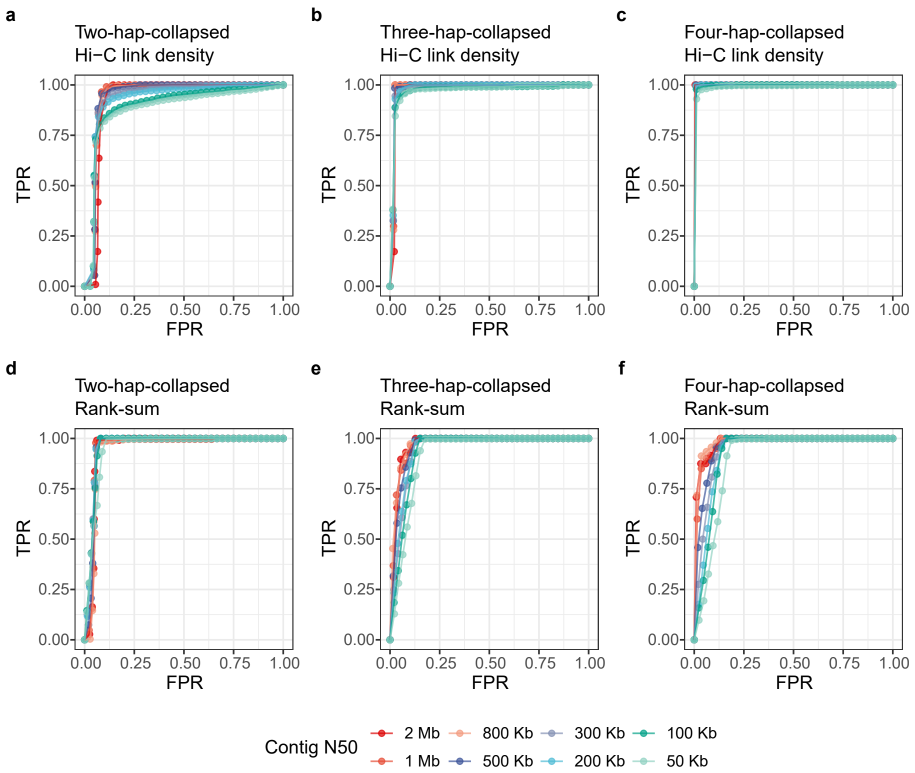


**Supplementary Fig. 23 |** Receiver operating characteristics (ROC) curves demonstrate the performance of link density and rank-sum algorithm in identifying chimeric contigs of varying contig lengths. The collapsed contigs are classified based on the number of haplotypes contributing to the collapse: two haplotypes (two-haplotype-collapsed, **a** and **d**), three haplotypes (three-haplotype-collapsed, **b** and **e**), and four haplotypes (four-haplotype-collapsed, **c** and **f**).

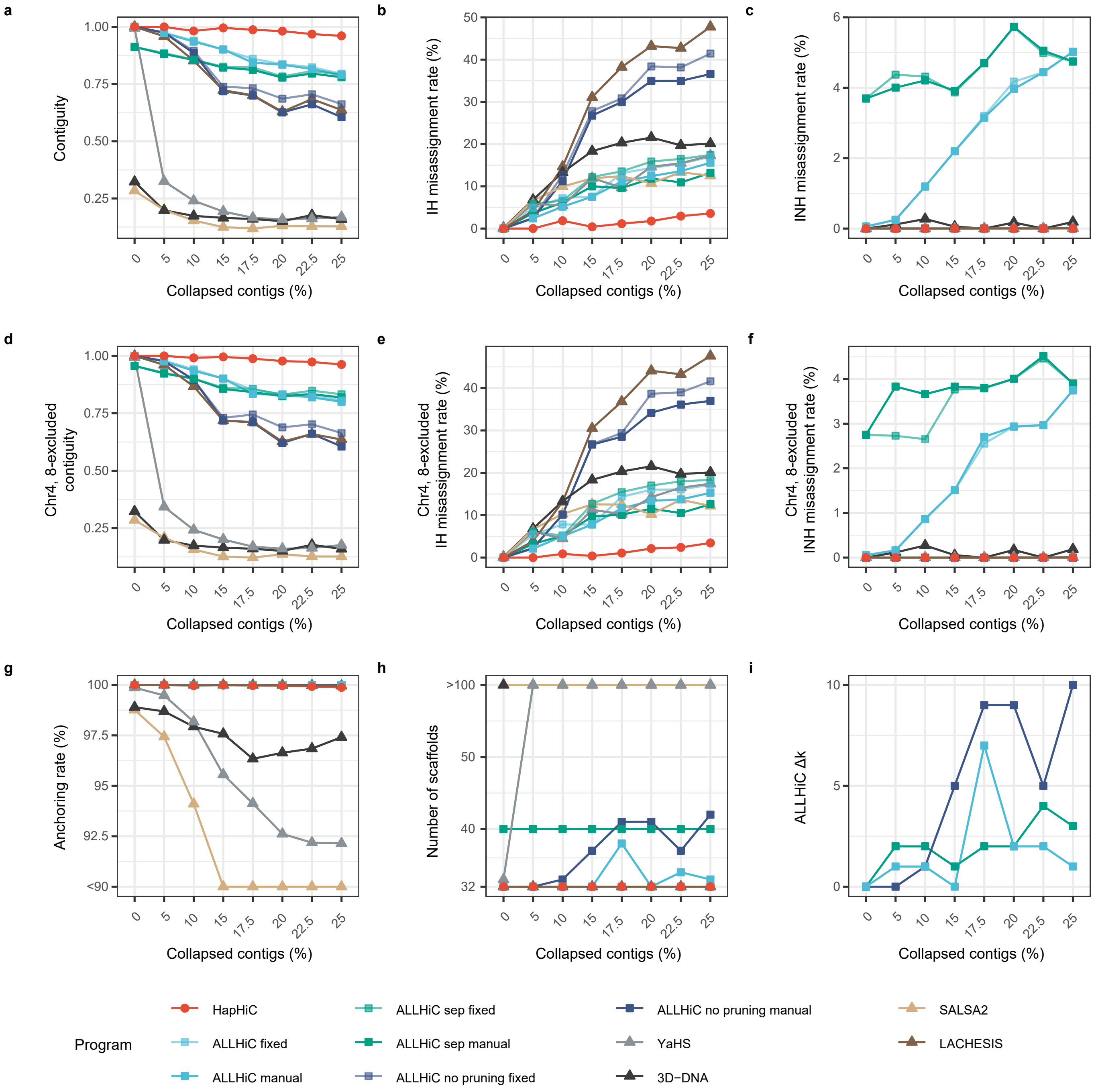


**Supplementary Fig. 24 |** Performance evaluation of Hi-C-based scaffolding tools in chromosome assignment on assemblies with varying proportions of collapsed contigs. The evaluation metrics include contiguity (**a**), misassignment rate between homologous chromosomes (**b**), misassignment rate between non-homologous chromosomes (**c**), anchoring rate (**g**), number of scaffolds (**h**), and $\Delta k$ of manual parameter tuning for ALLHiC (**i**). During ALLHiC pruning, the genome of a closely related species, *M. truncatula* (MtrunA17r5.0-ANR), was used as a reference. However, chromosomes 4 and 8 of *M. truncatula* have some structural differences compared to those of *M. sativa*. Therefore, we also calculated the contiguity and misassignment rate after excluding these two chromosomes (**d-f**). The parameters for these scaffolding tools are provided in Section **3 Hi-C-based scaffolding** of **Software Commands**.

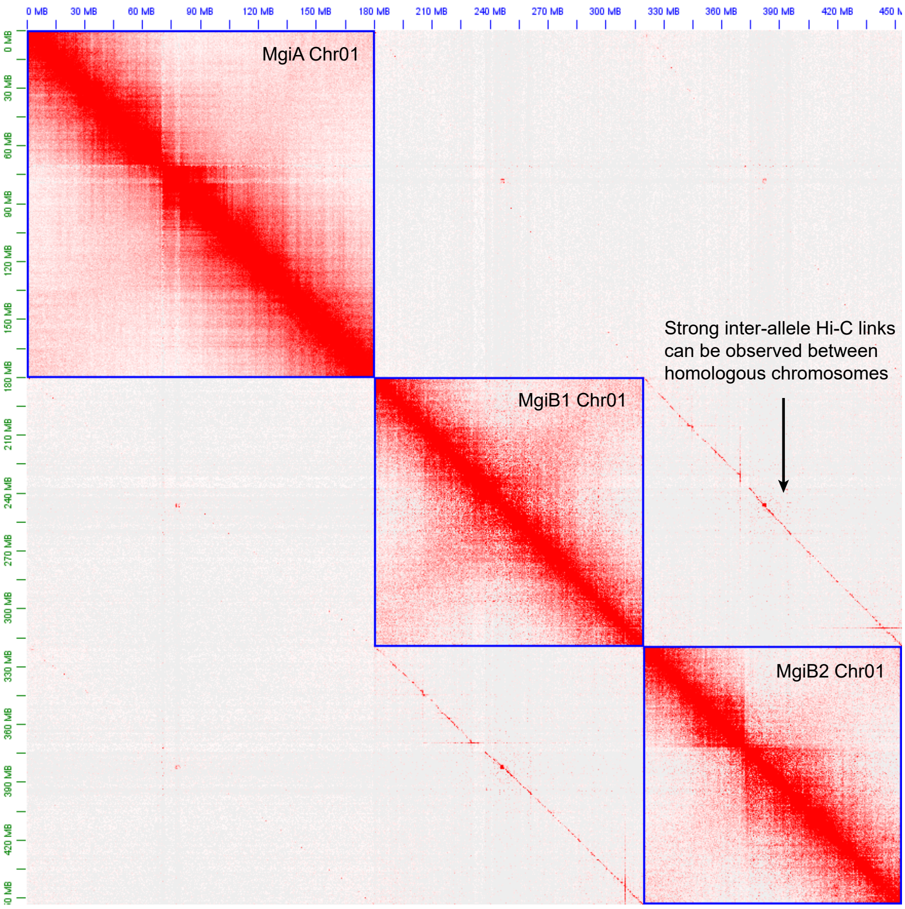


**Supplementary Fig. 25 |** Hi-C contact map for the three haplotypes of chromosome 1 in the *M.* × *giganteus* genome illustrates the strong inter-allele Hi-C links between homologous chromosomes.

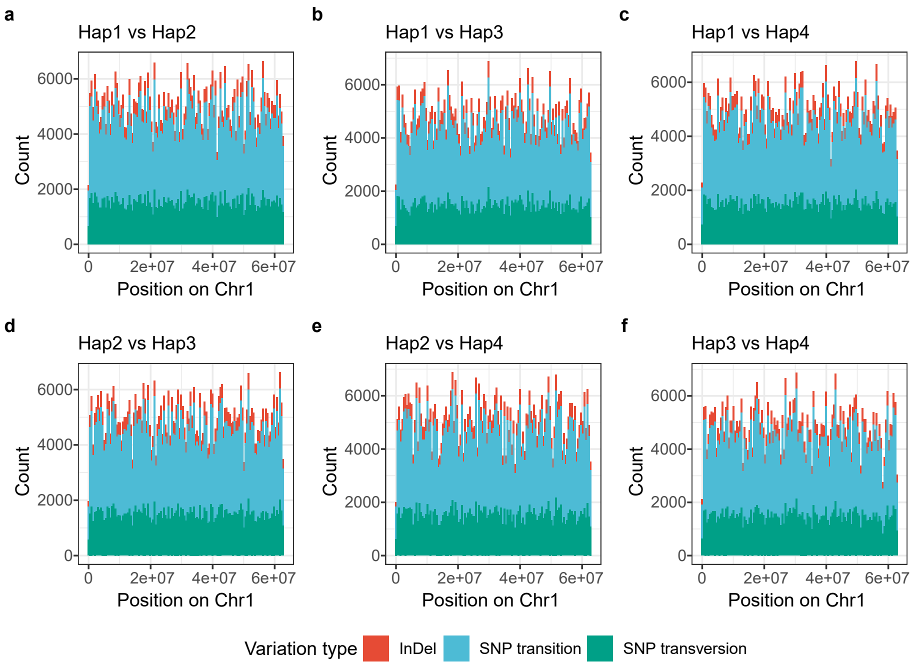


**Supplementary Fig. 26 |** Distribution of simulated variations between haplotypes along chromosome 1. The comparisons include: haplotype 1 vs haplotype 2 (**a**), haplotype 1 vs haplotype 3 (**b**), haplotype 1 vs haplotype 4 (**c**), haplotype 2 vs haplotype 3 (**d**), haplotype 2 vs haplotype 4 (**e**), haplotype 3 vs haplotype 4 (**f**).

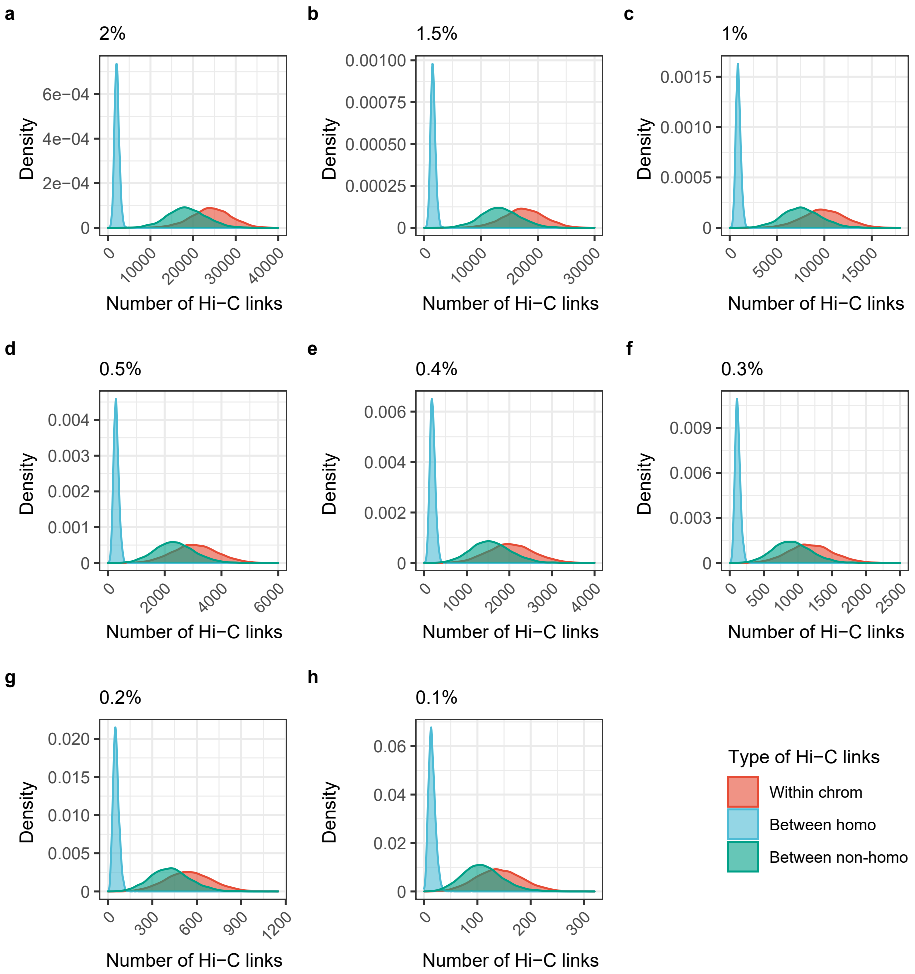


**Supplementary Fig. 27 |** Distribution of the number of Hi-C links for contigs, with the sequence divergence levels ranging from 2% to 0.1% (**a-h**).

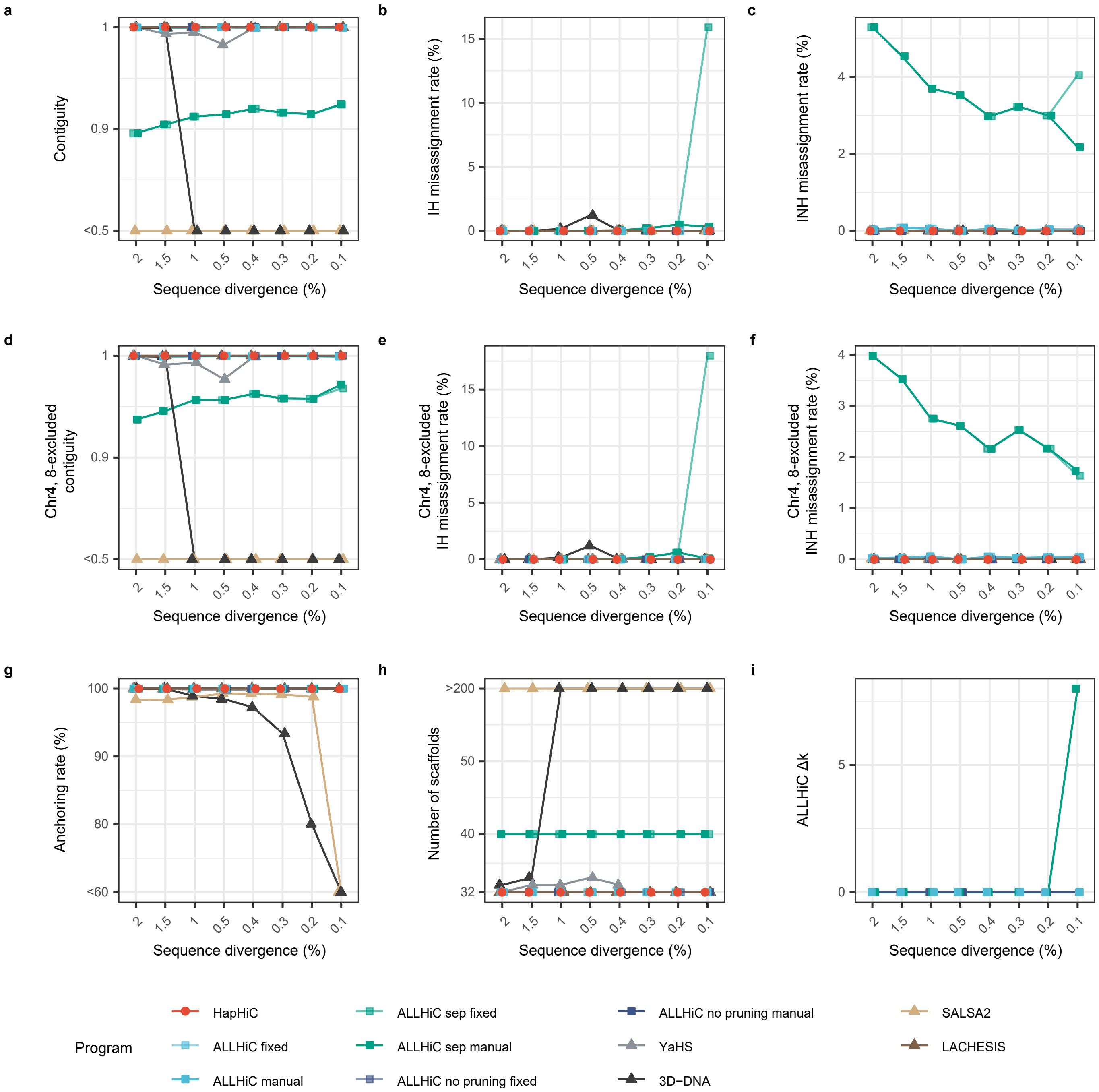


**Supplementary Fig. 28 |** Performance evaluation of Hi-C-based scaffolding tools in chromosome assignment on assemblies with varying sequence divergence levels. The evaluation metrics include contiguity (**a**), misassignment rate between homologous chromosomes (**b**), misassignment rate between non-homologous chromosomes (**c**), anchoring rate (**g**), number of scaffolds (**h**), and $\Delta k$ of manual parameter tuning for ALLHiC (**i**). During ALLHiC pruning, the genome of a closely related species, *M. truncatula* (MtrunA17r5.0-ANR), was used as a reference. However, chromosomes 4 and 8 of *M. truncatula* have some structural differences compared to those of *M. sativa*. Therefore, we also calculated the contiguity and misassignment rate after excluding these two chromosomes (**d-f**). The parameters for these scaffolding tools are provided in Section **3 Hi-C-based scaffolding** of **Software Commands**.

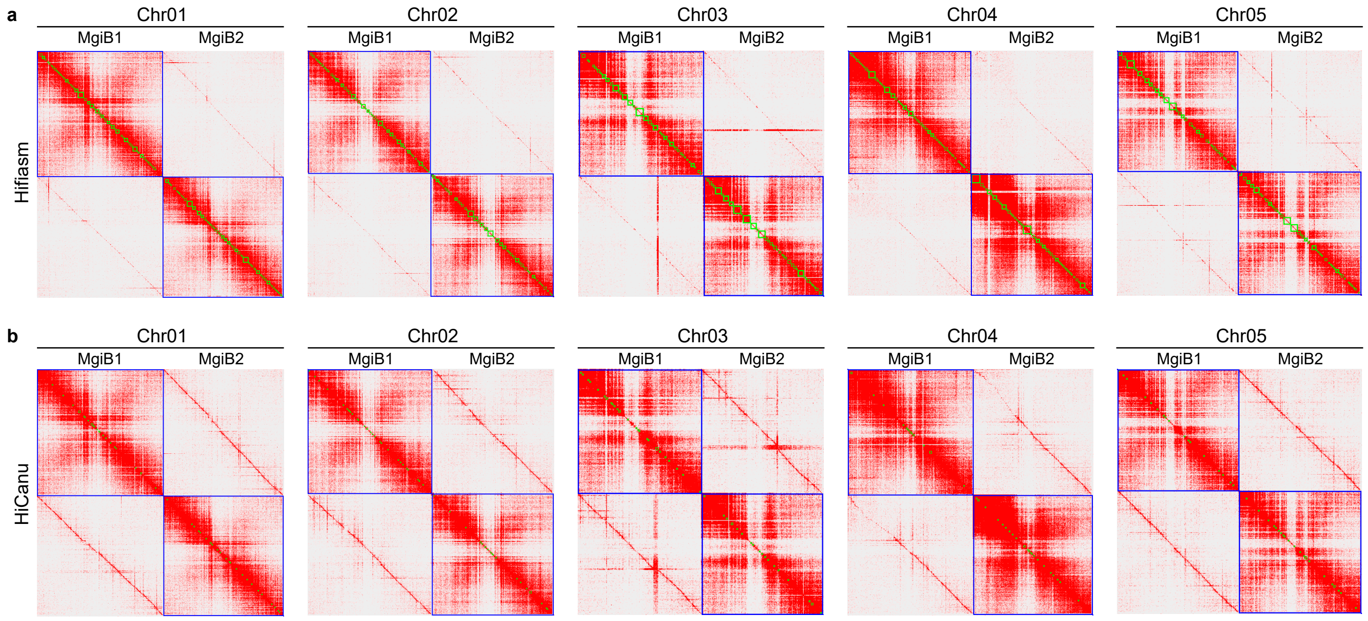


**Supplementary Fig. 29 |** Comparison of inter-allele Hi-C links between hifiasm and HiCanu assemblies. **a**, Hi-C contact maps of the *M.* × *giganteus* B subgenome (MgiB) assembled using Hifiasm. **b**, Hi-C contact maps of the MgiB assembled using HiCanu. Chromosomes 1-5 of MgiB are presented as examples.

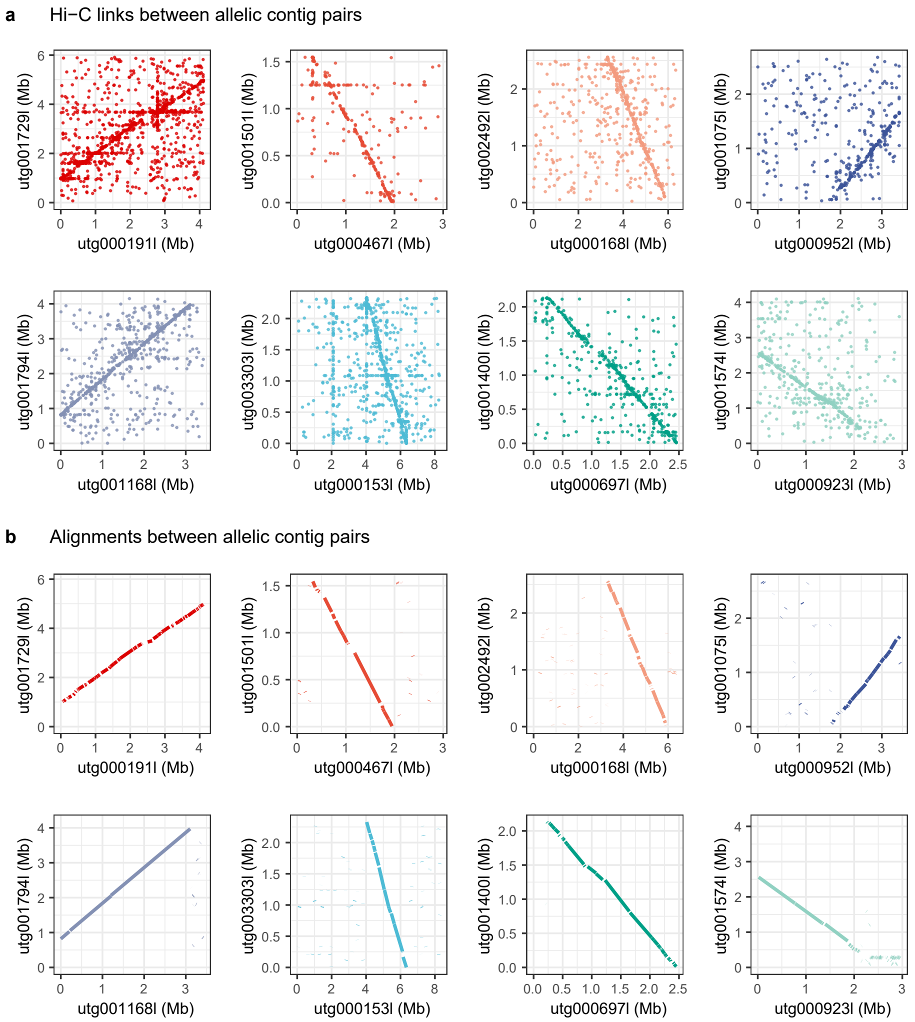


**Supplementary Fig. 30 |** Correlation between Hi-C links and alignments in allelic contig pairs. **a,** The Hi-C links between allelic contigs pairs. **b**, The alignments between allelic contig pairs. These examples of allelic contig pairs were selected from the hifiasm assembly of the *M.* × *giganteus* genome. The observed correlation suggests the potential utility of the distribution pattern of Hi-C links in identifying allelic contigs.

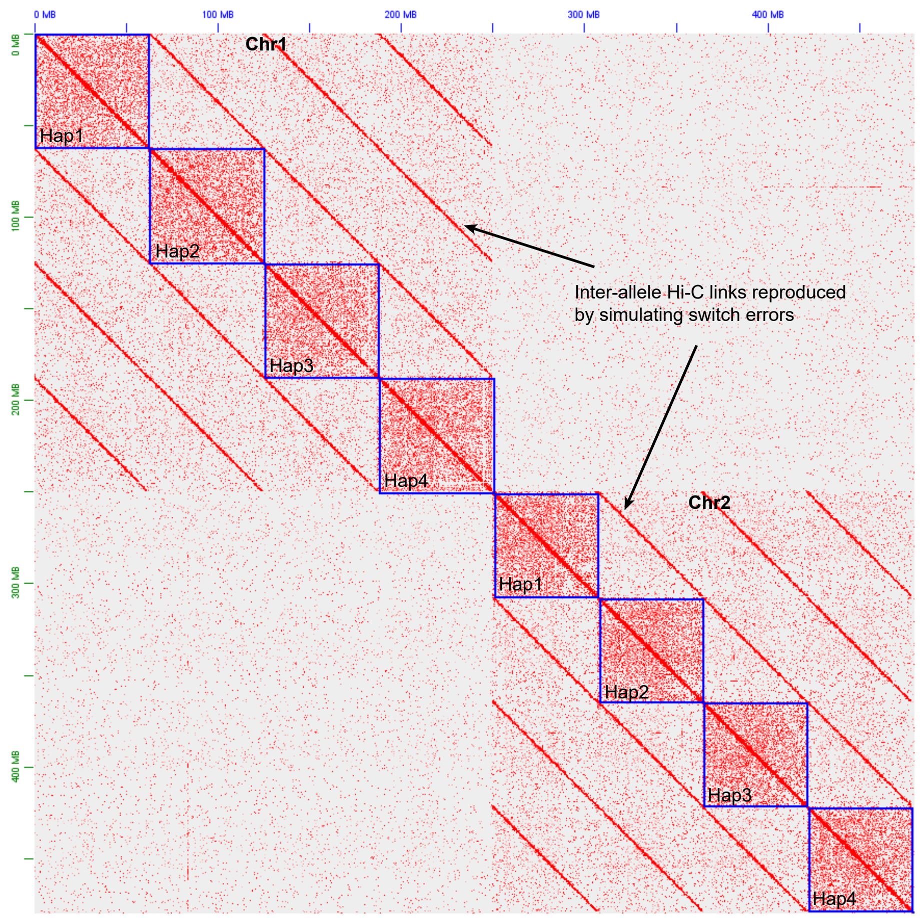


**Supplementary Fig. 31 |** Hi-C contact map of haplotypes with simulated switch errors. Arrows indicate inter-allele Hi-C links observed between haplotypes.

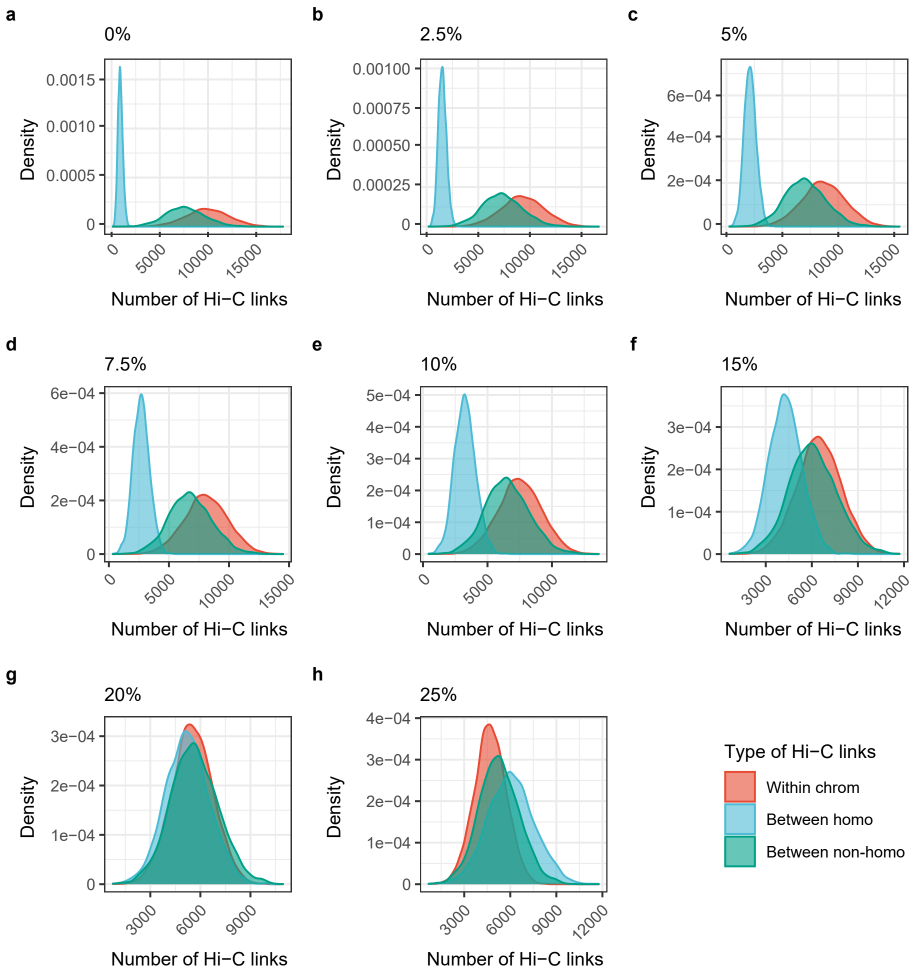


**Supplementary Fig. 32 |** Distribution of the number of Hi-C links for contigs with varying switch error rate ranging from 0 to 25% (**a**-**h**).

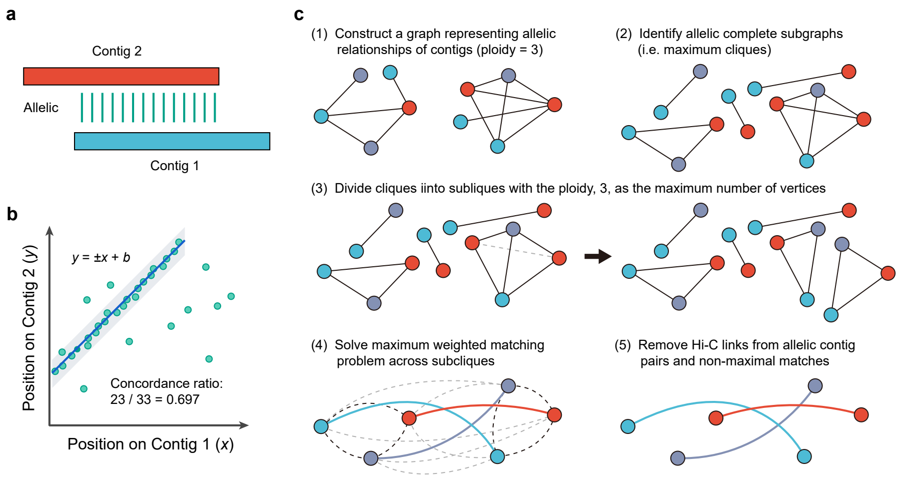


**Supplementary Fig. 33 |** Methodology for identifying allelic contigs and removing inter-allele Hi-C links. **a**, A schematic diagram illustrating the alignment between a pair of allelic contigs. **b**, The approach for calculating concordance ratios based on the distribution pattern of Hi-C links. **c**, The procedure for removing Hi-C links from allelic contig pairs and non-maximal matches.

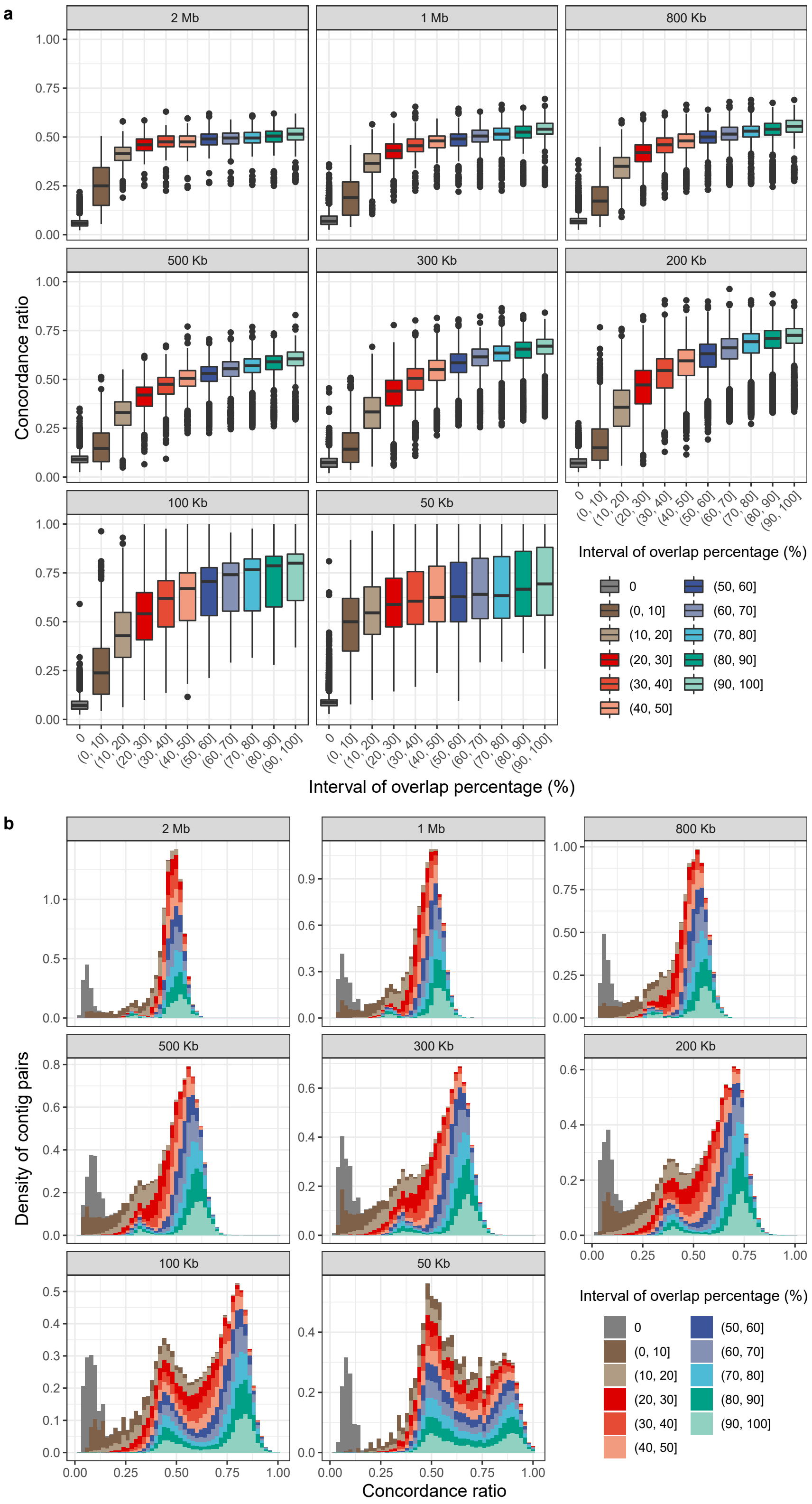


**Supplementary Fig. 34 |** Evaluation of the concordance ratio method in identifying allelic contig pairs. **a**, The concordance ratios increase with the percentage of overlap. **b**, The threshold demonstrates stable performance in detecting allelic contig pairs across varying contig lengths.

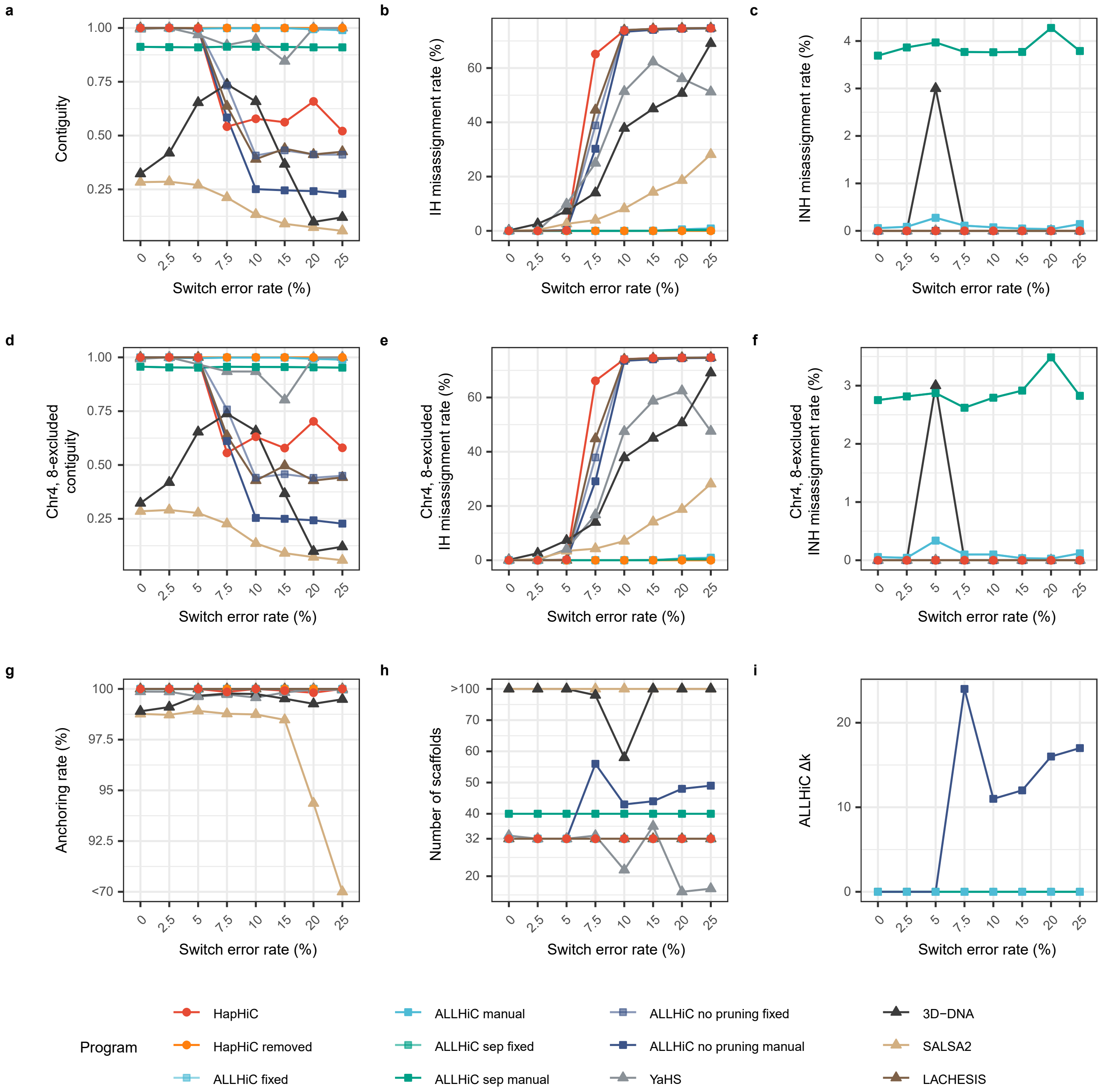


**Supplementary Fig. 35 |** Performance evaluation of Hi-C-based scaffolding tools in chromosome assignment on assemblies with varying switch error rates. The evaluation metrics include contiguity (**a**), misassignment rate between homologous chromosomes (**b**), misassignment rate between non-homologous chromosomes (**c**), anchoring rate (**g**), number of scaffolds (**h**), and $\Delta k$ of manual parameter tuning for ALLHiC (**i**). During ALLHiC pruning, the genome of a closely related species, *M. truncatula* (MtrunA17r5.0-ANR), was used as a reference. However, chromosomes 4 and 8 of *M. truncatula* have some structural differences compared to those of *M. sativa*. Therefore, we also calculated the contiguity and misassignment rate after excluding these two chromosomes (**d-f**). The parameters for these scaffolding tools are provided in Section **3 Hi-C-based scaffolding** of **Software Commands**.

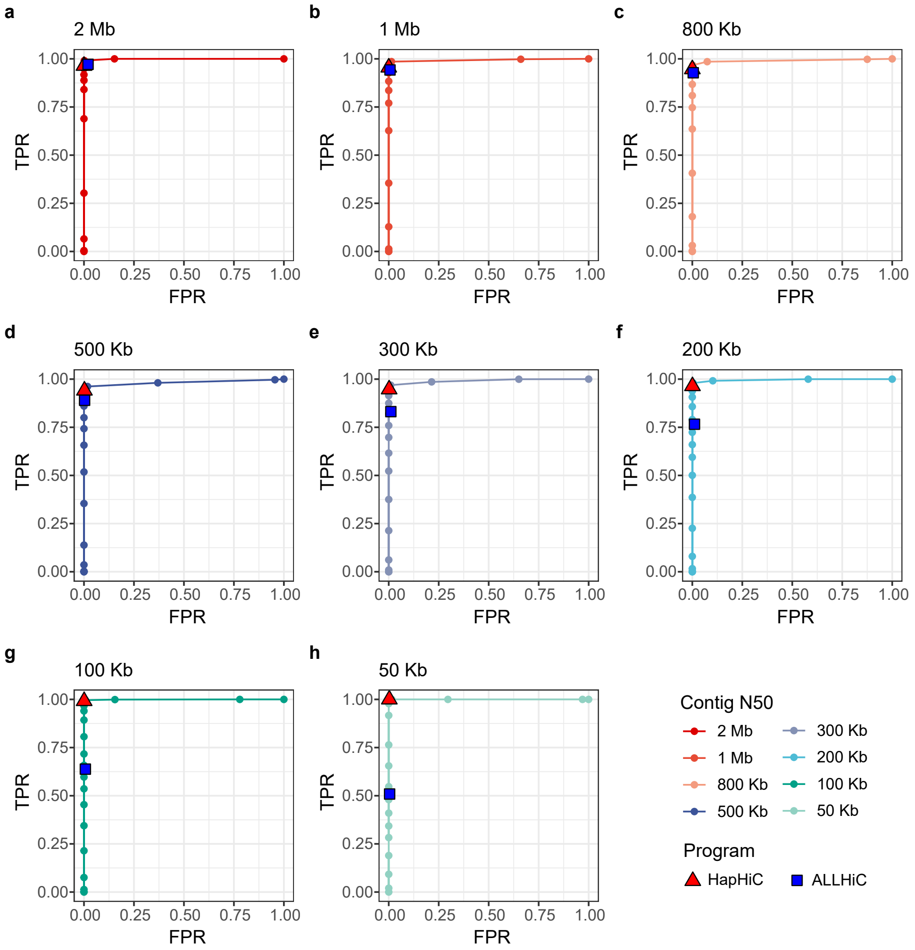


**Supplementary Fig. 36 |** Receiver operating characteristics (ROC) curves demonstrate the performance of the concordance ratio method in identifying allelic contig pairs of varying lengths ranging from 2 Mb to 50 Kb (**a**-**h**). Red triangles represent the performance of the default concordance ratio (0.2), while blue rectangles denote the performance of the reference-dependent method implemented in ALLHiC.

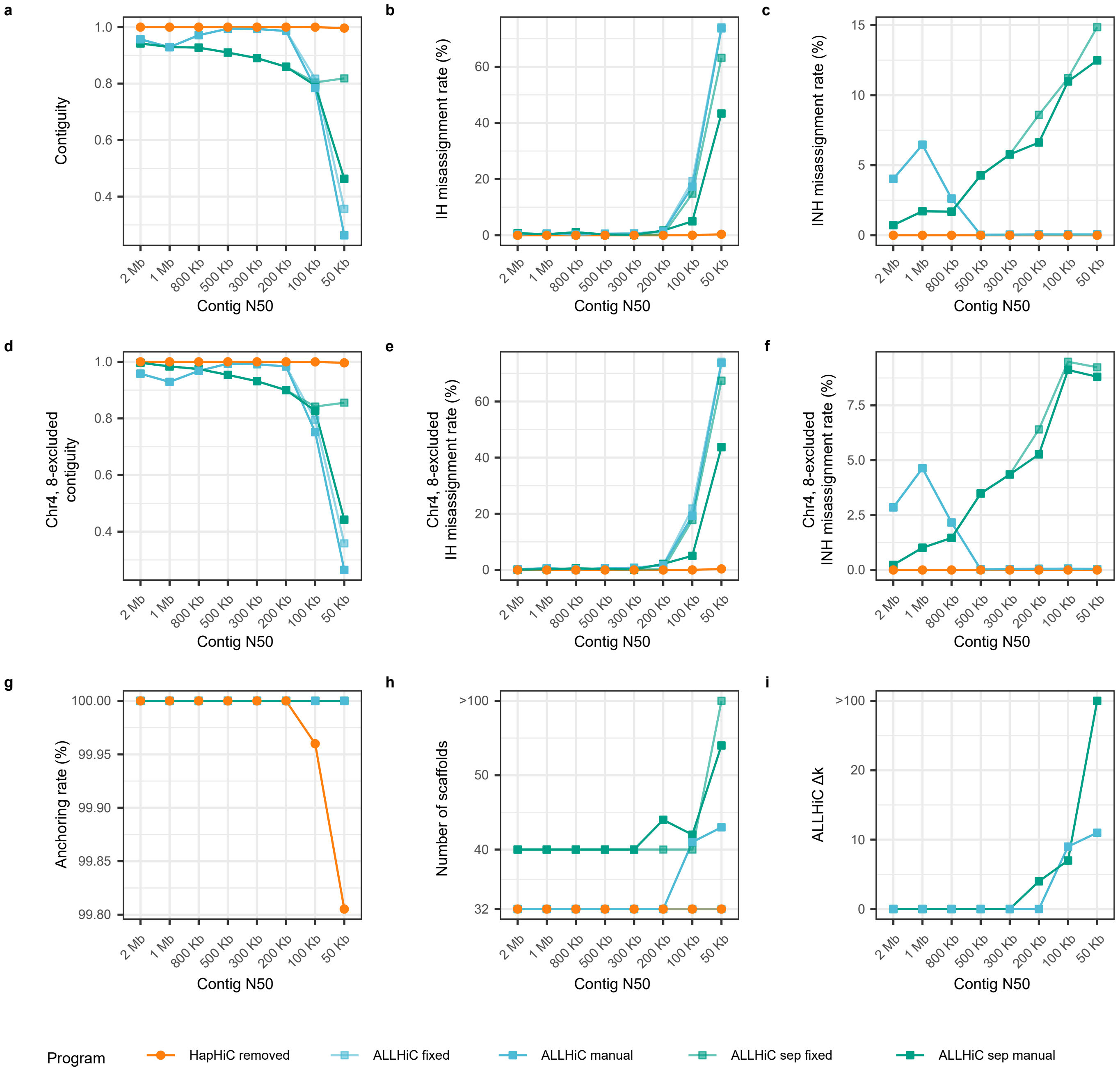


**Supplementary Fig. 37 |** Performance evaluation of Hi-C-based scaffolding tools in chromosome assignment on assemblies with identical switch error rate (20%) but varying contig N50 values. The evaluation metrics include contiguity (**a**), misassignment rate between homologous chromosomes (**b**), misassignment rate between non-homologous chromosomes (**c**), anchoring rate (**g**), number of scaffolds (**h**), and Δ*k* of manual parameter tuning for ALLHiC (**i**). During ALLHiC pruning, the genome of a closely related species, *M. truncatula* (MtrunA17r5.0-ANR), was used as a reference. However, chromosomes 4 and 8 of *M. truncatula* have some structural differences compared to those of *M. sativa*. Therefore, we also calculated the contiguity and misassignment rate after excluding these two chromosomes (**d-f**). The parameters for these scaffolding tools are provided in Section **3 Hi-C-based scaffolding** of **Software Commands**.

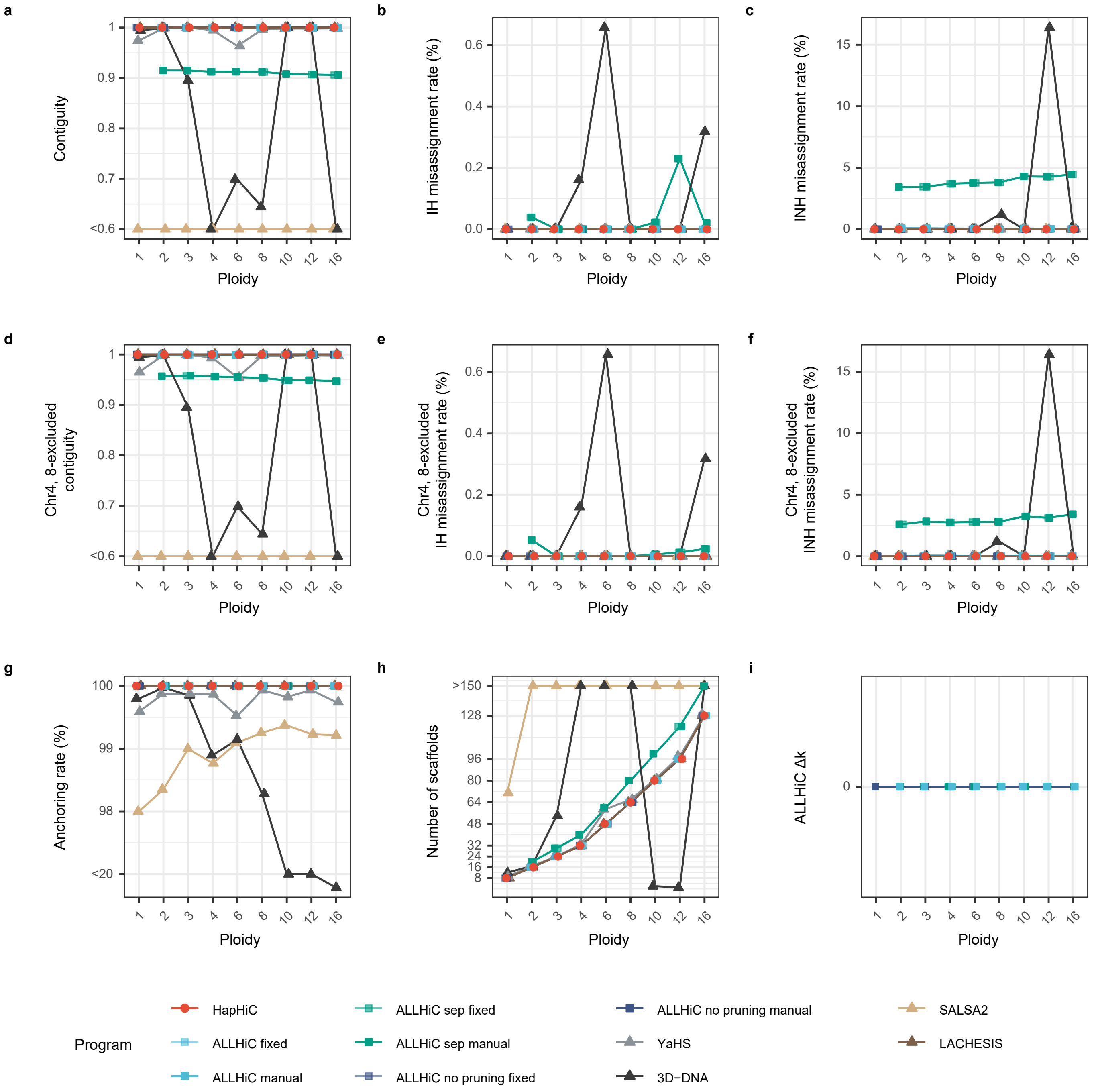


**Supplementary Fig. 38 |** Performance evaluation of Hi-C-based scaffolding tools in chromosome assignment on assemblies with varying ploidies. The evaluation metrics include contiguity (**a**), misassignment rate between homologous chromosomes (**b**), misassignment rate between non-homologous chromosomes (**c**), anchoring rate (**g**), number of scaffolds (**h**), and $\Delta k$ of manual parameter tuning for ALLHiC (**i**). During ALLHiC pruning, the genome of a closely related species, *M. truncatula* (MtrunA17r5.0-ANR), was used as a reference. However, chromosomes 4 and 8 of *M. truncatula* have some structural differences compared to those of *M. sativa*. Therefore, we also calculated the contiguity and misassignment rate after excluding these two chromosomes (**d**-**f**). The parameters for these scaffolding tools are provided in Section **3 Hi-C-based scaffolding** of **Software Commands**.

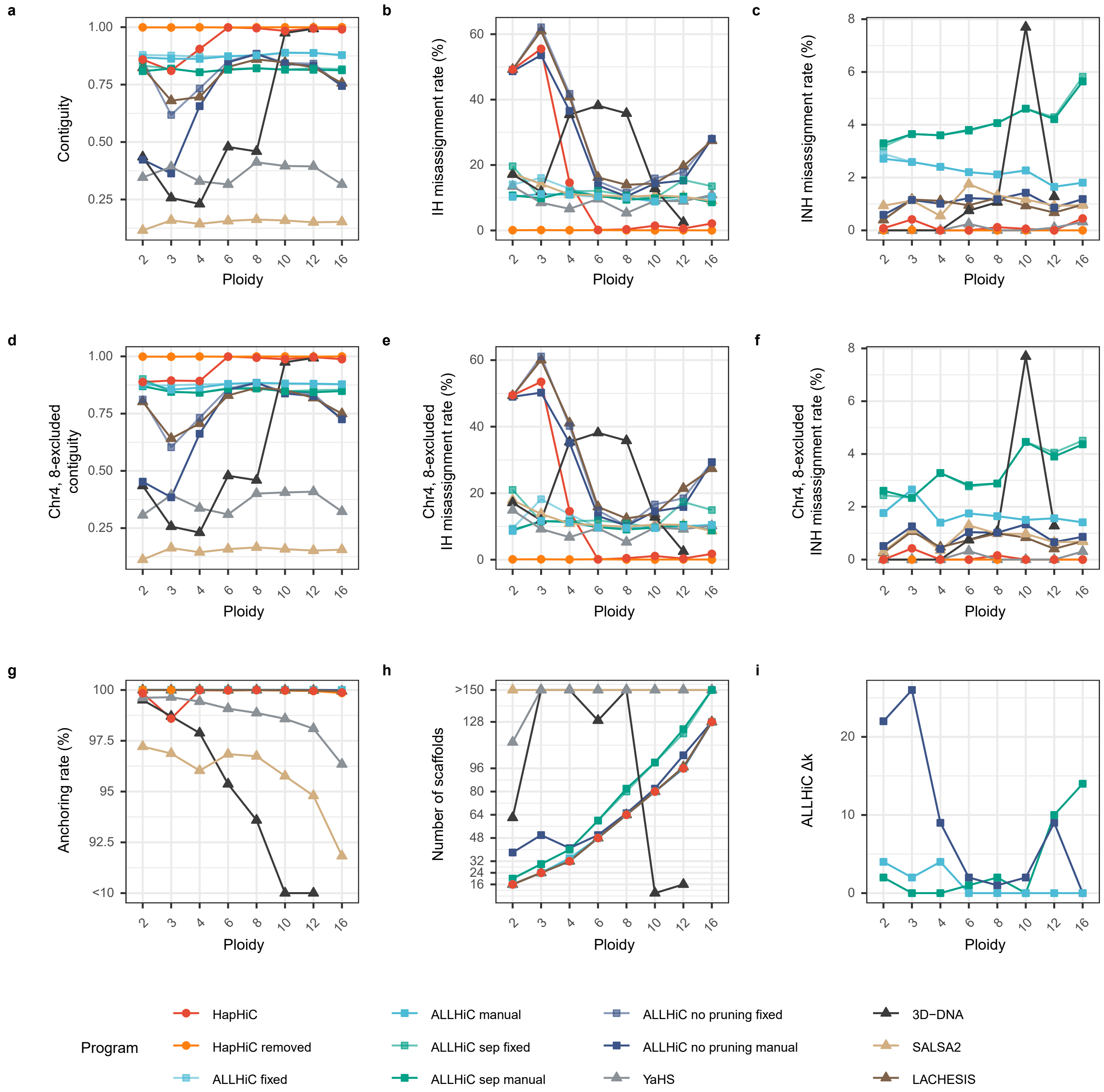


**Supplementary Fig. 39 |** Performance evaluation of Hi-C-based scaffolding tools in chromosome assignment on assemblies with varying ploidies and 5% each of chimeric, collapsed, and switch errors. The evaluation metrics include contiguity (**a**), misassignment rate between homologous chromosomes (**b**), misassignment rate between non-homologous chromosomes (**c**), anchoring rate (**g**), number of scaffolds (**h**), and $\Delta k$ of manual parameter tuning for ALLHiC (**i**). During ALLHiC pruning, the genome of a closely related species, *M. truncatula* (MtrunA17r5.0-ANR), was used as a reference. However, chromosomes 4 and 8 of *M. truncatula* have some structural differences compared to those of *M. sativa*. Therefore, we also calculated the contiguity and misassignment rate after excluding these two chromosomes (**d-f**). The parameters for these scaffolding tools are provided in Section **3 Hi-C-based scaffolding** of **Software Commands**.

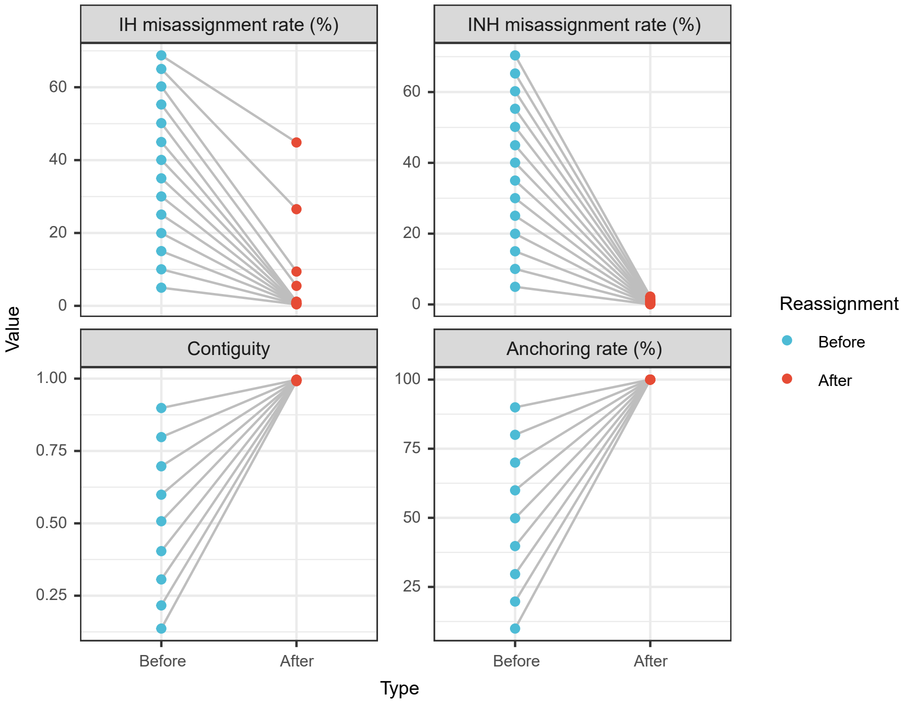


**Supplementary Fig. 40 |** Performance evaluation of the reassignment process in HapHiC. The evaluation metrics include misassignment rate between homologous chromosomes (IH misassignment rate), misassignment rate between non-homologous chromosomes (INH misassignment rate), contiguity, and anchoring rate.

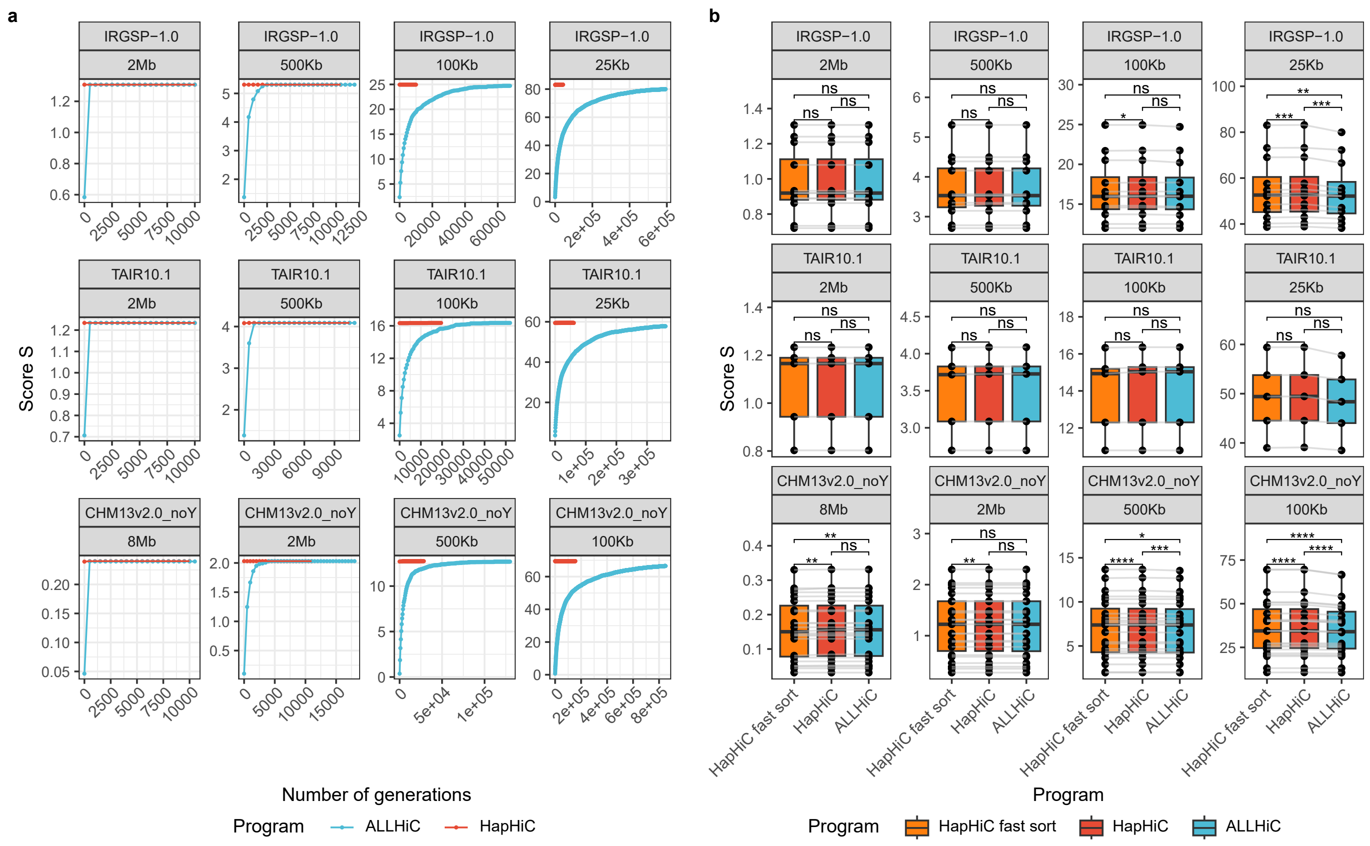


**Supplementary Fig. 41 |** Performance evaluation of HapHiC and ALLHiC in contig ordering and orientation using the built-in scoring system of ALLHiC. This evaluation was conducted on simulated assemblies of rice IRGSP-1.0, *Arabidopsis* TAIR10.1, and human CHM13v2.0_noY genomes with varying contig N50 values. **a**, The scores increase with the number of generations, with HapHiC demonstrating a significant decrease in the iterations needed for the genetic algorithm to converge. **b**, A comparison of the final scores between HapHiC and ALLHiC. When only fast sorting was employed, the initial scores of HapHiC were already comparable to or even higher than the final scores achieved by ALLHiC. In assemblies with low contig contiguity, these scores were further improved during the subsequent optimization process. P values were derived from two-sided Wilcoxon signed-rank tests ($n$ = 12, 5, 23 for IRGSP-1.0, TAIR10.1, and CHM13v2.0_noY, respectively).

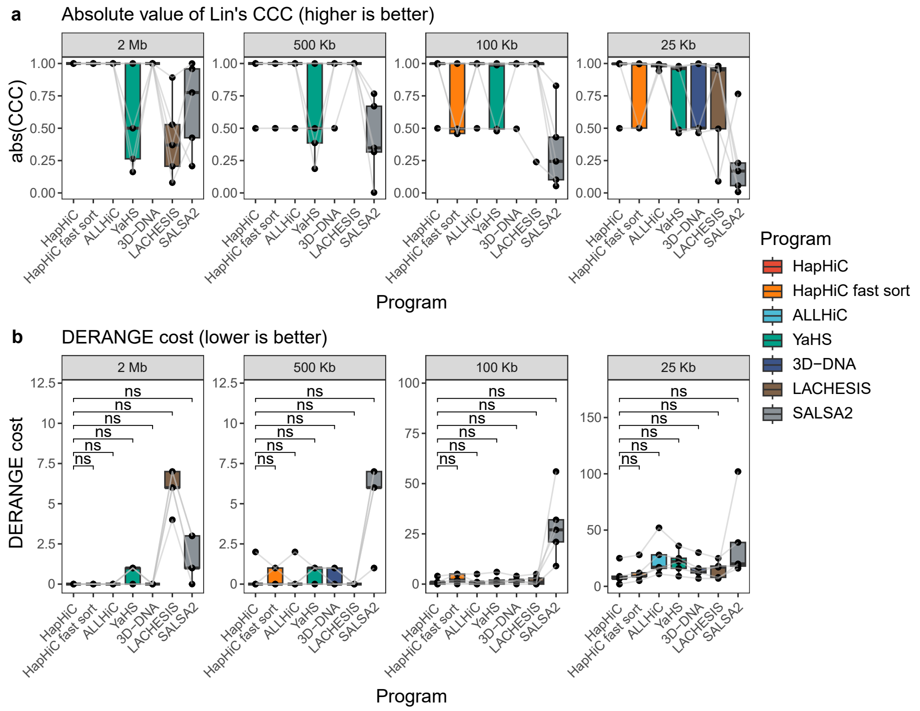


**Supplementary Fig. 42 |** Performance of each Hi-C-based scaffolding tools in ordering and orienting the contigs of the *Arabidopsis* TAIR10.1 chromosomes with varying contig N50 values. **a**, The absolute values of Lin's concordance correlation coefficients (CCCs) for each Hi-C-based scaffolding tool. **b**, The DERANGE costs for each Hi-C-based scaffolding tool ($p$ values from two-sided Wilcoxon signed-rank tests, $n = 5$).

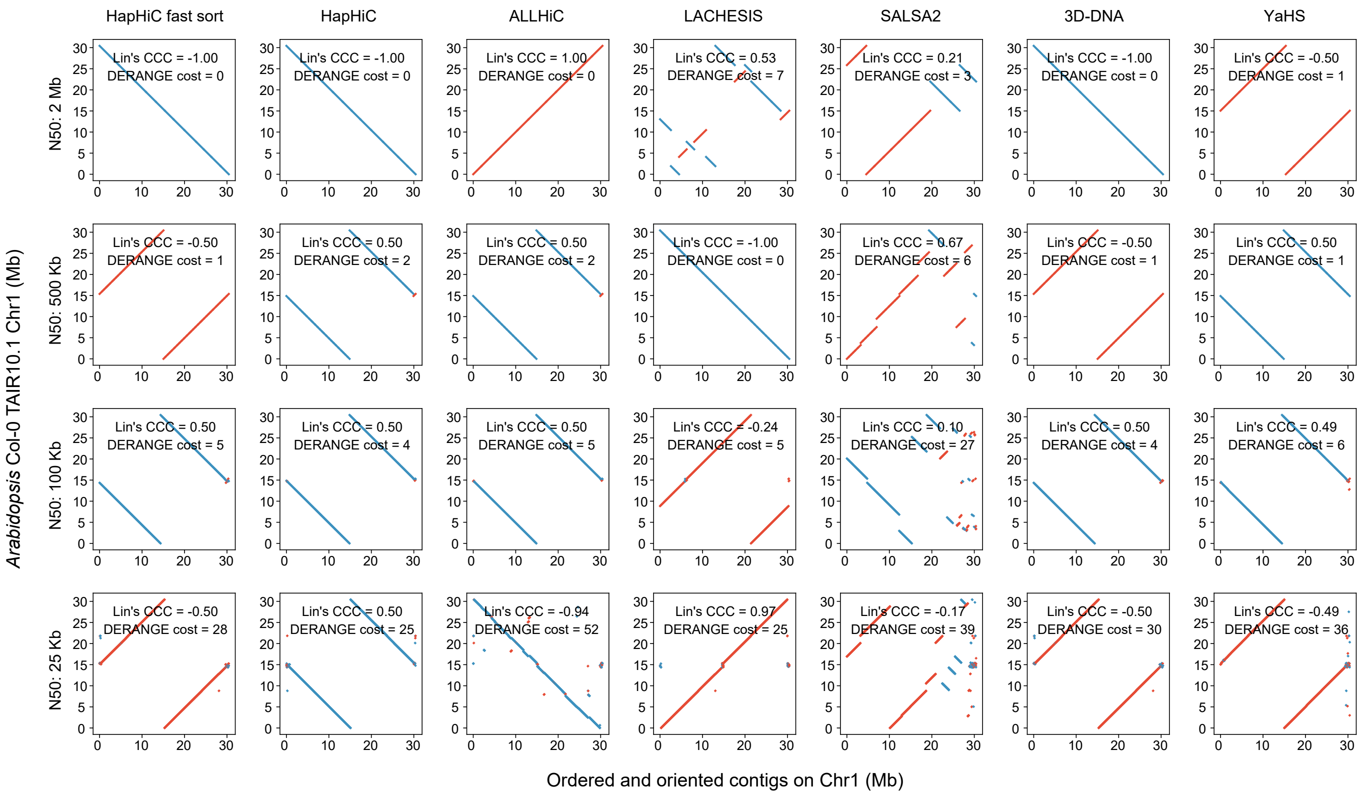


**Supplementary Fig. 43 |** Dot plots illustrate the concordance between chromosome 1 of the *Arabidopsis* TAIR10.1 genome and the contig ordering and orientation result produced by each Hi-C-based scaffolding tool. Red and blue dots represent forward and reverse complementary alignments, respectively. The corresponding Lin's concordance correlation coefficients (CCCs) and DERANGE costs are also displayed in these dot plots.

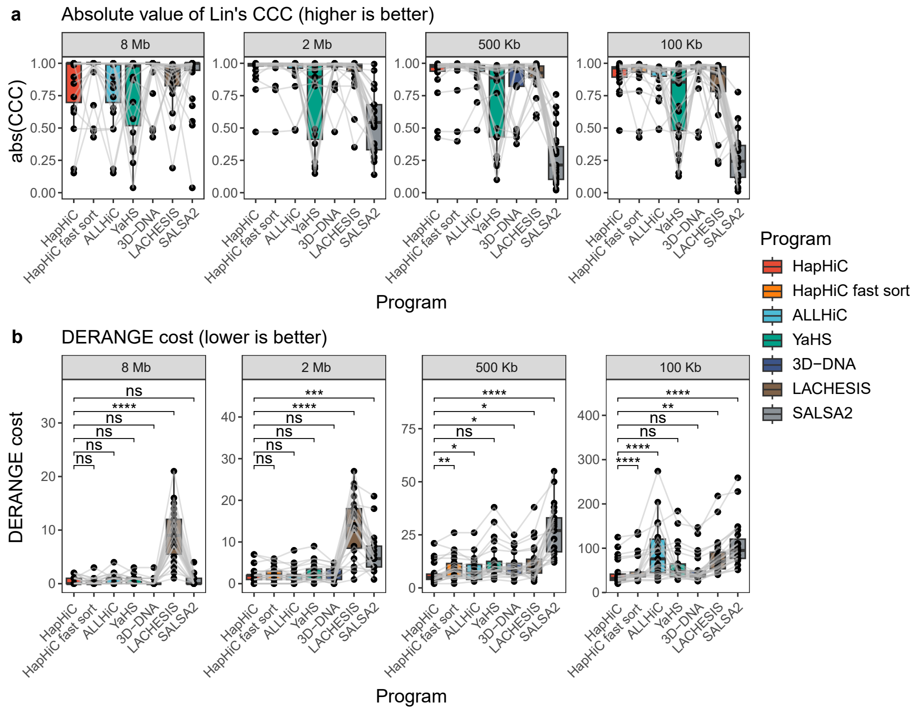


**Supplementary Fig. 44 |** Performance of each Hi-C-based scaffolding tools in ordering and orienting the contigs of the human CHM13v2.0_noY chromosomes with varying contig N50 values. **a**, The absolute values of Lin's concordance correlation coefficients (CCCs) for each Hi-C-based scaffolding tool. **b**, The DERANGE costs for each Hi-C-based scaffolding tool ($p$ values from two-sided Wilcoxon signed-rank tests, $n$ = 23).

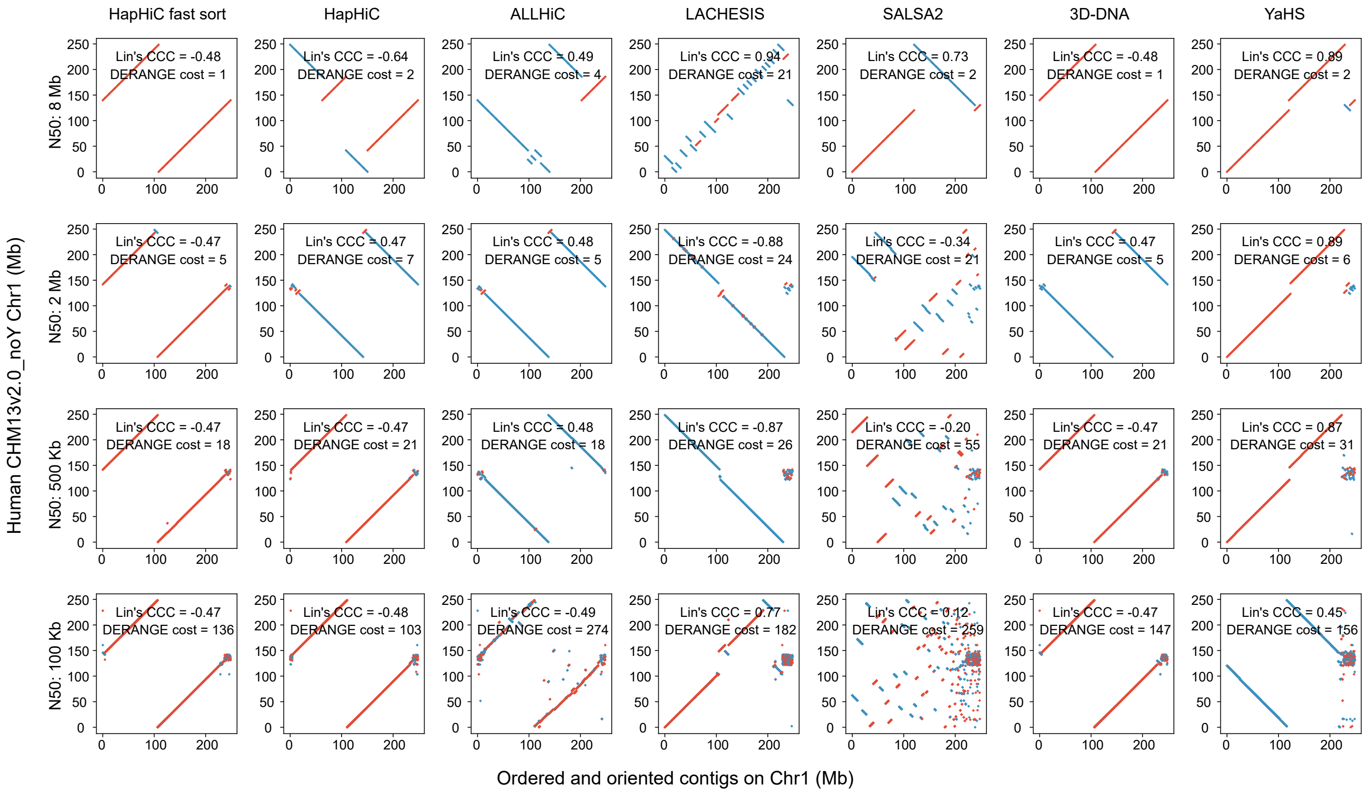


**Supplementary Fig. 45 |** Dot plots illustrate the concordance between chromosome 1 of the human CHM13v2.0_noY genome and the contig ordering and orientation result produced by each Hi-C-based scaffolding tool. Red and blue dots represent forward and reverse complementary alignments, respectively. The corresponding Lin's concordance correlation coefficients (CCCs) and DERANGE costs are also displayed in these dot plots.

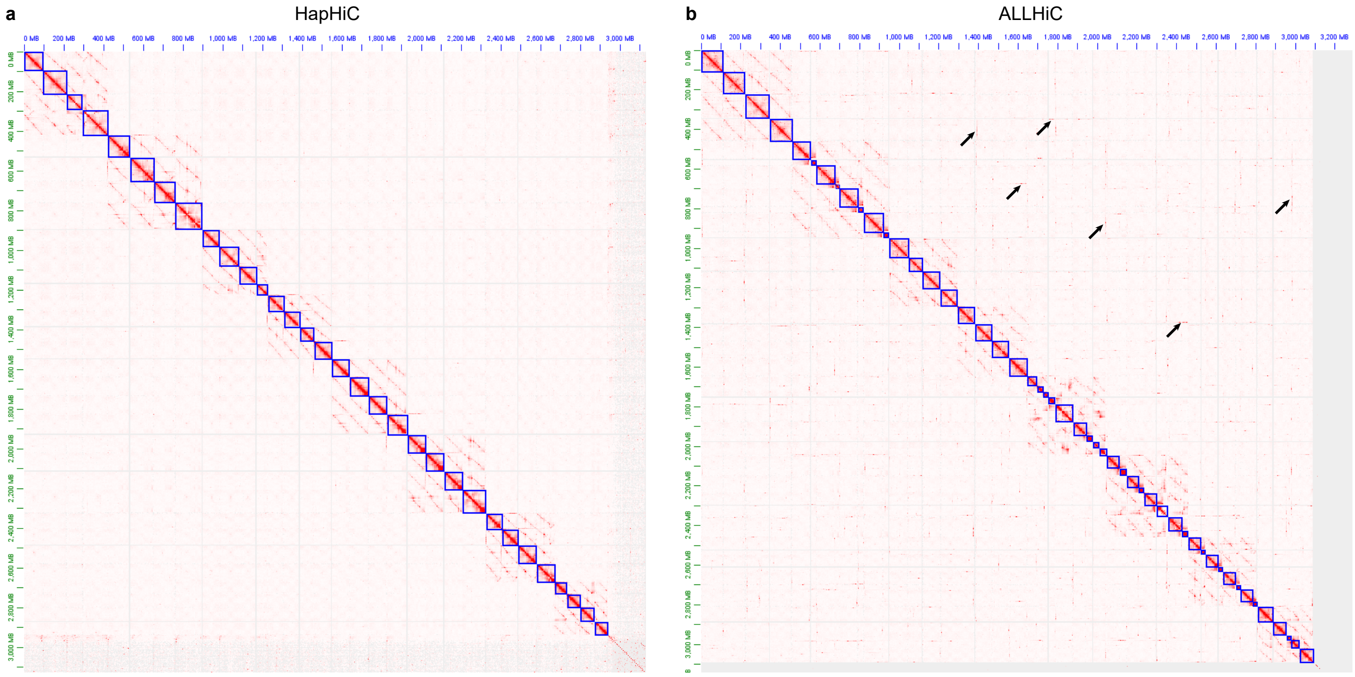


**Supplementary Fig. 46 |** Hi-C contact maps of HapHiC (**a**) and ALLHiC (**b**) scaffolds of the *S. spontaneum* AP85-441 genome. Black arrows indicate examples of misassignments between non-homologous chromosomes in the ALLHiC scaffolds.

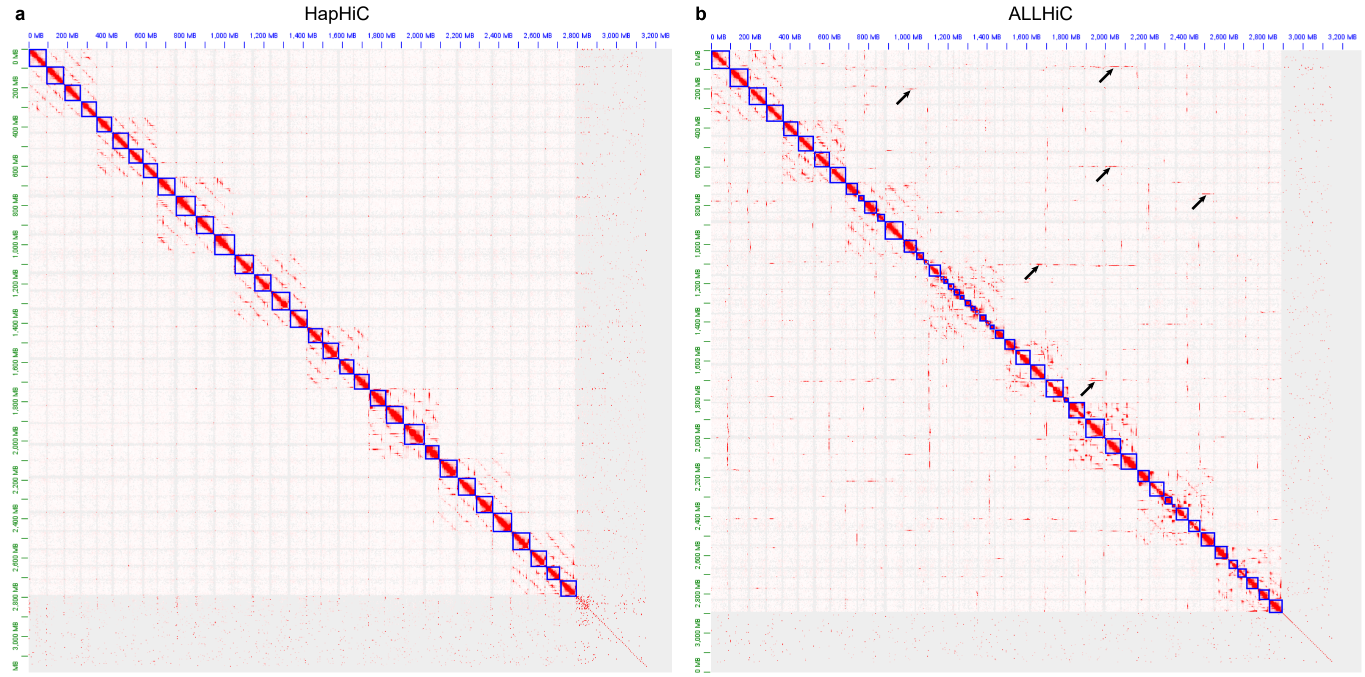


**Supplementary Fig. 47 |** Hi-C contact maps of HapHiC (**a**) and ALLHiC (**b**) scaffolds of the *M. sativa* XinJiangDaYe genome. Black arrows indicate examples of misassignments between non-homologous chromosomes in the ALLHiC scaffolds.

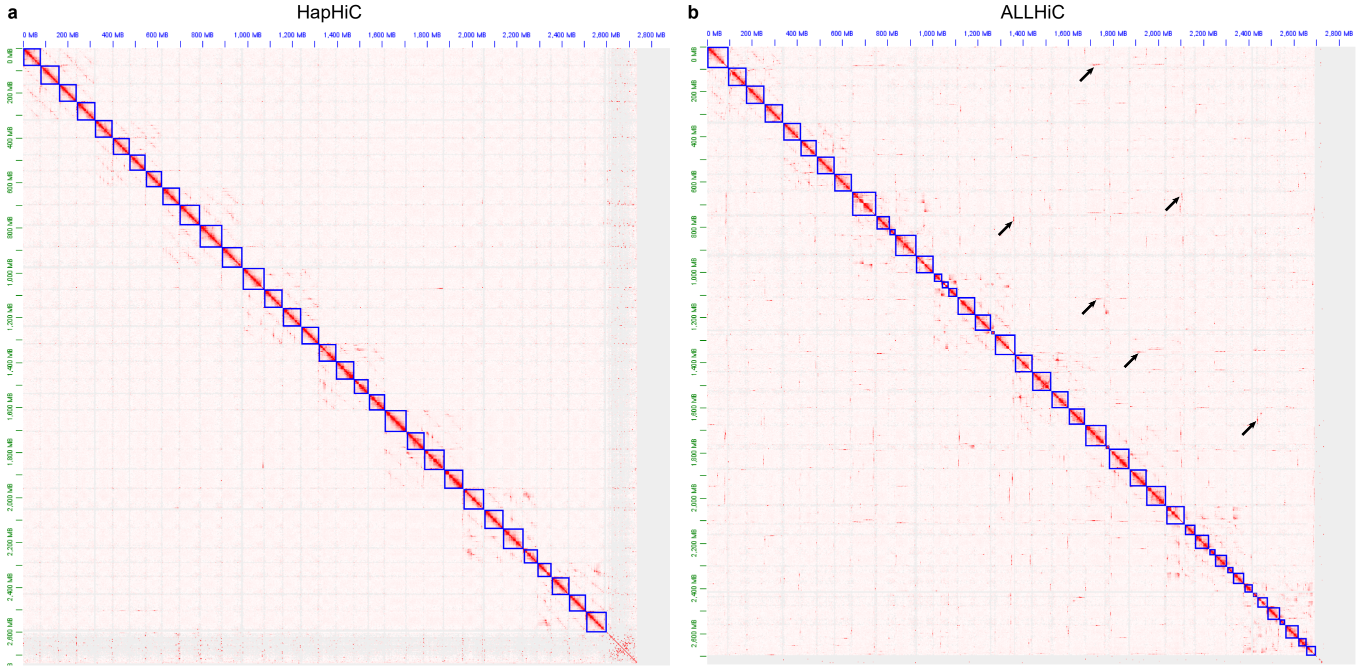


**Supplementary Fig. 48 |** Hi-C contact maps of HapHiC (**a**) and ALLHiC (**b**) scaffolds of the *M. sativa* Zhongmu-4 genome. Black arrows indicate examples of misassignments between non-homologous chromosomes in the ALLHiC scaffolds.

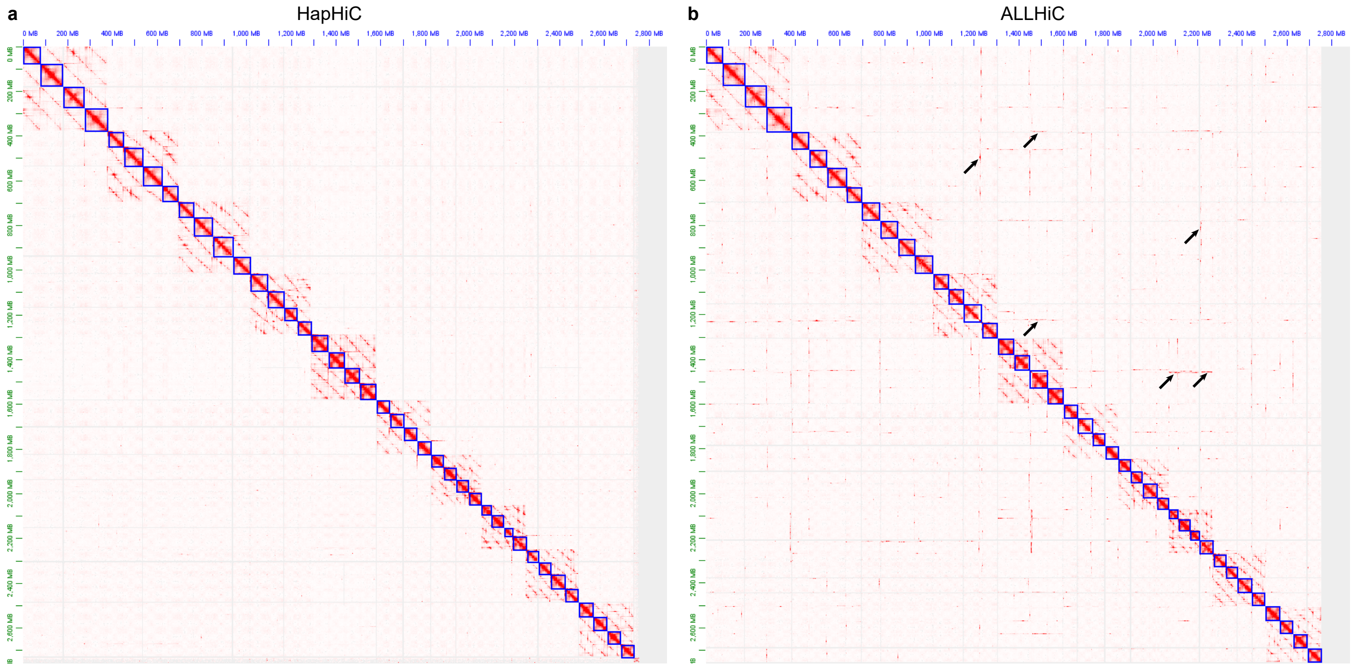


**Supplementary Fig. 49 |** Hi-C contact maps of HapHiC (**a**) and ALLHiC (**b**) scaffolds of the *S. spontaneum* Np-X genome. Black arrows indicate examples of misassignments between non-homologous chromosomes in the ALLHiC scaffolds.

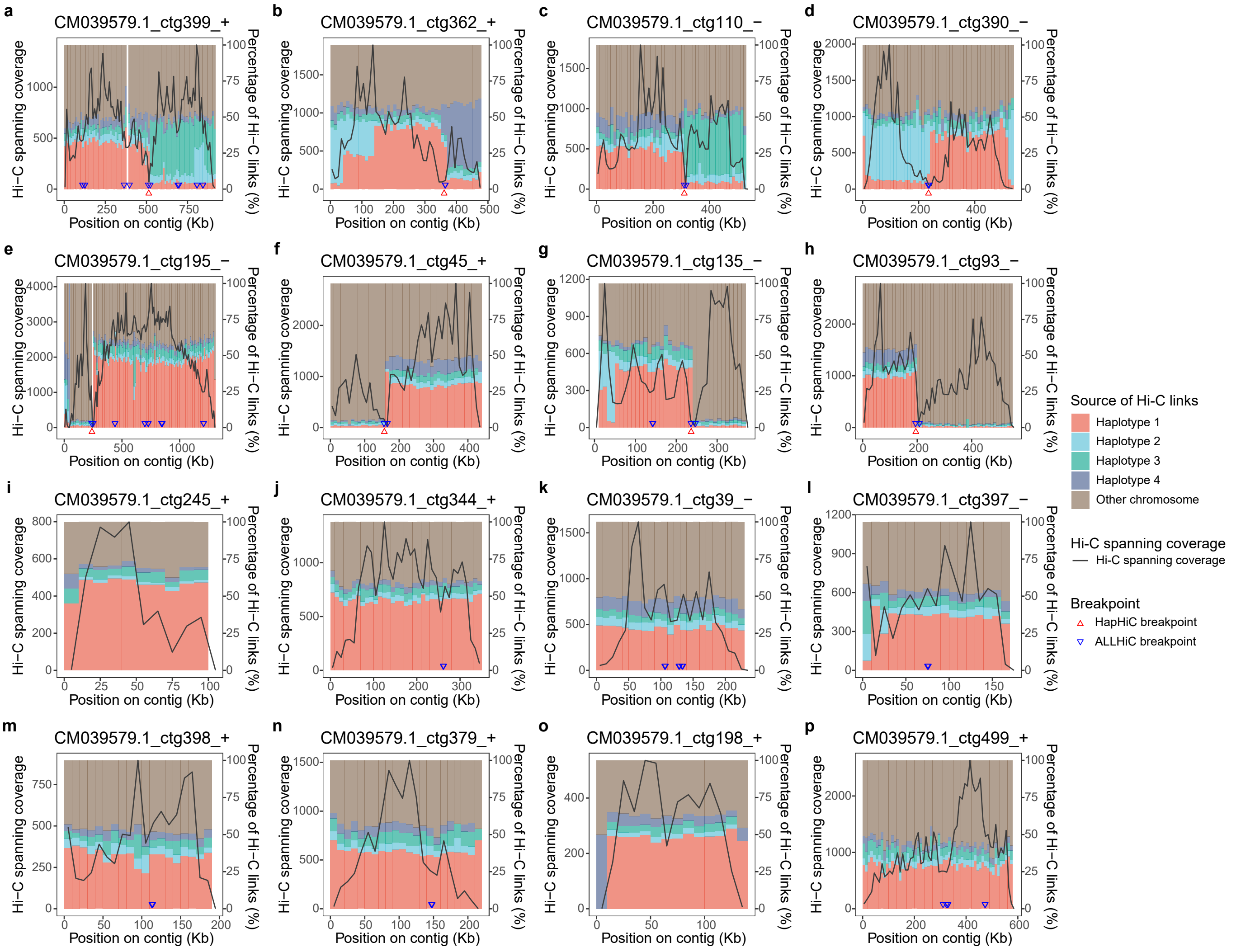


**Supplementary Fig. 50 |** Examples of assembly correction made by HapHiC and ALLHiC. These examples include four chimeric contigs formed between homologous chromosomes (**a-d**), four chimeric contigs formed between non-homologous chromosomes (**e-h**), and eight non-chimeric contigs (**i-p**). The line charts represent the Hi-C spanning coverages along contigs (left axes), while histograms depict the percentages of Hi-C links based on their sources along contigs (right axes). The source of each Hi-C link is determined by the mapping position of the other end of the read pair. Red and blue triangles indicate the breakpoints determined by HapHiC and ALLHiC, respectively.

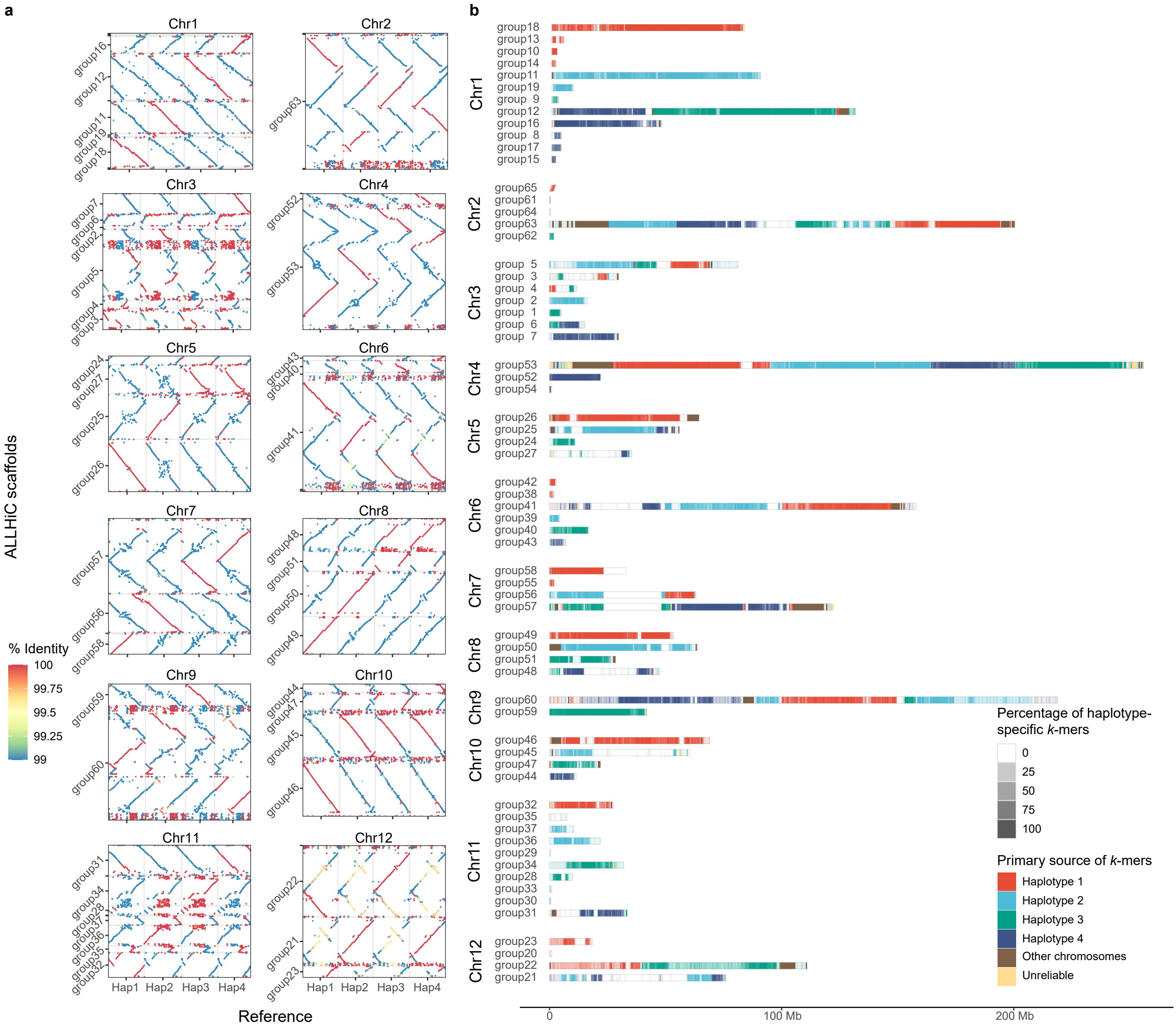

**Supplementary Fig. 51 |** Performance evaluation of ALLHiC in scaffolding the haplotypes of potato C88 genome. **a**, The dot plots illustrate the alignments between the ALLHiC scaffolds and the haplotypes of potato C88 genome, with dot colors indicating the sequence identities of alignments. **b**, A *k*-mer-based analysis reveals the primary source of each position along the contigs from the potato C88 haplotypes. The color indicates the primary source of *k*-mers, while the degree of transparency represents the percentage of haplotype-specific *k*-mers.

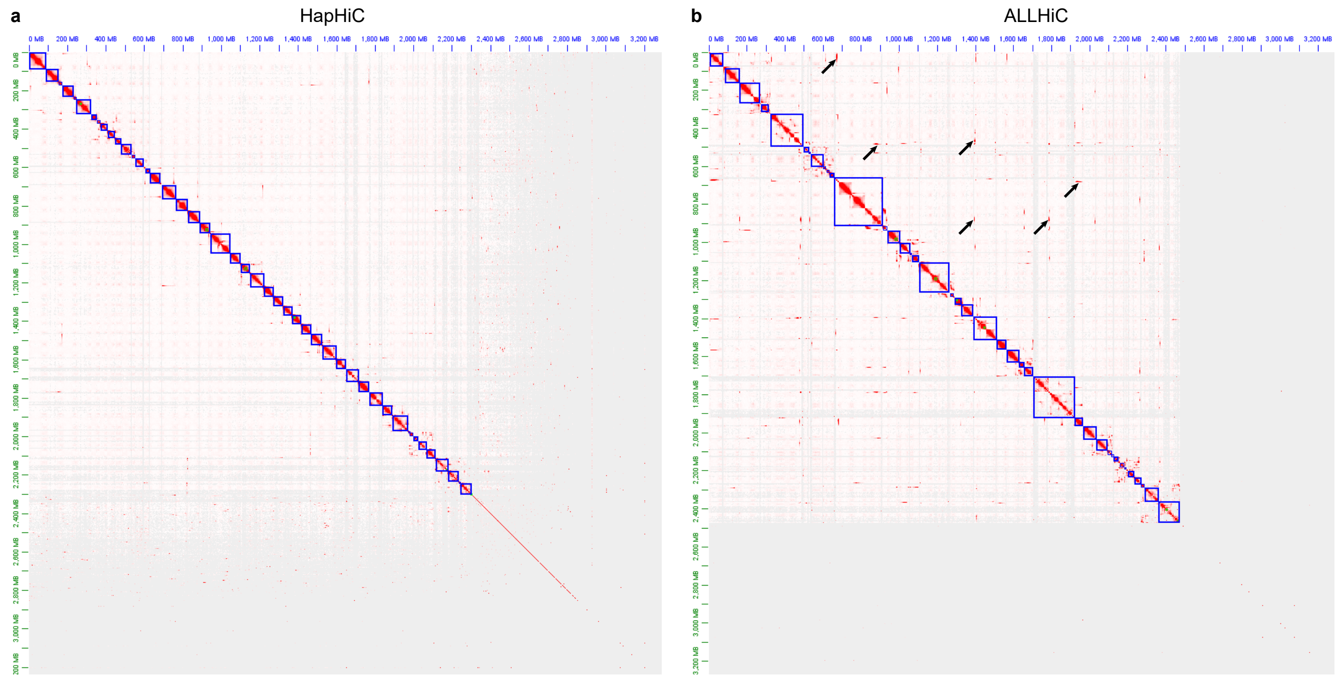


**Supplementary Fig. 52 |** Hi-C contact maps of HapHiC (**a**) and ALLHiC (**b**) scaffolds of the potato C88 genome. Black arrows indicate examples of misassignments between non-homologous chromosomes in the ALLHiC scaffolds.

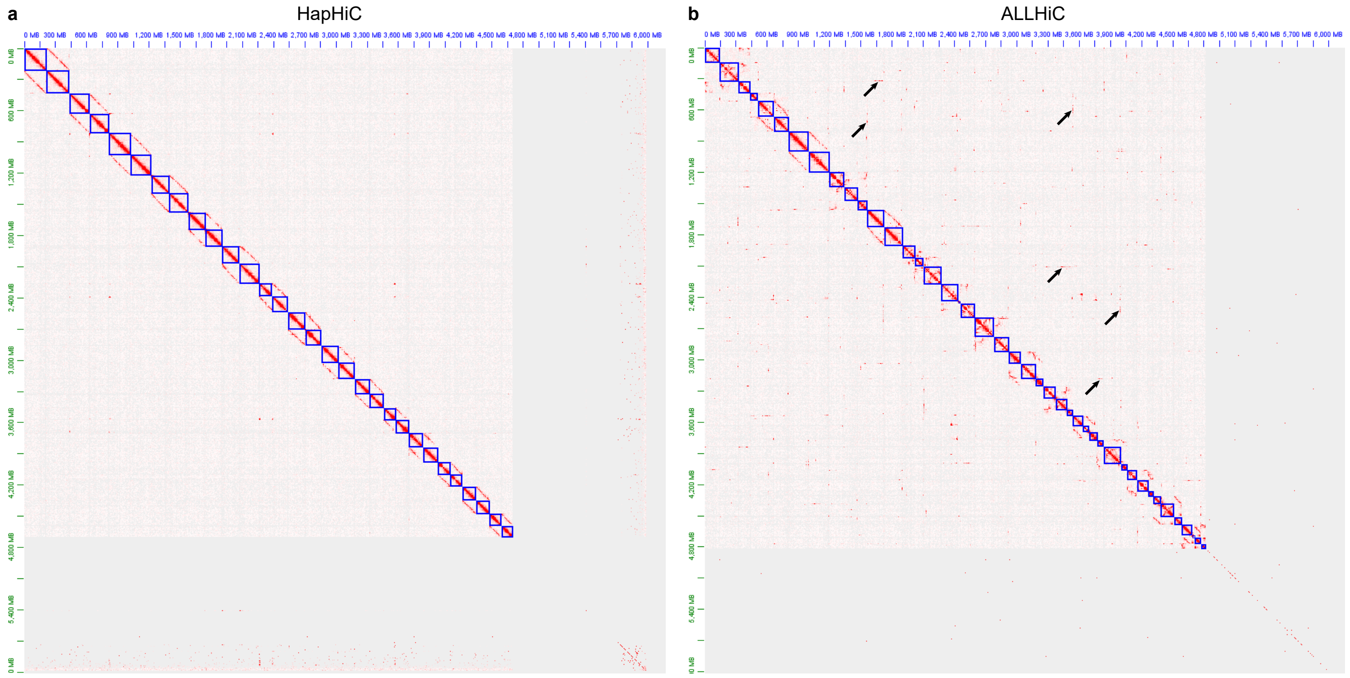


**Supplementary Fig. 53 |** Hi-C contact maps of HapHiC (**a**) and ALLHiC (**b**) scaffolds of the *C. sinensis* Tieguanyin genome. Black arrows indicate examples of misassignments between non-homologous chromosomes in the ALLHiC scaffolds.

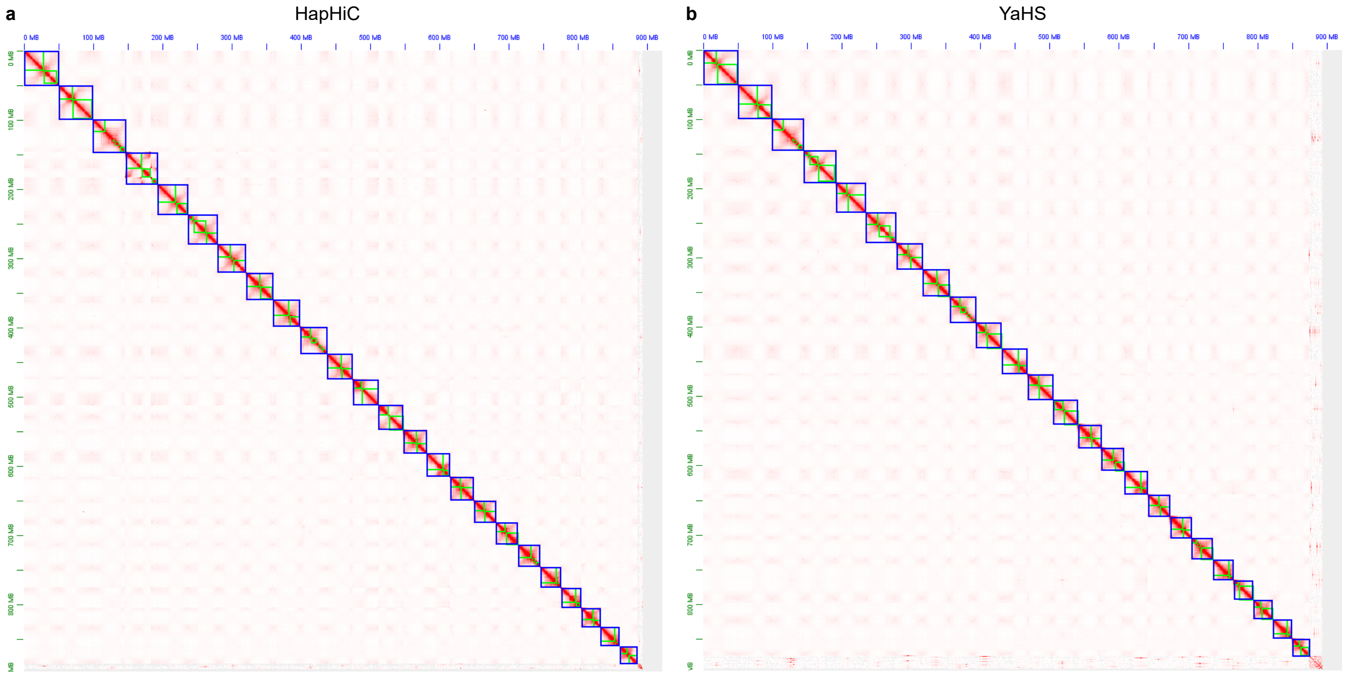


**Supplementary Fig. 54 |** Hi-C contact maps of HapHiC (**a**) and YaHS (**b**) scaffolds of the *Oryza alta* genome.

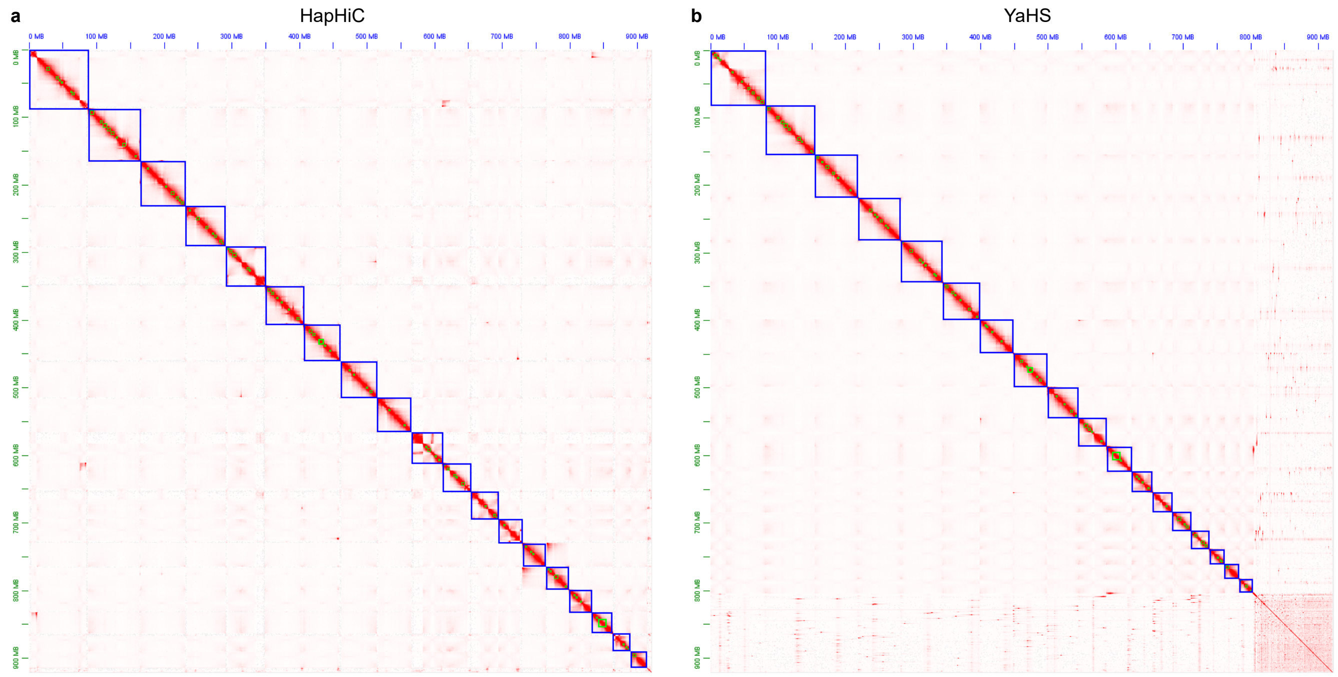


**Supplementary Fig. 55 |** Hi-C contact maps of HapHiC (**a**) and YaHS (**b**) scaffolds of the *Brassica napus* genome.

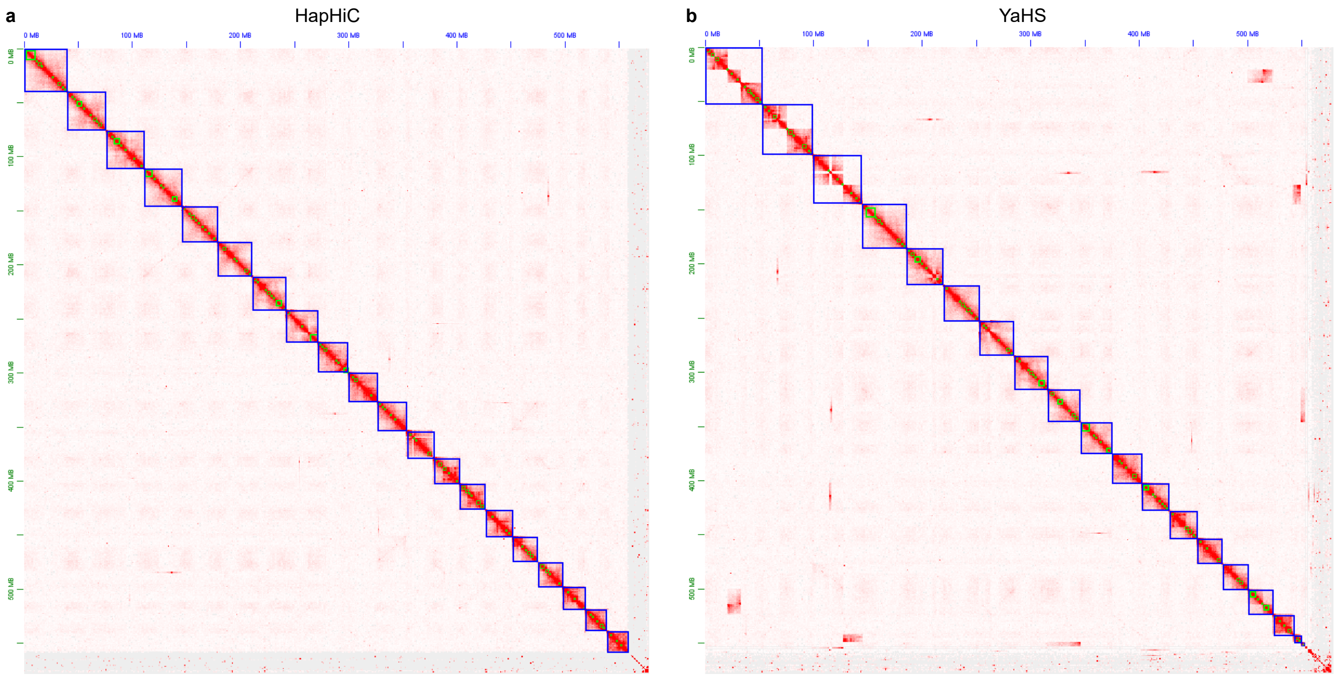


**Supplementary Fig. 56 |** Hi-C contact maps of HapHiC (**a**) and YaHS (**b**) scaffolds of the *Eragrostis tef* genome.

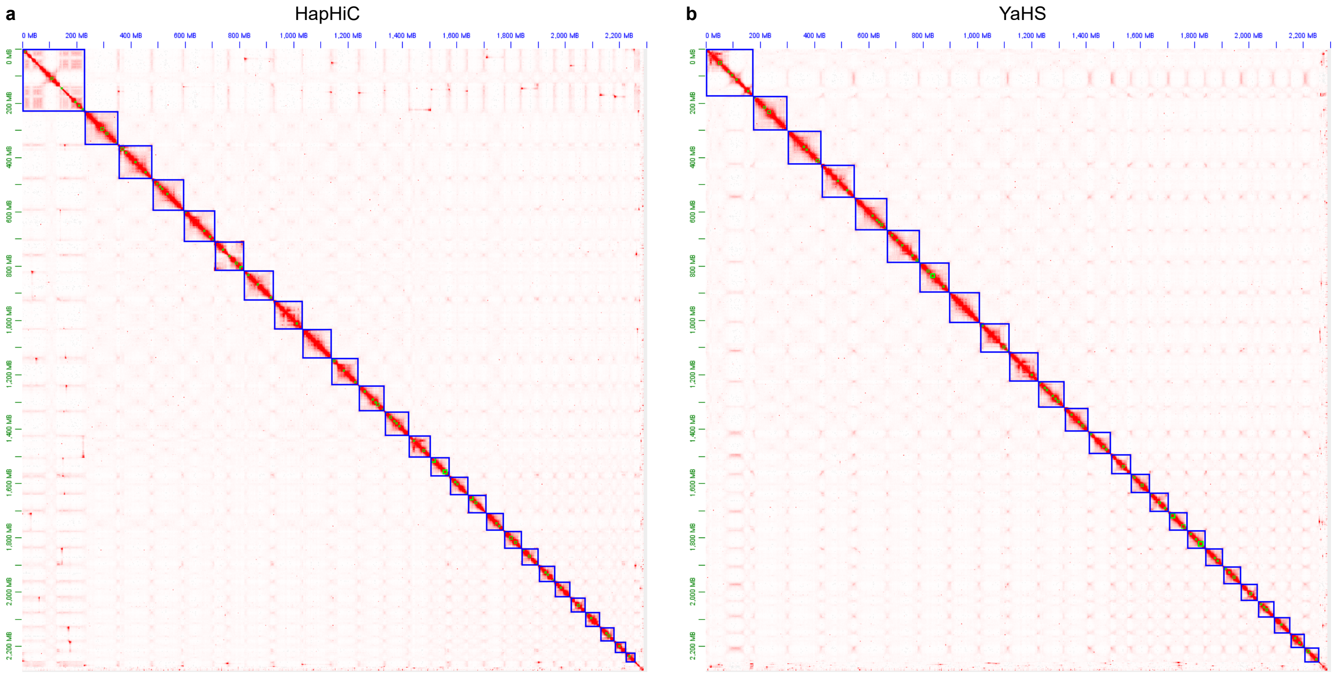


**Supplementary Fig. 57 |** Hi-C contact maps of HapHiC (**a**) and YaHS (**b**) scaffolds of the *Gossypium hirsutum* genome.

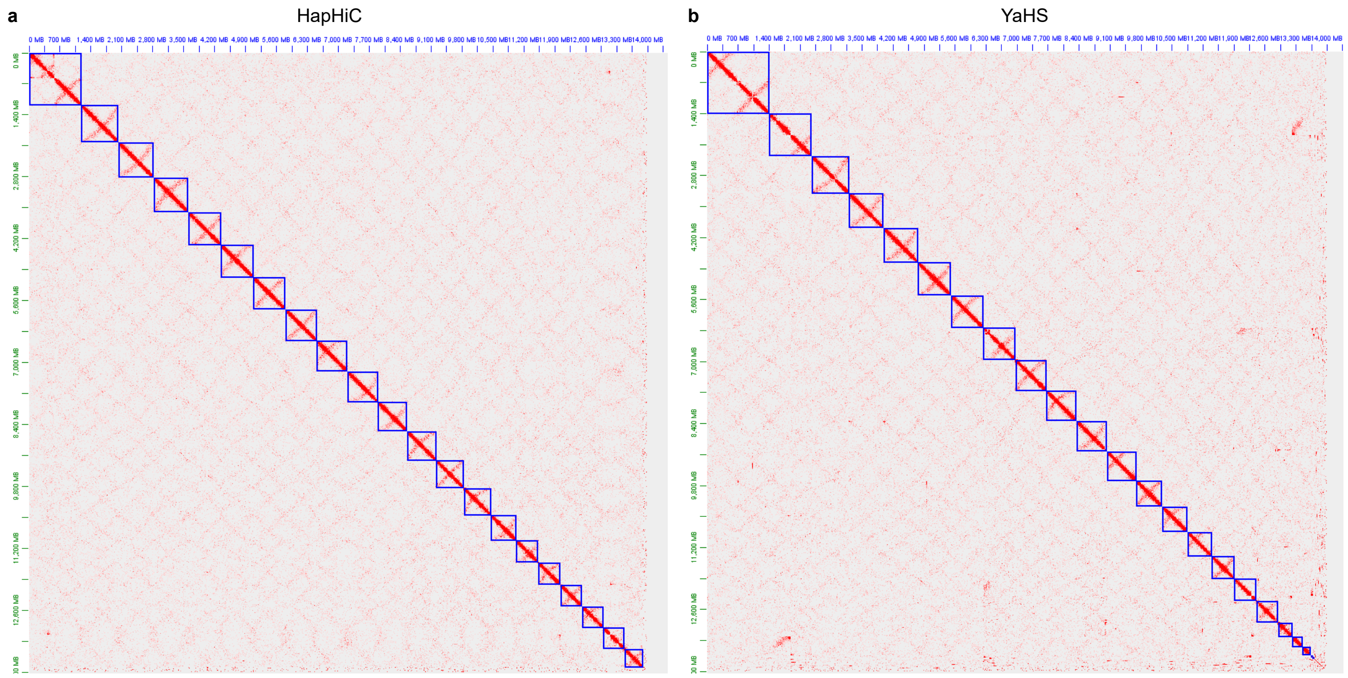


**Supplementary Fig. 58 |** Hi-C contact maps of HapHiC (**a**) and YaHS (**b**) scaffolds of the *Triticum aestivum* genome.

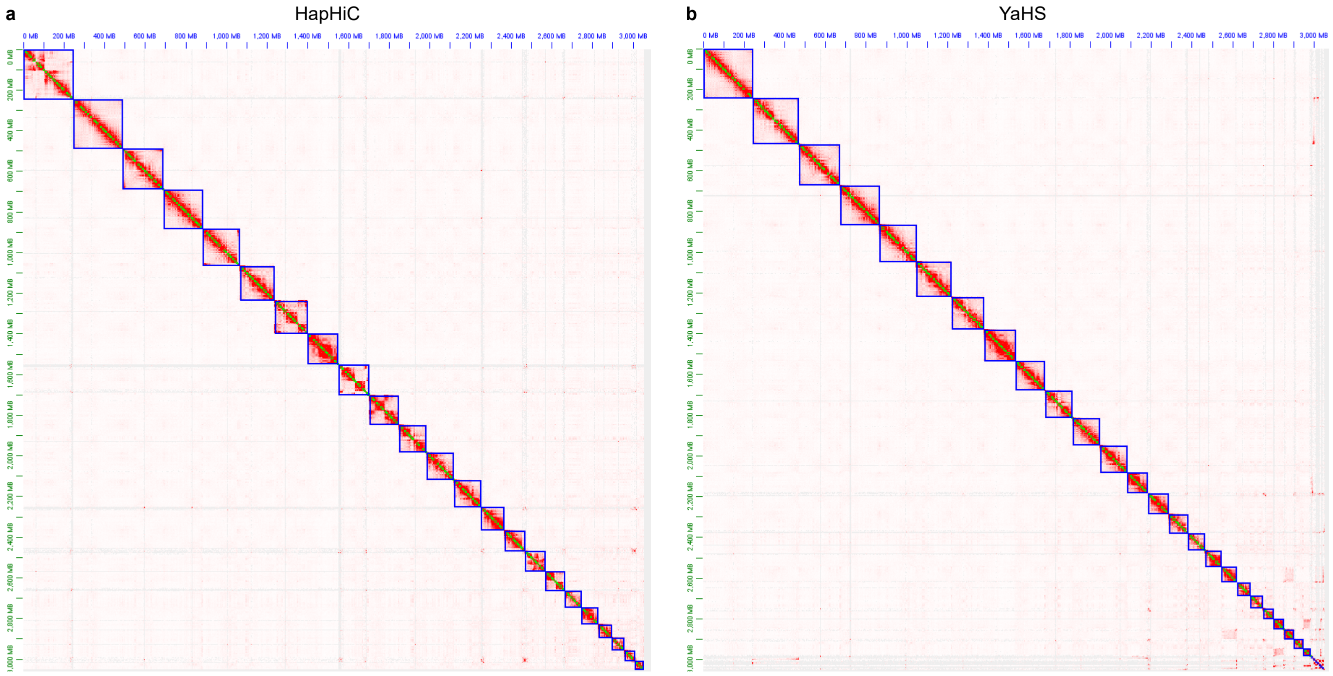


**Supplementary Fig. 59 |** Hi-C contact maps of HapHiC (**a**) and YaHS (**b**) scaffolds of the human CHM13v2.0_noY genome.

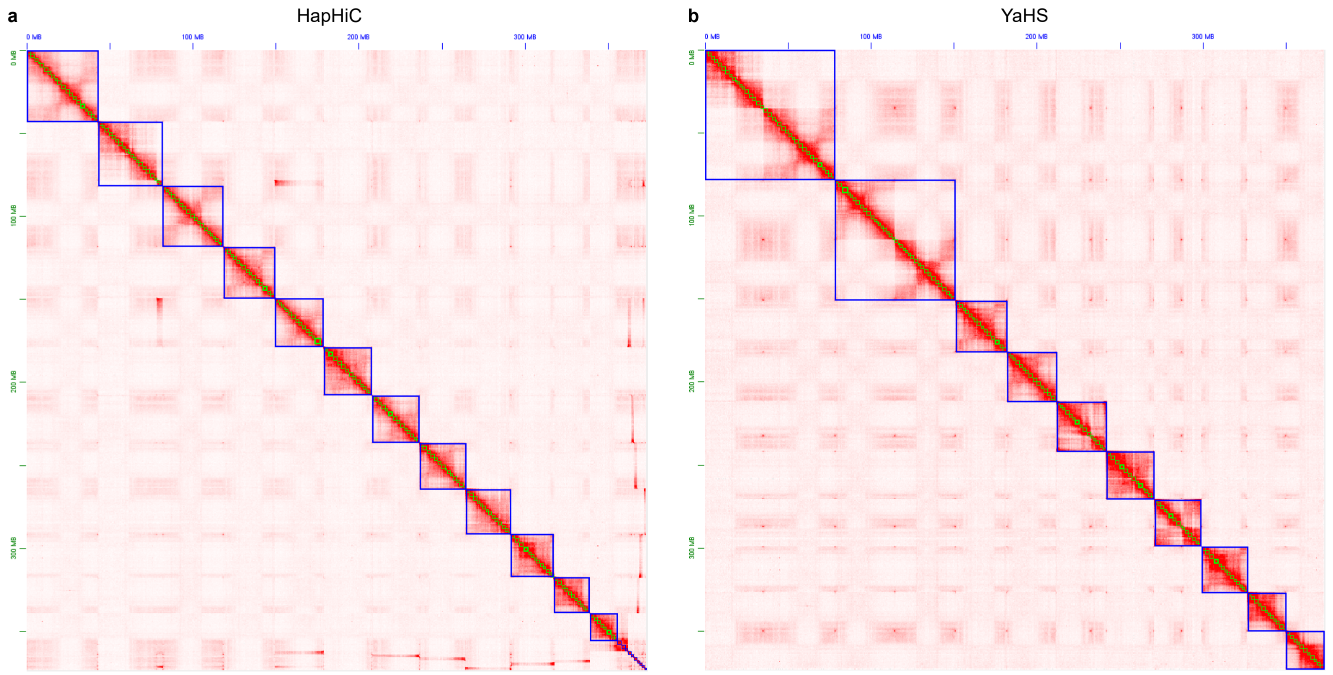


**Supplementary Fig. 60 |** Hi-C contact maps of HapHiC (**a**) and YaHS (**b**) scaffolds of the rice IRGSP-1.0 genome.

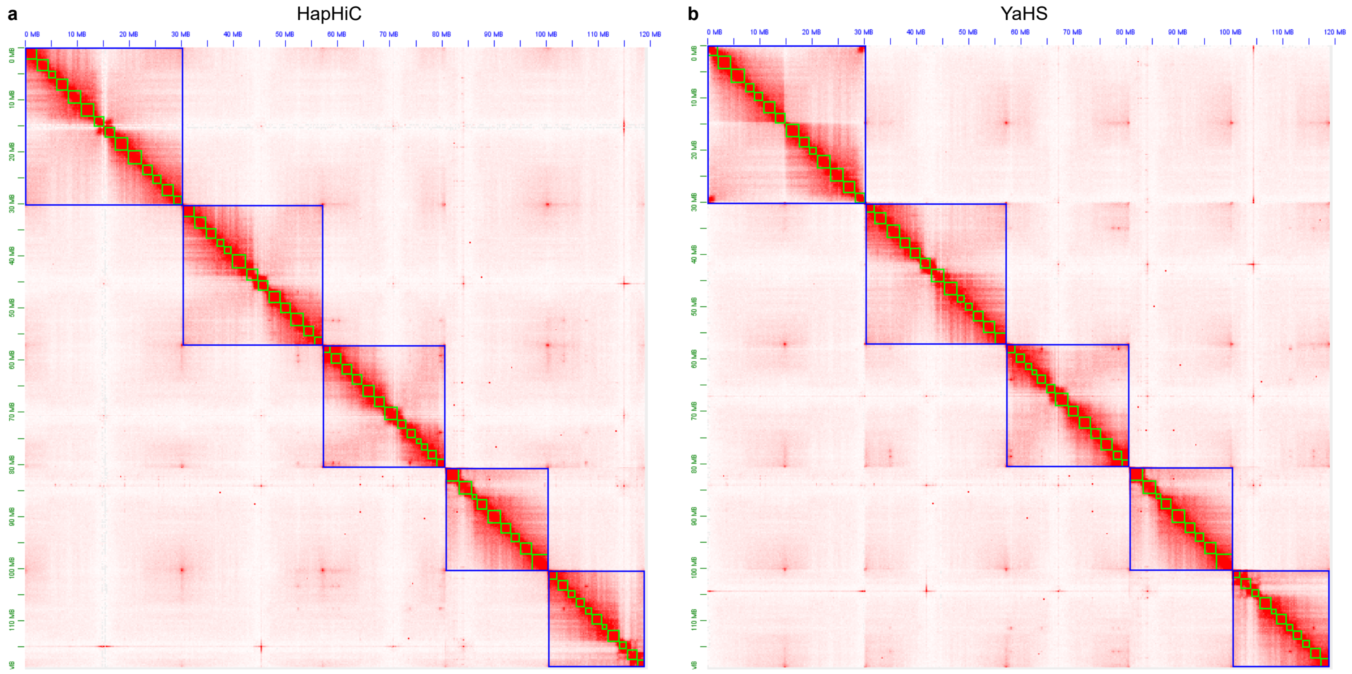


**Supplementary Fig. 61 |** Hi-C contact maps of HapHiC (**a**) and YaHS (**b**) scaffolds of the *Arabidopsis* TAIR10.1 genome.

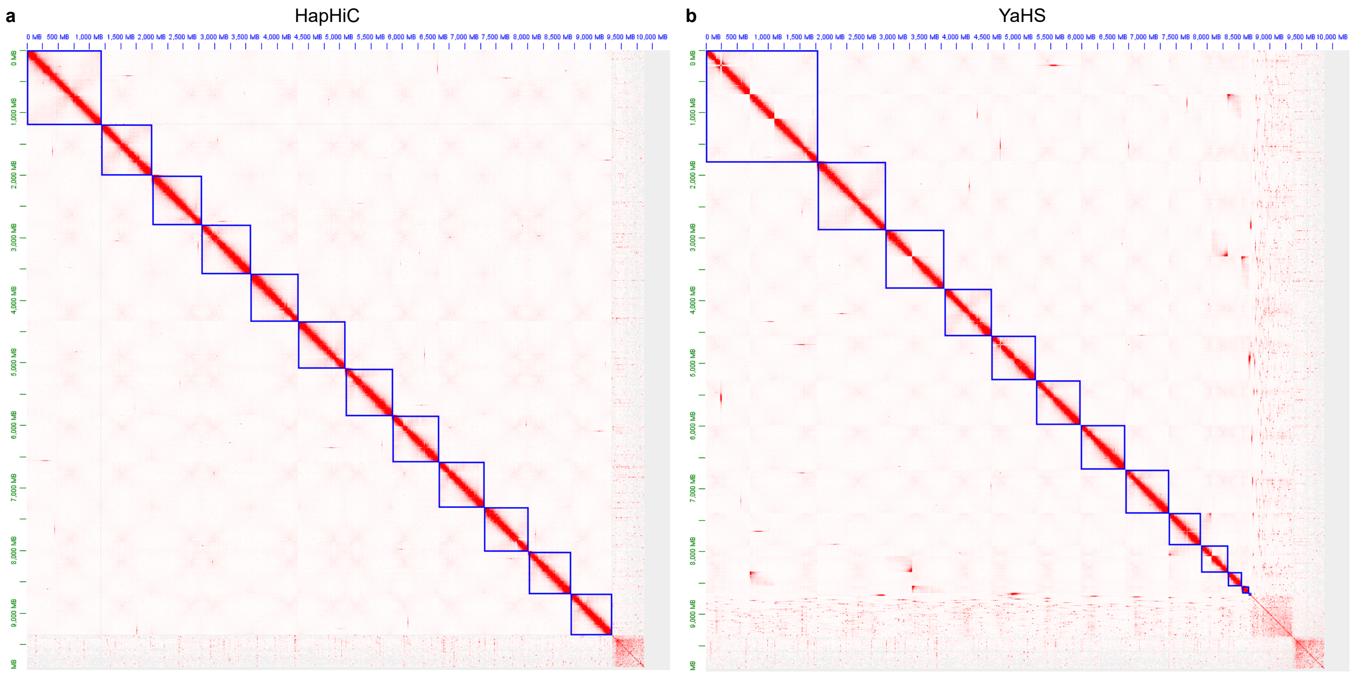


**Supplementary Fig. 62 |** Hi-C contact maps of HapHiC (**a**) and YaHS (**b**) scaffolds of the *Ginkgo biloba* genome.

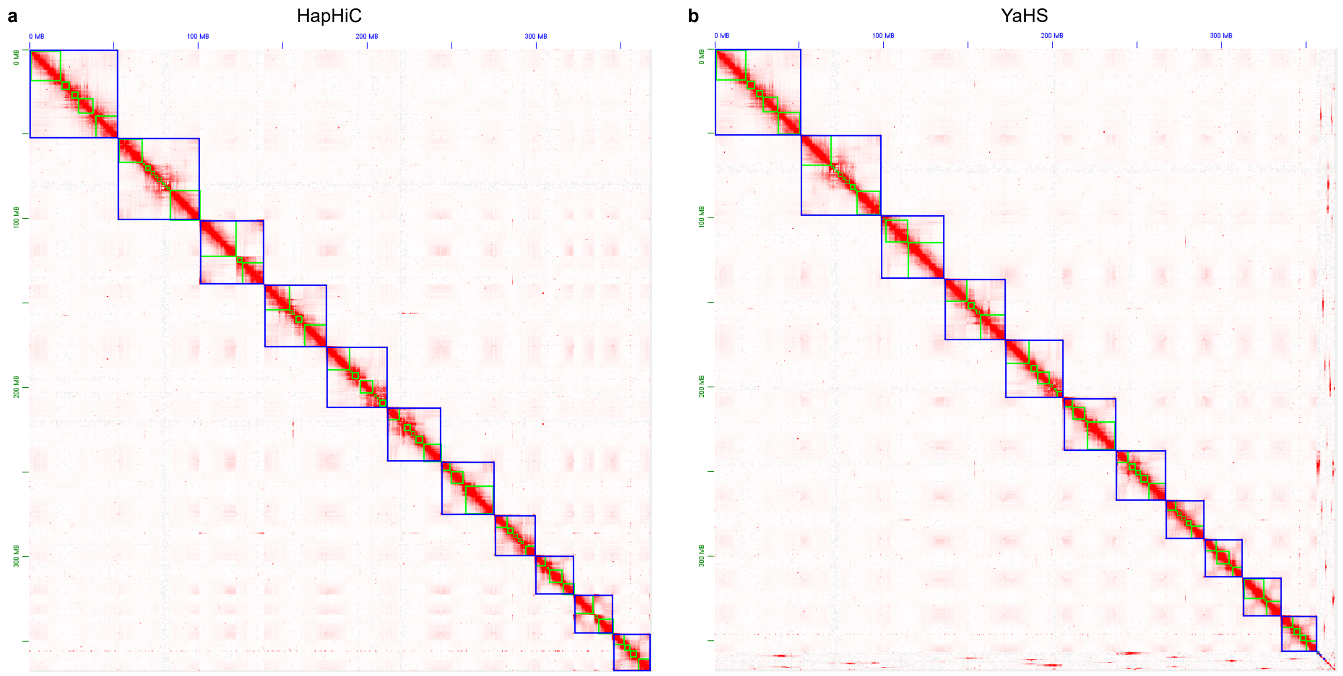


**Supplementary Fig. 63 |** Hi-C contact maps of HapHiC (**a**) and YaHS (**b**) scaffolds of the *Corylus mandshurica* genome.

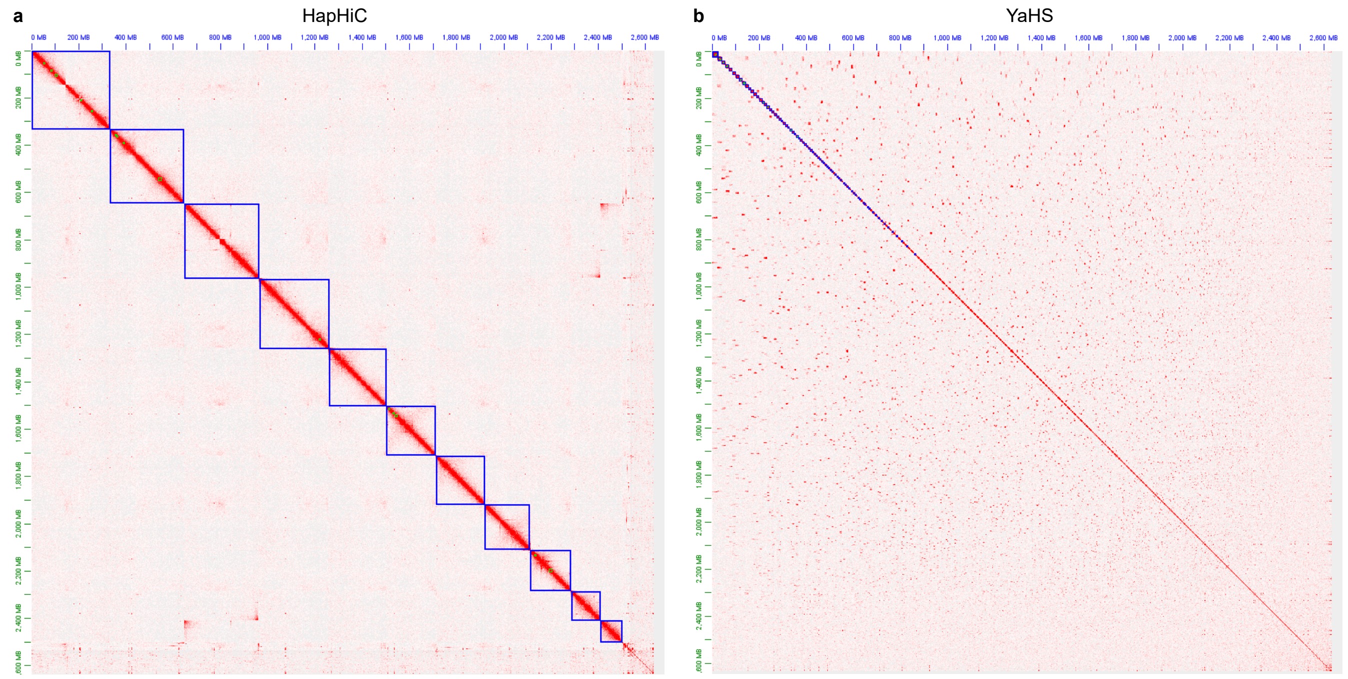


**Supplementary Fig. 64 |** Hi-C contact maps of HapHiC (**a**) and YaHS (**b**) scaffolds of the *Papaver somniferum* genome.

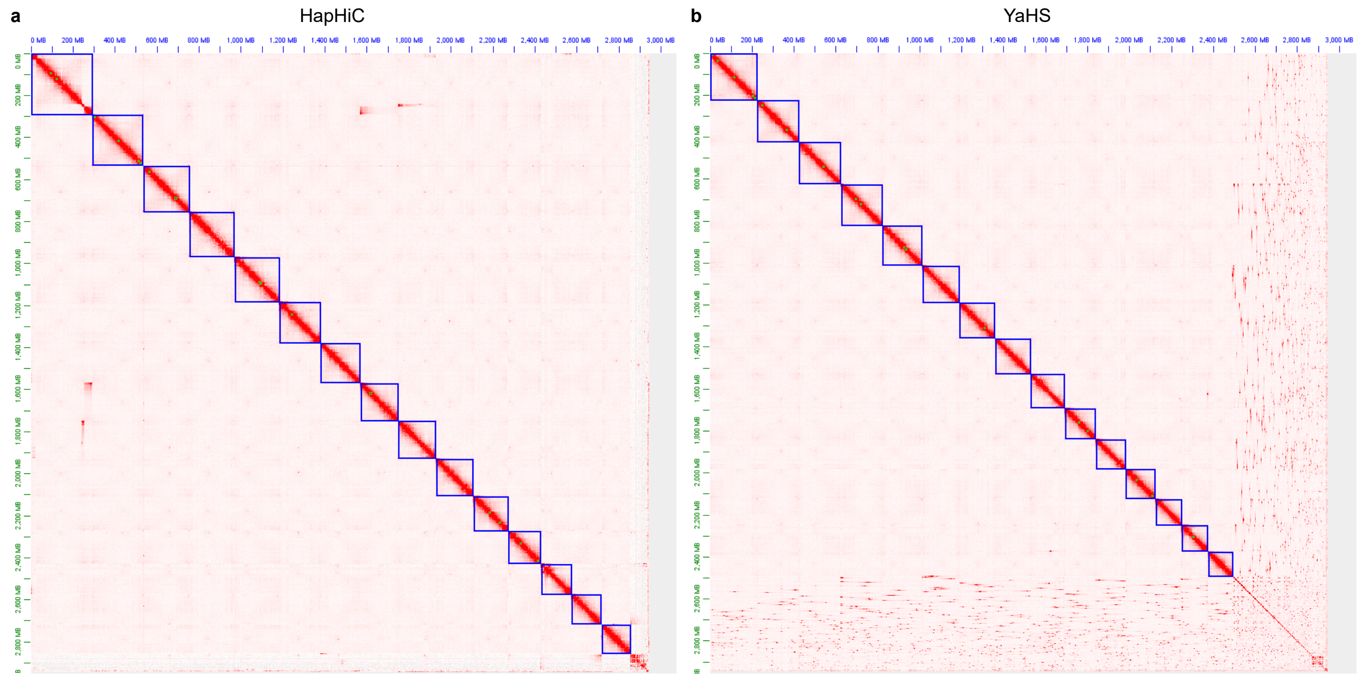


**Supplementary Fig. 65 |** Hi-C contact maps of HapHiC (**a**) and YaHS (**b**) scaffolds of the *Camellia sinensis* Huangdan genome.

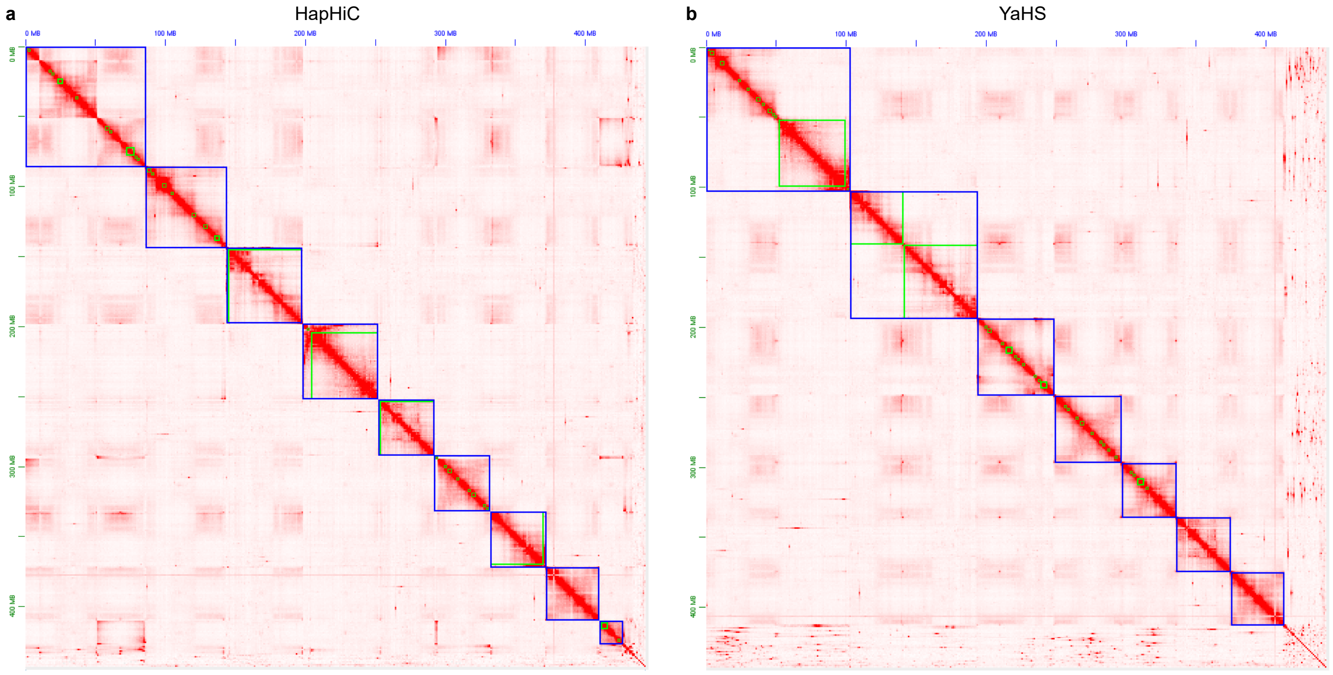


**Supplementary Fig. 66 |** Hi-C contact maps of HapHiC (**a**) and YaHS (**b**) scaffolds of the *Echinochloa haploclada* genome.

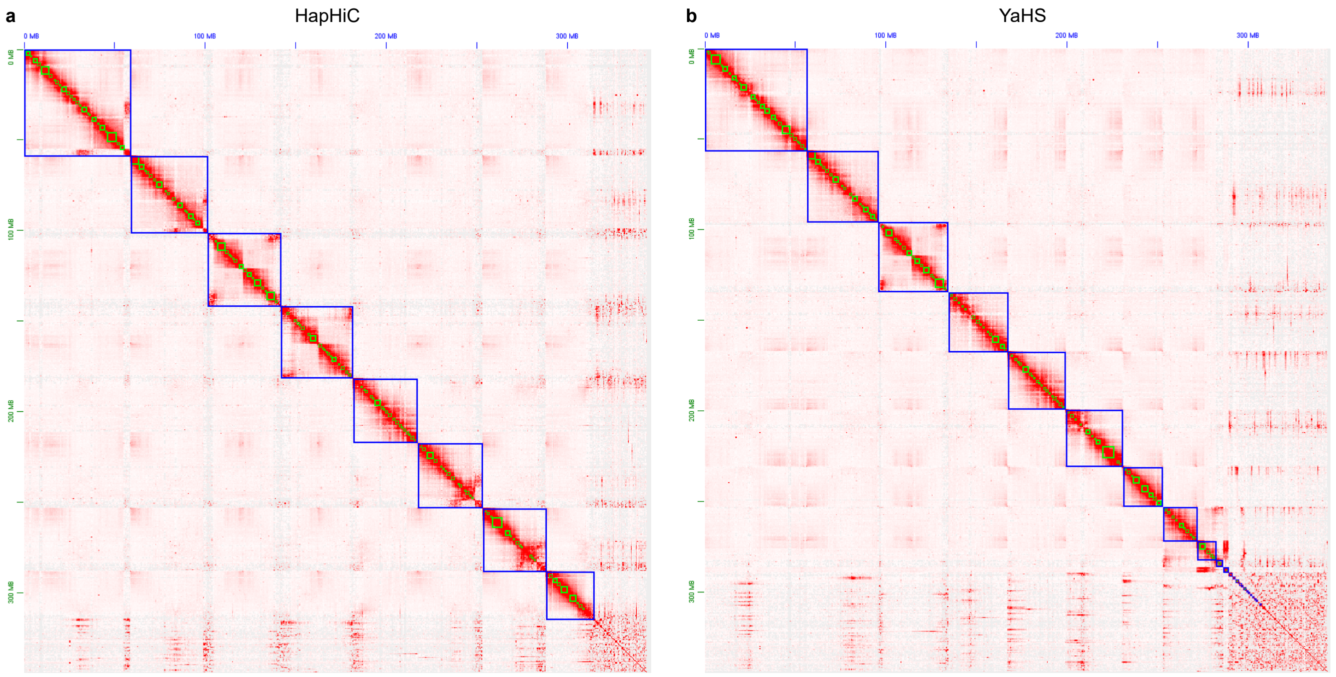


**Supplementary Fig. 67 |** Hi-C contact maps of HapHiC (**a**) and YaHS (**b**) scaffolds of the *Prunus avium* genome.

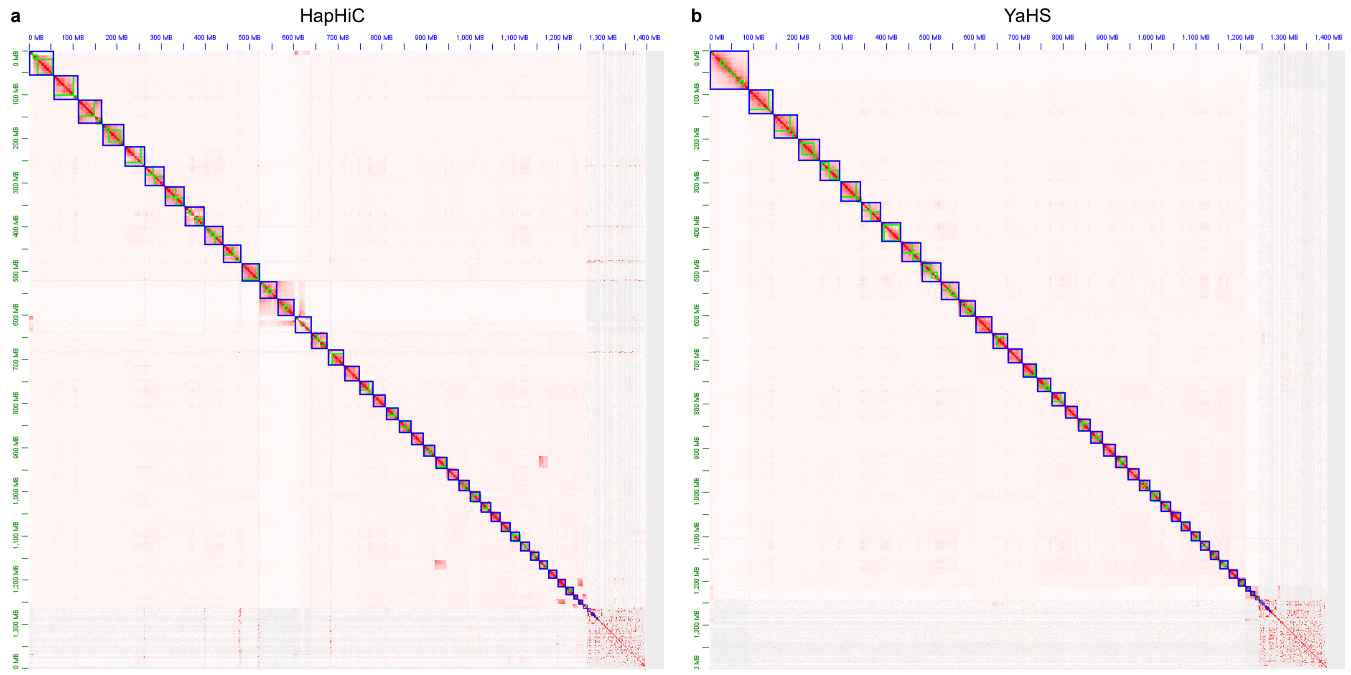


**Supplementary Fig. 68 |** Hi-C contact maps of HapHiC (**a**) and YaHS (**b**) scaffolds of the *Accipiter gentilis* genome.

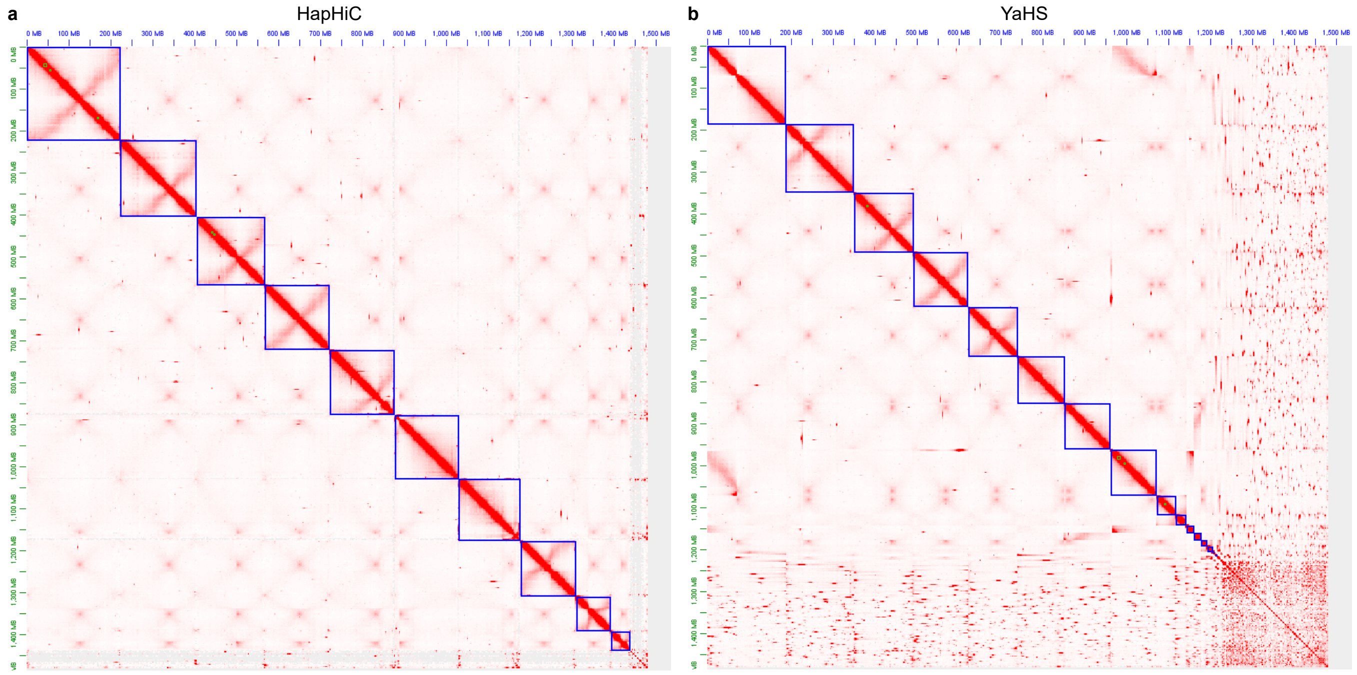


**Supplementary Fig. 69 |** Hi-C contact maps of HapHiC (**a**) and YaHS (**b**) scaffolds of the *Xenopus tropicalis* genome.

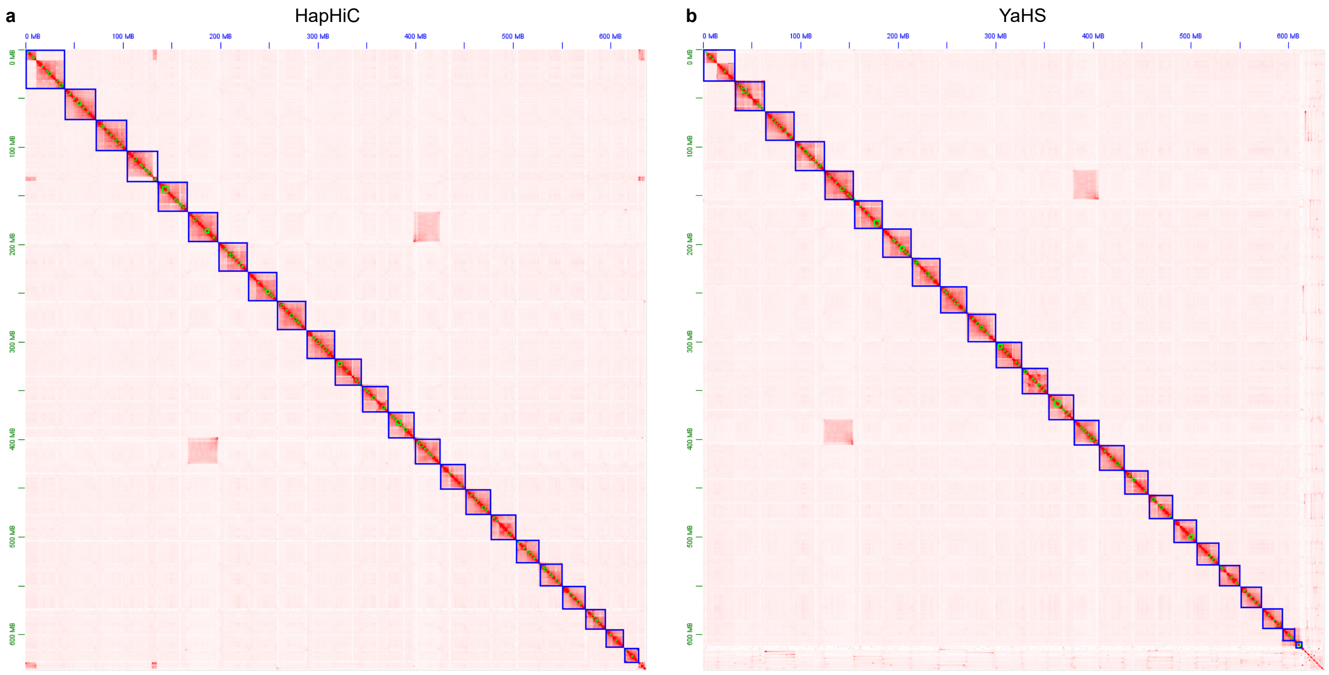


**Supplementary Fig. 70 |** Hi-C contact maps of HapHiC (**a**) and YaHS (**b**) scaffolds of the *Symphodus melops* genome.

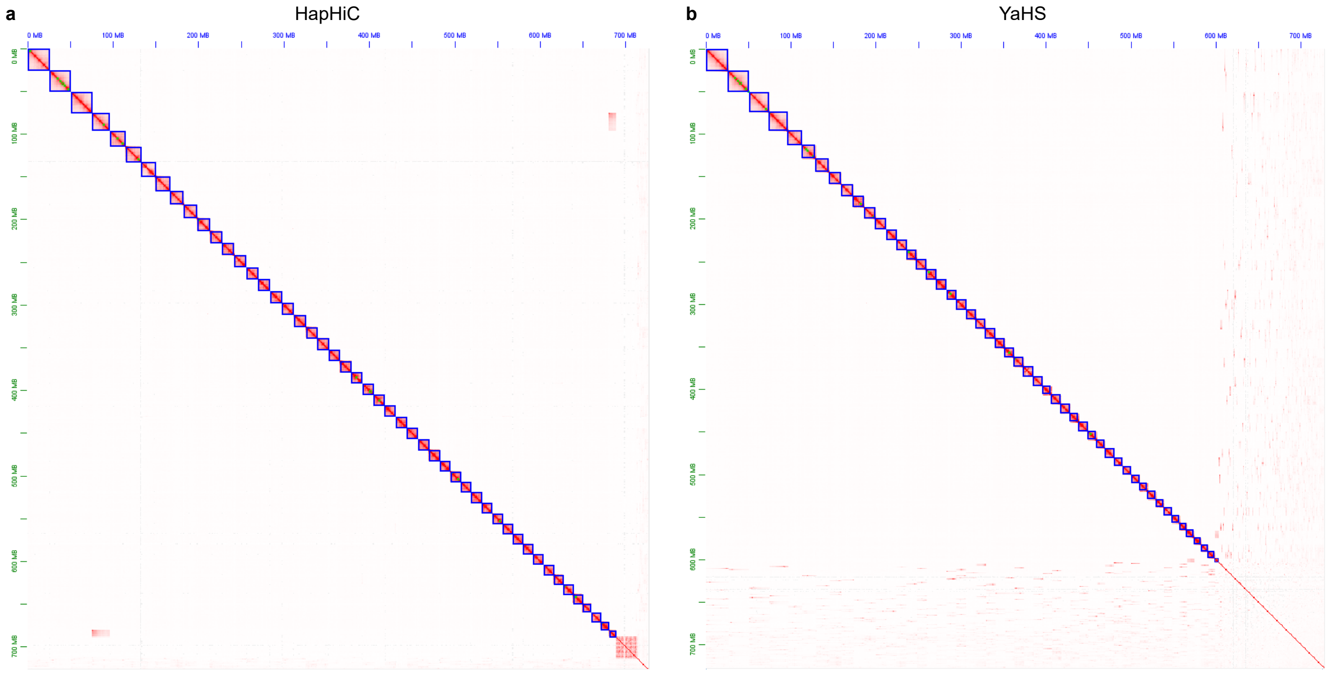


**Supplementary Fig. 71 |** Hi-C contact maps of HapHiC (**a**) and YaHS (**b**) scaffolds of the *Antheraea pernyi* genome.

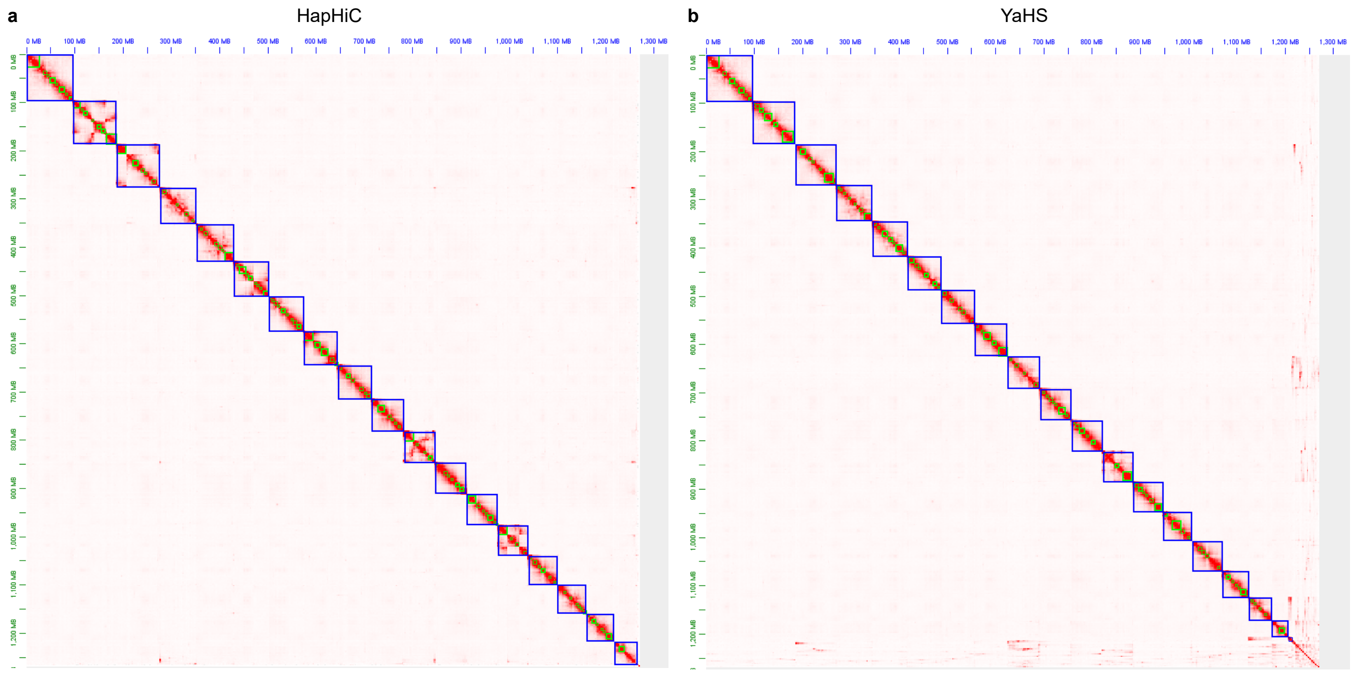


**Supplementary Fig. 72 |** Hi-C contact maps of HapHiC (**a**) and YaHS (**b**) scaffolds of the *Sterомphala cineraria* genome.

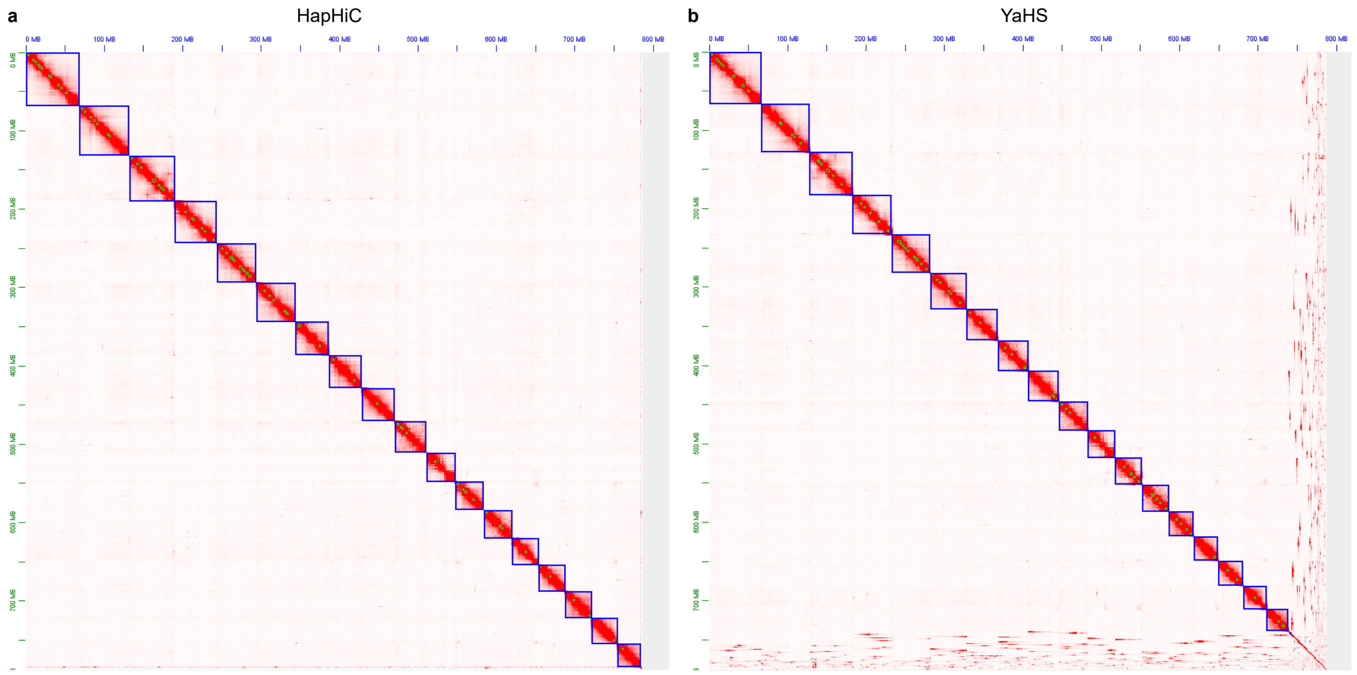


**Supplementary Fig. 73 |** Hi-C contact maps of HapHiC (**a**) and YaHS (**b**) scaffolds of the *Lumbricus rubellus* genome.

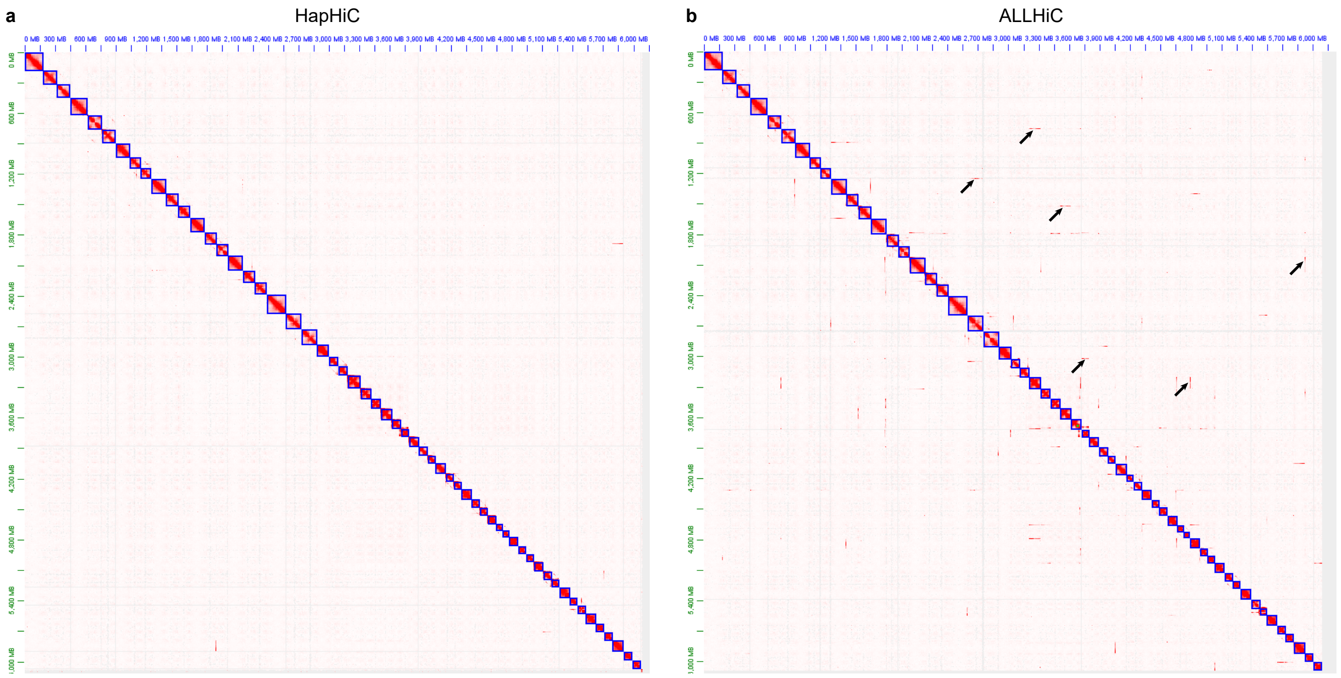


**Supplementary Fig. 74 |** Hi-C contact maps of HapHiC (**a**) and YaHS (**b**) scaffolds of the *M.* × *giganteus* genome. Black arrows indicate examples of misassignments between non-homologous chromosomes in the ALLHiC scaffolds.

#### Supplementary Data

**Supplementary Data 1-6** are provided online as separated excel files. Titles for each file and sheet are listed as follows:

**Supplementary Data 1:** Performance evaluation of Hi-C-based scaffolding tools in chromosome assignment under various adverse factors **(11 sheets in total)**

- **Sheet 1:** Performance evaluation of Hi-C-based scaffolding tools in chromosome assignment on assemblies with varying contig N50 values
- **Sheet 2:** Performance evaluation of Hi-C-based scaffolding tools in chromosome assignment on assemblies with varying coefficients of variation of contig length
- **Sheet 3:** Performance evaluation of Hi-C-based scaffolding tools in chromosome assignment with varying effective Hi-C sequencing depths
- **Sheet 4:** Performance evaluation of Hi-C-based scaffolding tools in chromosome assignment on assemblies with varying proportions of chimeric contigs
- **Sheet 5:** Performance evaluation of ALLHiC with and without assembly correction in chromosome assignment on assemblies with varying proportions of chimeric contigs
- **Sheet 6:** Performance evaluation of Hi-C-based scaffolding tools in chromosome assignment on assemblies with varying proportions of collapsed contigs
- **Sheet 7:** Performance evaluation of Hi-C-based scaffolding tools in chromosome assignment on assemblies with varying sequence divergence levels
- **Sheet 8:** Performance evaluation of Hi-C-based scaffolding tools in chromosome assignment on assemblies with varying switch error rates
- **Sheet 9:** Performance evaluation of Hi-C-based scaffolding tools in chromosome assignment on assemblies with identical switch error rate (20%) but varying contig N50 values
- **Sheet 10:** Performance evaluation of Hi-C-based scaffolding tools in chromosome assignment on assemblies with varying ploidies
- **Sheet 11:** Performance evaluation of Hi-C-based scaffolding tools in chromosome assignment on assemblies with varying ploidies and 5% each of chimeric, collapsed, and switch errors

**Supplementary Data 2:** Performance evaluation of Hi-C-based scaffolding tools in contig ordering and orientation on assemblies with varying contig N50 values **(two sheets in total)**

- **Sheet 1:** Performance evaluation of HapHiC and ALLHiC in contig ordering and orientation using the built-in scoring system of ALLHiC
- **Sheet 2:** Performance evaluation of all Hi-C-based scaffolding tools in contig ordering and orientation using Lin's concordance correlation coefficient and DERANGE cost

**Supplementary Data 3:** Time and memory usage of Hi-C-based scaffolding tools **(five sheets in total)**

- **Sheet 1:** Time and memory usage of HapHiC and ALLHiC in contig ordering and orientation
- **Sheet 2:** Time and memory usage of the entire pipeline for each Hi-C-based scaffolding tool
- **Sheet 3:** Time and memory usage of each Hi-C-based scaffolding tool in assembly correction
- **Sheet 4:** Time and memory usage of each Hi-C-based scaffolding tool in processing Hi-C data at varying depths
- **Sheet 5:** Time and memory usage of HapHiC, ALLHiC, and YaHS pipelines in scaffolding published assemblies

**Supplementary Data 4:** Details of published genome assemblies and Hi-C data used for HapHiC validation **(one sheet in total)**

**Supplementary Data 5:** Comparison of assembly correction results between HapHiC and ALLHiC on manually annotated contigs in the first haplotype of *S. spontaneum* Np-X chromosome 1 **(one sheet in total)**

**Supplementary Data 6:** Correlation analysis of genetic maps with the A subgenome of *M.* × *giganteus* (MgiA) and the genome of *M. sinensis* (MsiA) **(one sheet in total)**
